# Plant autophagosomes mature into amphisomes prior to their delivery to the central vacuole

**DOI:** 10.1101/2022.02.26.482093

**Authors:** Jierui Zhao, Mai Thu Bui, Juncai Ma, Fabian Künzl, Juan Carlos De La Concepcion, Yixuan Chen, Sofia Petsangouraki, Azadeh Mohseni, Marta Garcia Leon, Marta Salas Gomez, Caterina Giannini, Dubois Gwennogan, Roksolana Kobylinska, Marion Clavel, Swen Schellmann, Yvon Jaillais, Jiri Friml, Byung-Ho Kang, Yasin Dagdas

**Affiliations:** Gregor Mendel Institute (GMI), Austrian Academy of Sciences, Vienna BioCenter (VBC), Vienna, Austria; Vienna BioCenter PhD Program, Doctoral School of the University at Vienna and Medical University of Vienna, Vienna, Austria.; School of Life Sciences, Centre for Cell & Developmental Biology and State Key Laboratory of Agrobiotechnology, The Chinese University of Hong Kong, Shatin, New Territories, Hong Kong, China; Institute of Science and Technology (IST) Austria, 3400, Klosterneuburg, Austria; Laboratoire Reproduction et Développement des Plantes (RDP), Université de Lyon, ENS de Lyon, CNRS, INRAE, F-69342 Lyon, France; Botanik III, Biocenter, Universtiy of Cologne, Zülpicher Str. 47B, 50674, Cologne, Germany.

## Abstract

Autophagosomes are double-membraned vesicles that traffic harmful or unwanted cellular macromolecules to the vacuole for recycling. Although autophagosome biogenesis has been extensively studied, mechanisms of autophagosome maturation, i.e., delivery and fusion with the vacuole, remain largely unknown in plants. Here, we have identified an autophagy adaptor, CFS1, that directly interacts with the autophagosome marker ATG8 and localizes on both membranes of the autophagosome. Autophagosomes form normally in *Arabidopsis thaliana cfs1* mutants, but their delivery to the vacuole is disrupted. CFS1’s function is evolutionarily conserved in plants as it also localizes to the autophagosomes and plays a role in autophagic flux in the liverwort *Marchantia polymorpha*. CFS1 regulates autophagic flux by connecting autophagosomes with the ESCRT-I component VPS23, leading to the formation of amphisomes. Disrupting the VPS23-CFS1 interaction affects autophagic flux and renders plants sensitive to starvation stress. Altogether, our results reveal a deeply conserved mechanism of vacuolar delivery in plants that is mediated by amphisomes.

## Introduction

Macroautophagy (hereafter autophagy) is a conserved vacuolar trafficking pathway that mediates three **R**s in eukaryotic cells, including plants: (1) It **R**emodels the cellular environment for developmental and temporary reprogramming events that underlie cellular differentiation and adaptation. In plants for example, autophagy is essential for callus regeneration in *Arabidopsis thaliana*, wound-induced de-differentiation in *Physcomitrium patens*, sperm maturation in *Marchantia polymorpha*, and pollen formation in rice (Rodriguez et al., 2020; Norizuki et al., 2021; Kurusu et al., 2014). Autophagy-mediated cellular adaptation is also crucial for stress tolerance including drought, infection, and high temperature stress. Studies involving Arabidopsis, maize, and rice have shown that autophagy mutants such as *atg2*, *atg5*, and *atg7* are highly susceptible to biotic and abiotic stress factors and undergo early senescence (Signorelli et al., 2019; McLoughlin et al., 2018; Wada et al., 2015). (2) At the cellular level, autophagy **R**enovates the cell by removing the organelles, protein complexes, and other dysfunctional macromolecules that would otherwise reduce cellular fitness (Dikic, 2017; Marshall and Vierstra, 2018). Finally, (3) during nutrient limitation, autophagy **R**eplenishes cellular energy pools and prolongs survival by recycling surplus cellular material (McLoughlin et al., 2020; Rabinowitz and White, 2010). Thus, autophagy is a major degradation and recycling pathway that keeps the cell in tune with the ever-changing environment and maintains cellular homeostasis.

The main vehicle of autophagy is a *de novo* formed, double membrane vesicle termed the autophagosome. Autophagosomes capture their cargo and deliver them to the vacuole (or lysosomes in metazoans) for recycling. Autophagosome biogenesis involves the concerted action of highly conserved ATG (autophagy-related gene) proteins that coordinate the nucleation and growth of a cup shaped phagophore around the autophagic cargo (Nakatogawa, 2020; Chang et al., 2021; Weidberg et al., 2011). The two opposing membranes are then sealed with ESCRT (endosomal sorting complex required for transport) proteins to form the autophagosome (Chang et al., 2021). Both autophagosome membranes are labelled with lipidated ATG8 family proteins that interact with (i) other ATG proteins to coordinate autophagosome formation, (ii) cargo receptors that selectively recruit cargo macromolecules and underlie selective autophagy, and (iii) adaptor proteins that mediate the trafficking and vacuolar fusion of autophagosomes (Stolz et al., 2014). Most of these ATG8-interacting proteins contain highly conserved short linear motifs termed the ATG8 interacting motif (AIM). The core AIM is denoted as [W/F/Y]xx[L/I/V], where x represents any amino acid. The AIM peptide is bound by the highly conserved W and L loops, collectively known as the AIM-docking site (ADS), on ATG8 (Birgisdottir et al., 2013). Despite recent advances in cargo receptor identification and characterization, no autophagy adaptors have been characterized in plants. As a result, we have only a limited understanding of how autophagosome are delivered to the vacuole in plants.

Autophagosome maturation is logistically different in yeast, metazoans, and plants. In yeast, autophagosome biogenesis and maturation happens in the vicinity of the vacuole; the coordination of these processes is aided by spatial proximity (Zhao and Zhang, 2019). In metazoans, autophagosomes are formed at various sites around the cell and subsequently fuse with endosomes or lysosomes (Zhao et al., 2021). Autophagosome-endosome fusions create amphisomes, which mature into autolysosomes by acquiring lytic enzymes (Sanchez-Wandelmer and Reggiori, 2013). Despite these logistical differences, in both yeast and metazoans the concerted action of dedicated SNARE (soluble N-ethylmaleimide-sensitive factor attachment protein receptor) proteins, tethering factors, and adaptors mediate the fusion of autophagosomes with the lytic compartments (Zhao et al., 2021). In plants, autophagosomes are formed at sites around the cell and are then delivered to the central vacuole, which can occupy as much as 80% of the cell volume (Marshall and Vierstra, 2018). The molecular details of autophagosome trafficking, fusion with the vacuole, and how these events are coordinated with other vacuolar trafficking pathways are currently unknown in plants. Whether or not plant autophagosomes converge into amphisomes before arriving to the central vacuole also remains unknown.

To address these questions, we focused on the identification of autophagy adaptors. We developed a differential centrifugation protocol to enrich for intact autophagosomes prior to affinity purification-mass spectrometry (AP-MS) with ATG8 as bait. This approach identified CFS1 (<u>c</u>ell death related endosomal <u>F</u>YVE/<u>S</u>YLF protein 1), a highly conserved FYVE (Fab-1, YGL023, Vps27, and EEA1) and SYLF (SH3YL1, Ysc84p/Lsb4p, Lsb3p, and plant FYVE) domain-containing protein that was previously linked to autophagy (Sutipatanasomboon et al., 2017; Kim et al., 2022). Characterization of CFS1 revealed that it interacts with ATG8 in an AIM-dependent manner and specifically regulates autophagic flux in both *Arabidopsis thaliana* and *Marchantia polymorpha*. Genome wide yeast two hybrid screening showed that CFS1 also interacts with the multivesicular body-localized ESCRT-I complex protein VPS23. The CFS1-VPS23 interaction is crucial for autophagic flux. Live cell imaging and electron microscopy analyses demonstrate that CFS1 colocalizes with VPS23 at amphisomes. Altogether, we define a new, prevacuolar sorting hub for the coordination of vacuolar trafficking pathways in plants.

## Results

### Differential centrifugation coupled to affinity purification-mass spectrometry (AP-MS) revealed autophagosome associated proteins in *Arabidopsis thaliana*

Since autophagy adaptors play crucial roles in autophagic flux but are unknown in plants, we first set out to identify autophagy adaptors. We induced autophagy in GFP-ATG8A expressing *A. thaliana* seedlings via Torin treatment and performed differential centrifugation experiments to enrich for small membranous compartments, including intact autophagosomes, while removing larger, bulkier compartments such as organelles (Liu and Bassham, 2010; LaMontagne et al., 2016) (Fig. 1A). To verify that the membrane-associated fractions (P4) contained intact autophagosomes, we performed protease protection assays. Both NBR1, a well characterized autophagy receptor that is localized within the autophagosomes, and ATG8, which localizes on both sides of the autophagosome, were protected in these assays (Svenning et al., 2011; Stolz et al., 2014) (Fig. S1A-B). After addition of Triton X-100, a detergent that destabilizes membranes, both proteins became sensitive to protease treatment (Fig. S1B). These experiments demonstrate the enrichment of intact autophagosomes (Borner et al., 2005). We combined this approach with AP-MS and screened for protease sensitive proteins (i.e., those localized on the outer autophagosome membrane) in the membrane enriched fractions (Fig. 1A). Mass spectrometry analysis revealed 48 proteins that are ATG8E-associated, P4-enriched, and proteinase K-sensitive, including SH3P2, FRA3, NUP93A, and NUP98A (Fig. 1B-C, Table S1-2). One of these 48 proteins was the FYVE and SYLF domain-containing protein CFS1 (At3g43230) (Fig. 1B-C). Since CFS1 has previously been linked to autophagy and FYVE and SYLF domain-containing proteins are well-known players in vesicle trafficking (Sutipatanasomboon et al., 2017; Kim et al., 2022; Melia et al., 2020), we decided to characterize CFS1 in depth.

**Figure 1.**
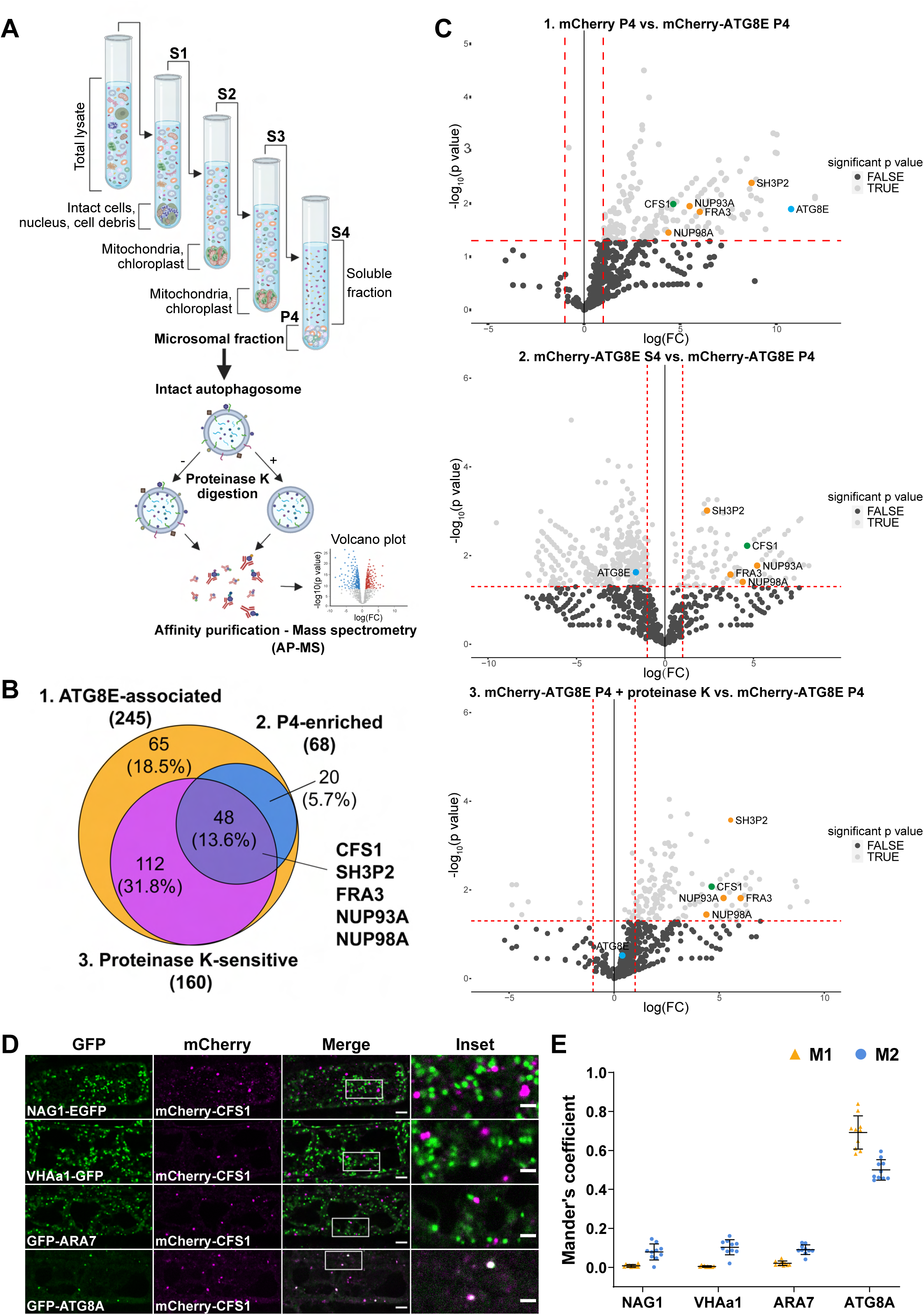
Differential centrifugation coupled to affinity purification-mass spectrometry (AP-MS) revealed autophagosome associated proteins in *Arabidopsis thaliana*. **(A)** Schematic diagram showing the differential centrifugation coupled to affinity purification-mass spectrometry (AP-MS) workflow. Total lysate of Arabidopsis seedlings underwent several differential centrifugation steps where each time the supernatant (S) was transferred and the pellet (P) was left untouched. Samples were spun for 10 min at 1,000 g (S1), to remove intact cells, nuclei and cell debris; 10 min at 10,000g, to remove bigger organelles like mitochondria and chloroplasts (S2); 10 min at 15,000g, to further remove organelles (S3) and finally 60 min at 100,000 g (S4 and P4) (LaMontagne et al., 2016). The P4 fraction, containing small vesicles and microsomes (microsomal fraction), i.e., autophagosomes, was further subjected to 30 ng/μl proteinase K treatment (P4 + proteinase K). All S4, P4 and P4 + proteinase K samples were processed for AP-MS. **(B)** Venn diagram showing the overlap between ATG8E-associated, P4-enriched and protease K-sensitive proteins which were identified by the AP-MS described in (A). Seven-day-old Arabidopsis seedlings expressing pUBQ::mCherry or pUBQ::mCherry-ATG8E were treated with 3 μM Torin for 90 min for autophagy induction before lysing. Diagram was generated using Venny 2.1.0 (Oliveros, 2016). **(C)** Volcano plots of mCherry-ATG8E AP-MS datasets identified CFS1 in all three pairwise comparisons. Upper panel, volcano plot of the pairwise comparison of “mCherry P4” and “mCherry-ATG8E P4” shows proteins enriched by the bait mCherry-ATG8E. X- and Y-axis displays log fold-change (log(FC)) and - log10(p value), respectively. Dashed lines represent threshold for log(FC)>1 and p value<0.05. Only proteins passing both p value<0.05 and log(FC)<0 filter were considered in the first set “ATG8E- associated” in (B). Middle panel, volcano plot of the pairwise comparison of “mCherry-ATG8E S4” and “mCherry-ATG8E P4” shows proteins enriched by mCherry-ATG8E in the pellet (P4) relative to the supernatant (S4). X- and Y-axis displays log(FC) and -log10(p value), respectively. Dashed lines represent threshold for log(FC)>1 and p value<0.05. Only proteins passing both p value<0.05 and log(FC)<0 filter were considered in the second set “P4-enriched” in (B). Lower panel, volcano plot of the pairwise comparison of “mCherry-ATG8E P4 + proteinase K” and “mCherry-ATG8E P4” shows proteins enriched by mCherry-ATG8E in the pellet (P4) before proteinase K treatment. X- and Y-axis displays log(FC) and -log10(p value), respectively. Dashed lines represent threshold for log(FC)>1 and p value<0.05. Only proteins passing both p value<0.05 and log(FC)<0 filter were considered in the third set “Proteinase K-sensitive” in (B). For all volcano plots, CFS1, interested proteins, and ATG8E are labeled by green, orange or light blue dots, respectively. **(D)** Confocal microscopy images of Arabidopsis root epidermal cells co-expressing pUBQ::mCherry-CFS1 with either Golgi body marker p35S::NAG1-EGFP, trans Golgi network marker pa1::VHAa1-GFP, MVB marker pRPS5a::GFP-ARA7 or autophagosome marker pUBQ::GFP-ATG8A under nitrogen starvation conditions. Five-day-old Arabidopsis seedlings were incubated in nitrogen-deficient 1/2 MS media for 4 h for autophagy induction prior to imaging. Area highlighted in the white-boxed region in the merge panel was further enlarged and presented in the inset panel. Scale bars, 5 μm. Inset scale bars, 2 μm. **(E)** Quantification of confocal experiments in (D) showing the Mander’s colocalization coefficients between mCherry-CFS1 and the GFP-fused marker proteins NAG1, VHAa1, ARA7 or ATG8A. M1, fraction of GFP-fused marker signal that overlaps with mCherry-CFS1 signal. M2, fraction of mCherry- CFS1 signal that overlaps with GFP-fused marker signal. Bars indicate the mean ± standard deviation of 10 biological replicates.

### CFS1 localizes to the autophagosomes

To map the cellular distribution of CFS1, we generated transgenic Arabidopsis lines that stably co-express mCherry-CFS1 with endomembrane compartment markers, including NAG1-EGFP (Golgi bodies), VHAa1-GFP (trans-Golgi network), GFP-ARA7 (late endosomes), and GFP-ATG8A (autophagosomes) (Geldner et al., 2009; Bassham, 2015). Live cell confocal imaging and quantification under control and two different autophagy inducing conditions (nitrogen starvation and salt stress) showed that mCherry-CFS1 puncta specifically localize to the autophagosomes (Thompson et al., 2005; Liu et al., 2009) (Fig. S2A, Fig. S3A, Fig. 1D). Mander’s colocalization coefficients showed that both the M1 and M2 between mCherry-CFS1 and GFP- ATG8A are higher than 0.4, while the M1 and M2 between mCherry-CFS1 and other GFP markers are less than 0.2 (Fig. 1E, Fig. S2B, Fig. S3B). We also performed microscopy experiments with the amphiphilic styryl dye FM4-64, a stain that is widely used for tracing endocytic vesicles (Rigal et al., 2015). CFS1 did not colocalize with FM4-64 even under autophagy inducing conditions (Fig. S4).

To further corroborate the autophagosome localization of CFS1, we visualized it with two other autophagosome localized proteins: ATG11, a core autophagy protein that is crucial for recruitment of selective autophagy receptors and cargoes to the autophagosome, and NBR1 (Zientara-Rytter and Subramani, 2020; Svenning et al., 2011). mCherry-CFS1 co-localized with both GFP-ATG11 and NBR1-GFP upon induction of autophagy with salt stress (Fig S5). Finally, we performed spinning disc time-lapse imaging of Arabidopsis lines that stably co-express mCherry-CFS1 with GFP-ATG8A or NBR1-GFP. Consistent with our confocal microscopy data, these time course experiments showed that CFS1 moves together with ATG8 and NBR1 puncta (Supplementary videos 1 and 2). Altogether, these results demonstrate that CFS1 specifically labels autophagic compartments.

A previous study analyzing FYVE domain-containing proteins in Arabidopsis identified another protein (CFS2, At1g29800) that also has FYVE and SYLF domains and shares 57.3% identity with CFS1 (Wywial and Singh, 2010) (Fig. S6A). Our phylogenetic analysis of CFS1 proteins across the plant kingdom detected no homology between CFS1 and CFS2 (Fig. S7). CFS1 and CFS2 formed separate, well-supported clades with different evolutionary histories. Nevertheless, we tested whether CFS2 plays a role in autophagy. We first carried out microscopy experiments similar to those described above. Although CFS2 was stably expressed, it did not form any puncta and had a diffuse localization pattern, even under autophagy inducing conditions (Fig. S6B-C). Nitrogen starvation plate assays, which are typically used to evaluate autophagy defects in Arabidopsis, showed no difference between *cfs2* mutants and wild type Col-0 plants (Phillips et al., 2008) (Fig. S6D). In contrast, *cfs1* mutants showed early senescence after 10-days of nitrogen starvation, similar to the autophagy deficient *atg5* mutant (Thompson et al., 2005) (Fig. S6D). We then compared autophagic flux in *cfs1* and *cfs2* mutants. NBR1 is degraded upon induction of autophagy, and this is used as a proxy for autophagic flux measurements (Bassham, 2015). Under both nitrogen starvation and salt stress conditions, *cfs1* mutants had higher NBR1 levels compared to wild-type plants (Fig. S6E-H). However, *cfs2* mutants showed no significant difference, and *cfs1cfs2* double mutants were comparable to *cfs1* (Fig. S6E-H). Altogether, these results suggest CFS2 is not involved in autophagy and prompted us to focus on CFS1 for further characterization.

### CFS1 interacts with ATG8 in an AIM dependent manner

Autophagy adaptors interact with ATG8 directly via an ATG8 interacting motif (AIM) (Birgisdottir et al., 2013; Stolz et al., 2014; Zaffagnini and Martens, 2016). To investigate whether CFS1 interacts with ATG8 in an AIM-dependent manner, we mixed crude extracts from *E. coli* expressing WT GST-ATG8A, an ADS (AIM-docking site) mutant of ATG8 (GST-ATG8A^ADS^), or GST alone with Arabidopsis lysates expressing mCherry-CFS1, and performed a GST pull downs. mCherry-CFS1 interacted with GST-ATG8A, but not with GST-ATG8A^ADS^ or GST alone (Fig. 2A). Consistently, the interaction between CFS1 and ATG8 could be outcompeted with a high affinity AIM peptide, but not with an AIM mutant peptide (Stephani et al., 2020) (Fig. 2A).We then set out to identify the ATG8 interacting motif in CFS1. Amino acid sequence alignments revealed a highly conserved candidate AIM between the FYVE and SYLF domains of CFS1 (Fig. S8). To test if this motif is important for ATG8 interaction, we mutated the core AIM residues into alanine (WLNL-267-ALNA) and expressed the resulting mCherry-CFS1^AIM^ in the *cfs1* mutant. This mutation did not affect CFS1 stability as both proteins accumulate to similar levels (Fig. 2B). Pull down experiments showed that, in contrast to mCherry-CFS1, mCherry-CFS1^AIM^ interacted significantly less with GST-ATG8A (Fig. 2B). We then performed *in vivo* co-immunoprecipitation experiments using plants stably co-expressing mCherry-CFS1 with GFP-ATG8A. We observed a strong association between CFS1 and ATG8A (Fig. 2C). This association was further strengthened during nitrogen starvation, suggesting recruitment of CFS1 to the autophagosomes upon autophagy induction (Fig. 2C). In contrast, mCherry-CFS1^AIM^-GFP-ATG8A association was substantially weaker under both control and nitrogen starved conditions (Fig. 2C).

**Figure 2.**
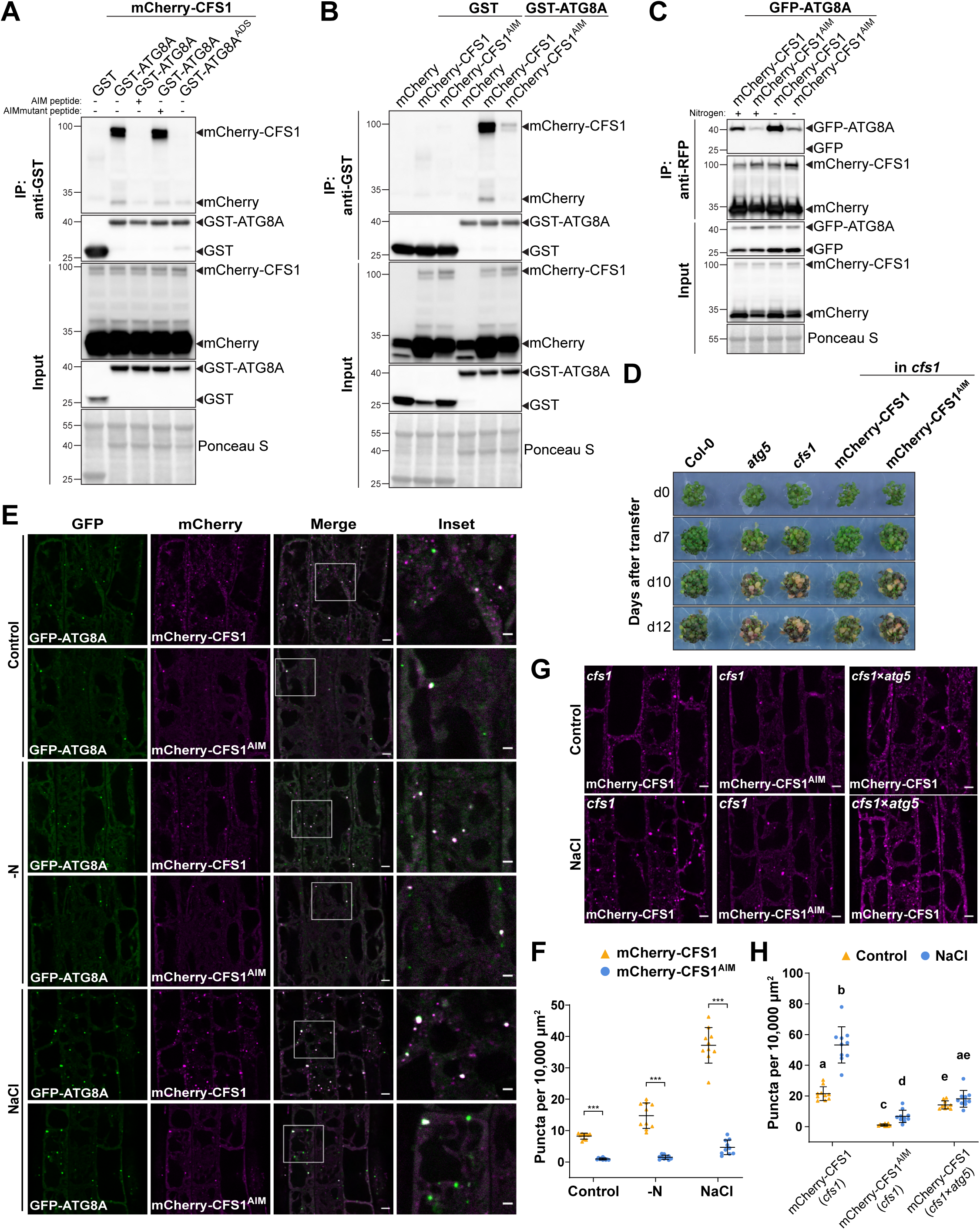
CFS1 interacts with ATG8 in an AIM dependent manner. **(A)** GST pull-down coupled with peptide competition with *E. coli* lysates containing GST, GST-ATG8A or GST-ATG8A^ADS^ and *A. thaliana* whole seedling lysates containing mCherry-CFS1. The peptides were added to a final concentration of 200 µM. Proteins were visualized by immunoblotting with anti-GST and anti-RFP antibodies. Representative images of 3 independent replicates are shown. ADS, AIM docking site. **(B)** GST pull-down with *E. coli* lysates containing GST or GST-ATG8A and *A. thaliana* whole seedling lysates containing mCherry, mCherry-CFS1 or mCherry-CFS1^AIM^. Proteins were visualized by immunoblotting with anti-GST and anti-RFP antibodies. Representative images of 3 independent replicates are shown. **(C)** RFP-Trap pull-down of Arabidopsis seedlings co-expressing pUBQ::GFP-ATG8A and pUBQ::mCherry-CFS1 or pUBQ::mCherry-CFS1^AIM^. Seven-day-old seedlings were incubated in either control or nitrogen-deficient 1/2 MS media for 12 h. Protein extracts were immunoblotted with anti-GFP and anti-RFP antibodies. Representative images of 4 biological replicates are shown. **(D)** Phenotypic characterization of Col-0, *atg5*, *cfs1*, *cfs1* complemented with pUBQ::mCherry-CFS1 or *cfs1* complemented with pUBQ::mCherry-CFS1^AIM^ upon nitrogen starvation. Twenty-five seeds per genotype were grown on 1/2 MS media plates (+1% plant agar) for 1 week and 7-day-old seedlings were subsequently transferred to nitrogen-deficient 1/2 MS media plates (+0.8% plant agar) and grown for 2 weeks. Plants were grown at 21°C under LEDs with 85 µM/m²/sec and a 14 h light/10 h dark photoperiod. d0 depicts the day of transfer. Brightness of pictures was enhanced ≤19% with Adobe Photoshop (2020). Representative images of 4 independent replicates are shown. **(E)** Confocal microscopy images of Arabidopsis root epidermal cells co-expressing pUBQ::GFP-ATG8A and pUBQ::mCherry-CFS1 or pUBQ::mCherry-CFS1^AIM^. Five-day-old Arabidopsis seedlings were incubated in either control, nitrogen-deficient (-N) or 150 mM NaCl-containing 1/2 MS media before imaging. Area highlighted in the white-boxed region in the merge panel was further enlarged and presented in the inset panel. Scale bars, 5 μm. Inset scale bars, 2 μm. **(F)** Quantification of confocal experiments in (E) showing the number of mCherry-CFS1 puncta per normalized area (10,000 μm^2^). Bars indicate the mean ± standard deviation of 10 biological replicates. Two-tailed and paired Student t-tests were performed to analyze the significance differences of the mCherry-CFS1 puncta number. ***, p value < 0.001. **(G)** Confocal microscopy images of root epidermal cells of *cfs1* expressing pUBQ::mCherry-CFS1 or pUBQ::mCherry-CFS1^AIM^ or *cfs1*×*atg5* expressing pUBQ::mCherry-CFS1. Five-day-old Arabidopsis seedlings were incubated in either control or 150 mM NaCl-containing 1/2 MS media for 2 h before imaging. Scale bars, 5 μm. **(H)** Quantification of confocal experiments in (G) showing the number of mCherry-CFS1 puncta per normalized area (10,000 μm^2^). Bars indicate the mean ± standard deviation of 10 biological replicates. Brown-Forsythe and Welch one-way ANOVA test was performed to analyze the differences of the mCherry-CFS1 puncta number between each group. Unpaired t-tests with Welch’s correction were used for multiple comparisons. Family-wise significance and confidence level, 0.05 (95% confidence interval) was used for analysis.

We performed nitrogen starvation plate assays to test the physiological relevance of the CFS1 AIM. Expression of wild type mCherry-CFS1 rescued the nitrogen sensitivity phenotype of *cfs1* mutants. mCherry-CFS1^AIM^ expressing plants remained sensitive to nitrogen starvation, similar to the *cfs1* and *atg5* mutants (Fig. 2D). Finally, we compared the localization patterns of mCherry-CFS1 and mCherry-CFS1^AIM^ relative to GFP-ATG8A. mCherry-CFS1 formed a significantly higher number of colocalizing puncta compared to mCherry-CFS1^AIM^ under both control and autophagy inducing conditions (Fig. 2E-F). Collectively, these results suggest that CFS1 interacts with ATG8 in an AIM-dependent manner and that the CFS1-ATG8 interaction is essential for CFS1 function and autophagosome localization.

To further study the residual puncta formed by mCherry-CFS1^AIM^, we crossed mCherry-ATG8E with the *atg5* mutant, which does not form autophagosomes (Thompson et al., 2005). mCherry-CFS1 was still able to form a limited number of puncta in both basal and autophagy inducing conditions, indicating that CFS1 forms puncta independently of autophagosomes. (Fig. 2G-H). To understand how CFS1 could still form puncta in the absence of autophagy, we looked at the other functional domains on CFS1. CFS1 has well-defined FYVE and SYLF domains that bind phosphatidylinositol-3-phosphate (Pi3P) and actin, respectively (Sutipatanasomboon et al., 2017) (Fig. S9A). Mutating these domains, either individually or in combination, did not alter protein stability but did lead to diffuse localization patterns and disrupted ATG8 co-localization (Fig. S9B-C). These domain-function analyses suggest that CFS1 bridges Pi3P-rich endomembrane compartments with autophagosomes.

### CFS1 functions as an autophagy adaptor

We next explored the molecular function of CFS1 in autophagy. Three types of autophagy related proteins contain AIMs: (i) core autophagy proteins involved in autophagosome biogenesis, (ii) autophagy receptors that mediate selective cargo recruitment and undergo autophagic degradation together with their respective cargoes, and (iii) autophagy adaptors, which interact with ATG8 on the outer autophagosome membrane and mediate autophagosome trafficking and maturation (Stolz et al., 2014). To determine if CFS1 is involved in autophagosome biogenesis, we performed transmission electron microscopy (TEM) experiments in Arabidopsis root cells. Ultrastructural analysis of autophagosomes in *cfs1* mutants showed fully formed autophagosomes, indistinguishable from those in wild type cells, ruling out a role for CFS1 in autophagosome biogenesis (Fig. 3A). To test if CFS1 functions as an autophagy receptor or adaptor, we performed comparative vacuolar flux analysis, using NBR1 as a representative autophagy receptor. We stained the vacuolar lumen with BCECF-AM and blocked vacuolar degradation with the vATPase inhibitor concanamycin A (conA) (Krebs et al., 2010; Bassham, 2015), which enabled us to quantify puncta within the vacuolar lumen. Upon induction of autophagy with salt stress, we found significantly less CFS1 puncta compared to ATG8 or NBR1 puncta (Fig.3 B-C). Consistently, although both CFS1 and NBR1 colocalize with ATG8E at cytosolic autophagosomes, upon conA treatment there were fewer colocalizing CFS1-ATG8 puncta inside the vacuole compared to NBR1-ATG8 puncta (Fig.3D). These results suggest CFS1 functions as an autophagy adaptor. Since autophagy adaptors should localize on the outer autophagosome membrane, we performed immunogold-labeling TEM experiments on Arabidopsis seedlings expressing mCherry-CFS1. We could readily detect gold particles on the outer autophagosome membranes (Fig. 3F-G), consistent with CFS1 having an adaptor function. Of note, we also detected gold particles at the inner autophagosome membrane (Fig. 3F), suggesting CFS1 is recruited to the autophagosomes during phagophore growth. In sum, the comparative vacuolar flux analysis and the ultrastructural localization experiments support the role of CFS1 as autophagy adaptor.

**Figure 3.**
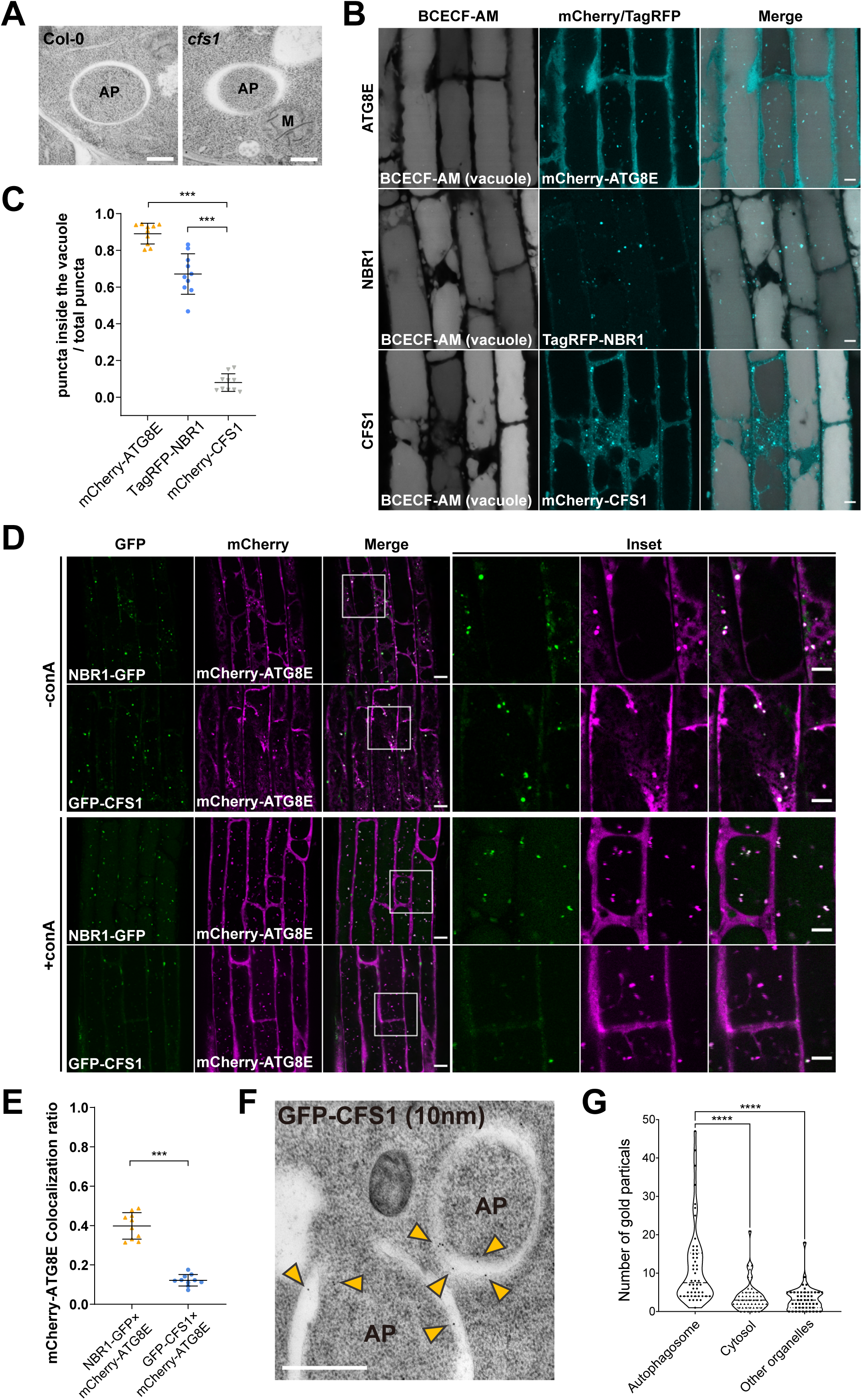
CFS1 functions as an autophagy adaptor. **(A)** Transmission electron microscopy (TEM) micrographs showing fully formed autophagosomes in the root cells of Col-0 and *cfs1*. Seven-day-old Arabidopsis seedlings were incubated in 150 mM NaCl-containing 1/2 MS media for 1 h for autophagy induction before cryofixation. Scale bars, 500 nm. AP, autophagosome. M, mitochondrion. **(B)** Confocal microscopy images of Arabidopsis root epidermal cells expressing pUBQ::mCherry-ATG8E, pNBR1::TagRFP-NBR1 or pUBQ::mCherry-CFS1. Five-day-old seedlings were first incubated in 5 μΜ BCECF-AM-containing 1/2 MS media for 30 minutes for vacuole staining and were subsequently transferred to 1/2 MS media containing 90 mM NaCl and 1 μΜ concanamycin A (conA) for 2 hours before imaging. Scale bars, 5 μm. **(C)** Quantification of confocal experiments in (B) showing the ratio between the number of mCherry-ATG8E, TagRFP-NBR1 or mCherry-CFS1 puncta inside the vacuole, compared to the total number of mCherry-ATG8E, TagRFP-NBR1, or mCherry-CFS1 puncta. Bars indicate the mean ± standard deviation of 10 biological replicates. Two-tailed and unpaired t-tests with Welch’s corrections were performed to analyze the differences of puncta numbers between mCherry-ATG8E and mCherry-CFS1 or between TagRFP-NBR1 and mCherry-CFS1. ***, p value < 0.001. **(D)** Confocal microscopy images of Arabidopsis root epidermal cells co-expressing pUBQ::mCherry-ATG8E and pUBQ::GFP-CFS1 or pNBR1::NBR1-GFP. Five-day-old Arabidopsis seedlings were incubated in 1/2 MS media containing either 150 mM NaCl (without concanamycin A; -conA), or 90 mM NaCl and 1 μΜ concanamycin A (+conA) for 2 hours before imaging. Area highlighted in the boxed region in the merge panel was further enlarged and presented in the inset panel. Scale bars, 10 μm. Inset scale bars, 5μm. **(E)** Quantification of confocal experiments in (D) showing the mCherry-ATG8E colocalization ratio of NBR1-GFP and GFP-CFS1 to mCherry-ATG8E. The mCherry-ATG8E colocalization ratio is calculated as the ratio between the number of mCherry-ATG8E puncta that colocalize with NBR1-GFP or GFP- CFS1 puncta compared with the number of total mCherry-ATG8E puncta. Bars indicate the mean ± standard deviation of 10 biological replicates. Two-tailed and unpaired t tests with Welch’s corrections were performed to analyze the differences of mCherry-ATG8E colocalization ratio between GFP-CFS1 and NBR1-GFP. ***, p value < 0.001. **(F)** TEM images showing immuno-gold labeled GFP-CFS1 at the autophagosomes in Arabidopsis root cells. Seven-day-old seedlings were incubated in 150 mM NaCl-containing 1/2 MS media for 2 h for autophagy induction before cryofixation. Sections from pUBQ::GFP-CFS1 expressing cell samples were labelled with an anti-GFP primary antibody and a secondary antibody conjugated to 10 nm gold particles. Yellow arrowheads mark the gold particles associated with autophagosomes. Scale bars, 500 nm. AP, autophagosome. **(G)** Quantification of the localization of the GFP-specific gold particles imaged in the experiment shown in (F). Approximately 900 gold particles in 50 TEM images captured from 5 independent samples were grouped into autophagosomes, cytosol or other organelles according to their locations. One-way ANOVA was performed to analyze the significant difference between different gold particle locations. ****, p value < 0.0001.

### CFS1 is crucial for autophagic flux

Autophagy adaptors regulate the delivery or fusion of autophagosomes to the vacuole, known as autophagic flux (Stolz et al., 2014).We next examined the role of CFS1 in this process. First, we expressed mCherry-ATG8E in wild type Col-0, *cfs1*, or *atg5*, and quantified the number of mCherry-ATG8E-labelled autophagosomes in root epidermal cells of these lines. Upon autophagy induction with salt stress, significantly more autophagosomes accumulated in the cytosol of *cfs1* cells compared to Col-0 (Fig. 4A-B). However, after treatment with conA, which stabilizes autophagic bodies in the vacuole, *cfs1* mutants had significantly less mCherry-ATG8E puncta (Fig. 4A-B). *atg5* mutants had no autophagosomes in either condition (Fig. 4A-B). These results suggest that *cfs1* mutants have defects in the delivery of autophagosomes to the vacuole, which leads to the accumulation of autophagosomes in the cytosol. To support these data, we performed GFP-release assays under the same conditions. When GFP-ATG8 is delivered to the vacuole, a stable GFP fragment is released due to vacuolar protease activity. The ratio of free GFP to GFP-ATG8 can thus be used to quantify autophagic flux (Bassham, 2015; Yoshii and Mizushima, 2017). Quantification of five independent experiments showed that *cfs1* had higher levels of full length GFP-ATG8A and lower levels of free GFP under nitrogen starvation or salt stress conditions compared to wild type (Fig. 4C-D). As mentioned above, the rate of NBR1 degradation can be also used to measure autophagic flux (Bassham, 2015; Yoshii and Mizushima, 2017). Following autophagy induction with salt stress NBR1 levels remained high in *cfs1* compared to wild type (Fig. 4C, 4E and S6E-H). Collectively, these results show that CFS1 is crucial for autophagic flux in *Arabidopsis thaliana*.

**Figure 4.**
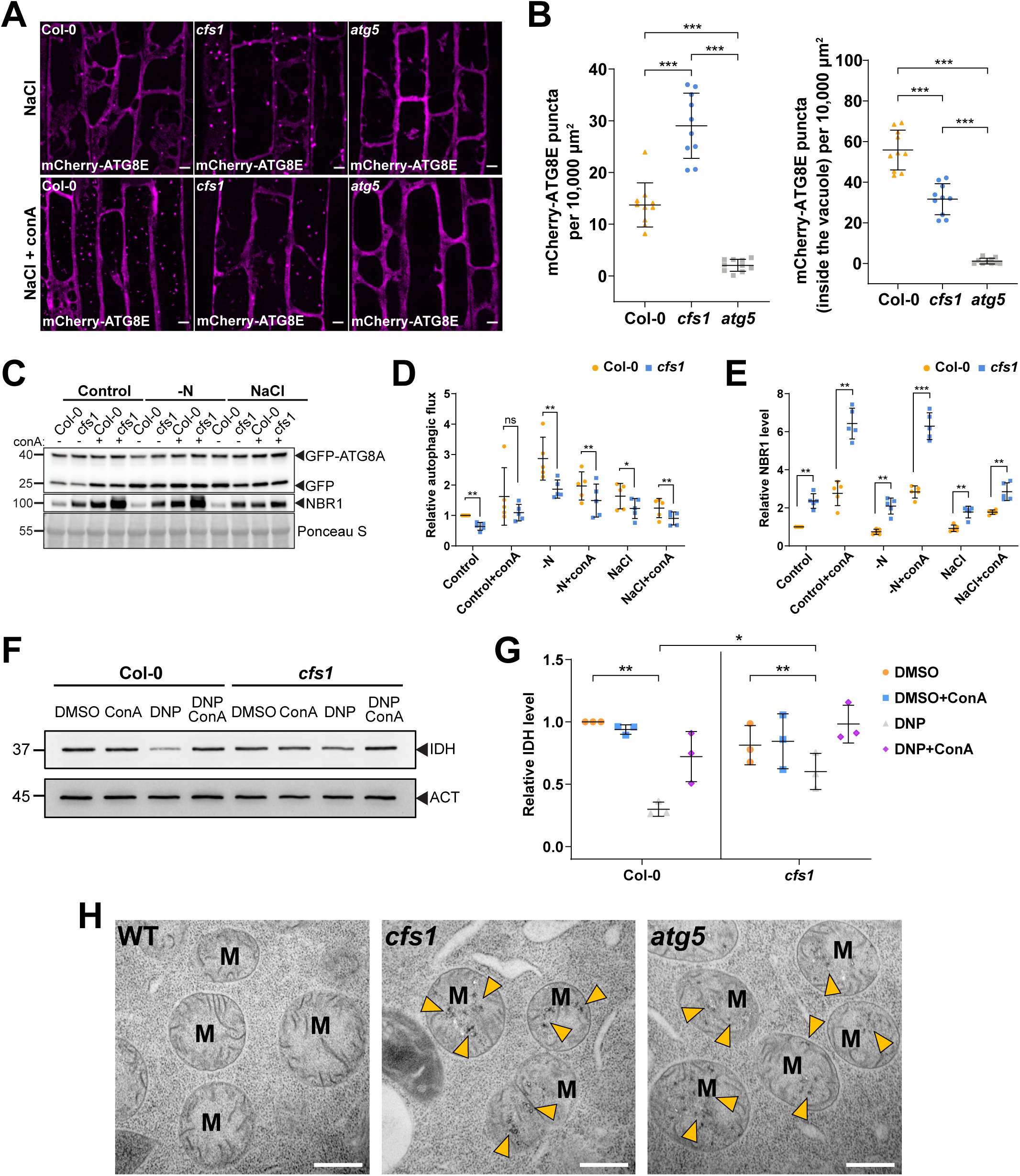
CFS1 is crucial for autophagic flux in *Arabidopsis thaliana*. **(A)** Confocal microscopy images of root epidermal cells of Col-0, *cfs1*, or *atg5* expressing pUBQ::mCherry- ATG8E under NaCl or NaCl + concanamycin A (conA) treatment. Five-day-old Arabidopsis seedlings were incubated in 1/2 MS media containing either 150 mM NaCl (NaCl) or 90 mM NaCl and 1 μΜ conA (NaCl + conA) for 2 h before imaging. Scale bars, 5 μm. **(B)** Left panel, quantification of the mCherry-ATG8E puncta per normalized area (10,000 μm^2^) of the NaCl-treated cells imaged in (A). Right panel, quantification of the mCherry-ATG8E puncta inside the vacuole per normalized area (10,000 μm^2^) of the NaCl + conA-treated cells imaged in (A). Bars indicate the mean ± standard deviation of 10 biological replicates. Two-tailed and unpaired Student t-tests with Welch’s correction were performed to analyze the significance of mCherry-ATG8E puncta density differences between Col-0 and *cfs1*, Col-0 and *atg5*, or *cfs1* and *atg5*. ***, p value < 0.001. **(C)** Western blots showing GFP-ATG8A cleavage level and endogenous NBR1 level in Col-0 or *cfs1* mutants under control, nitrogen deficient (-N) or salt stressed (NaCl) conditions. Arabidopsis seedlings were grown under continuous light in 1/2 MS media for one week and 7-day-old seedlings were subsequently transferred to 1/2 MS media ± 1 µM conA, nitrogen-deficient (-N) 1/2 MS media ± 1 µM conA, or 1/2 MS media containing 150 mM NaCl ± 1 µM conA for 12 h. Fifteen μg of total protein extract was loaded and immunoblotted with anti-GFP and anti-NBR1 antibodies. **(D)** Quantification of the relative autophagic flux in (C). Values were calculated via protein band intensities of GFP divided by GFP-ATG8A and normalized to untreated (Control) Col-0. Results are shown as the mean ± standard deviation of 5 independent replicates. One-tailed and paired Student t-tests were performed to analyze the significance of the relative autophagic flux differences. ns, not significant. *, p < 0.05. **, p < 0.01. **(E)** Quantification of the relative NBR1 level in (C) compared to untreated (control) Col-0. Values were calculated via normalization of protein bands to Ponceau S and shown as the mean ± standard deviation of 5 independent replicates. One-tailed and paired Student t-tests were performed to analyze the significance of the relative NBR1 level differences. **, p < 0.01. ***, p < 0.001. **(F)** Immunoblot assay of uncoupler-induced mitochondrial protein degradation in Arabidopsis Col-0 and *cfs1* seedlings. Seven-day-old seedlings were incubated in 1/2 MS media containing 50 μM DNP ± 1 μM conA for 4 h before protein extraction. A mitochondrial matrix protein, isocitrate dehydrogenase (IDH), was immunoblotted by anti-IDH antibodies. Actin (ACT) was immunoblotted by anti-ACT antibodies and was used as a loading control. **(G)** Quantification of relative IDH intensities in (F) compared to untreated (control) Col-0. Values were calculated via normalization of protein bands to ACT and shown as the mean ± standard deviation of 3 independent replicates. One-tailed and paired Student t-tests were performed to analyze the significance of the relative IDH level differences. ns, not significant. *, p < 0.05. **, p < 0.01. **(H)** Transmission electron microscopy (TEM) micrographs of mitochondria in Arabidopsis Col-0, *cfs1* and *atg5* root cells. Seven-day-old seedlings were incubated in 150 mM NaCl-containing 1/2 MS media for 2 h for autophagy induction before cryofixation. Yellow arrowheads mark the distinctive electron dense precipitates of compromised mitochondria that appear after NaCl treatment. Scale bars, 500 nm. M, mitochondria.

We then tested whether CFS1 is involved in autophagic flux during selective autophagy. Using our recently established uncoupler-induced mitophagy assays, we compared mitophagic flux in wild type and *cfs1* cells (Ma et al., 2021). Ultrastructural analysis of uncoupler treated *cfs1* cells showed fully formed mitophagosomes, further confirming that CFS1 does not play a role in autophagosome biogenesis (Fig. S10). We then measured mitophagic flux using western blots. Uncoupler treatment led to a decrease in levels of the mitochondrial matrix protein isocitrate dehydrogenase (IDH). This decrease was restored upon conA treatment, confirming the induction of mitophagy. *cfs1* mutants had higher IDH levels upon uncoupler treatment, suggesting a defect in mitophagic flux (Fig. 4F-G). When we examined mitochondrial ultrastructure in *cfs1* by electron microscopy, we saw accumulation of damaged mitochondria with distinctive electron dense precipitates, which were rare in wild type but common in *atg5* (Fig. 4H). Altogether, these results suggest CFS1 is also crucial for selective autophagy flux in *Arabidopsis thaliana*.

We next asked whether the function of CFS1 is conserved across plants by studying the *Marchantia polymorpha* (Mp) CFS1 homolog. Stable *M. polymorpha* plants co-expressing mScarlet-MpCFS1 with GFP-MpATG8A or GFP-MpATG8B showed that MpCFS1 colocalizes with both MpATG8 isoforms (Fig. S11A-B). Heterologous expression of MpCFS1 in *Arabidopsis thaliana* also showed colocalization of MpCFS1 with GFP-ATG8A (Fig. S11C-D). Finally, GFP-release assays in Marchantia showed that *Mpcfs1* mutants have a defect in GFP-ATG8 degradation (Fig. S11E-F). Together, these results suggest that CFS1 function is conserved across plants.

Since autophagic flux measurements assess vacuolar delivery, we decided to test if other vacuolar trafficking pathways are also affected in *cfs1* mutants. First, we measured the endocytic delivery of FM4-64 in Col-0, *cfs1*, *cfs2*, *cfs1cfs2*, and *cfs1* complementation lines. We quantified FM4-64 positive puncta upon conA treatment, which stabilizes endocytic vesicles. There was no difference between Col-0 and the mutant lines (Fig. S12A-B). We then measured the uptake of a proteinaceous endocytic cargo, the auxin efflux carrier protein PIN2, in Col-0 and *cfs1* (Kleine-Vehn et al., 2008). Similar to FM4-64, upon induction of PIN2 endocytosis with dark treatment, we did not observe any difference between Col-0 and *cfs1* (Fig. S12C-D). In addition, we compared the vacuolar morphology in Col-0, *cfs1*, *cfs2*, *cfs1cfs2*, and *cfs1* complementation lines, using BCECF-AM staining. We observed multiple small vacuoles in the epidermal cells of the root meristematic zone (Fig. S12E) and intact central vacuoles at the transition zone in all lines (Fig. S12F). Altogether, these results indicate that CFS1 specifically regulates autophagic flux without affecting other vacuolar pathways or vacuolar morphology.

### CFS1 interacts with VPS23A and mediates the formation of amphisomes

How then does CFS1 regulate autophagic flux? We hypothesized that it may interact with tethering factors, such as the CORVET or the HOPS complex, and thereby bridge the autophagosomes with the tonoplast (Takemoto et al., 2018). To test this hypothesis, we generated Arabidopsis lines that co-expressed mCherry-CFS1 with the CORVET complex component VPS3, the HOPS complex component VPS39, and the tonoplast localized SNARE protein VAMP711 (Takemoto et al., 2018; Geldner et al., 2009). Under both control and autophagy inducing conditions, CFS1 did not colocalize with any of those proteins, negating out our hypothesis (Fig. S13A-F).

This prompted us to step back and investigate the CFS1 interactome. We performed a genome-wide yeast two hybrid screen, which revealed 51 confident interactors (Table S4). Notably, one of these interactors was VPS23A, an ESCRT-I complex component that is known to regulate endomembrane trafficking (Nagel et al., 2017; Shen et al., 2018; Gao et al., 2015). Confocal microscopy analyses showed that both CFS1 and ATG8A partially colocalize with VPS23A, and the number of colocalizing puncta increases upon autophagy induction (Fig. 5A-C). Further airyscan and spinning disc microscopy analysis showed that two GFP-CFS1 puncta localize on distinct regions of VPS23A-TagRFP puncta (Fig. 5D) and CFS1 moves together with VPS23A (Supplementary video 3). To further test the association between CFS1 and the -ESCRT complex, we colocalized CFS1 with two other ESCRT-localized adaptor proteins, FREE1 and ALIX (Gao et al., 2015; Kalinowska et al., 2015). Similar to VPS23A, both proteins partially colocalized with CFS1 (Fig. S14A-B). These results suggest VPS23A could function as a hub for CFS1 labelled autophagosomes.

**Figure 5.**
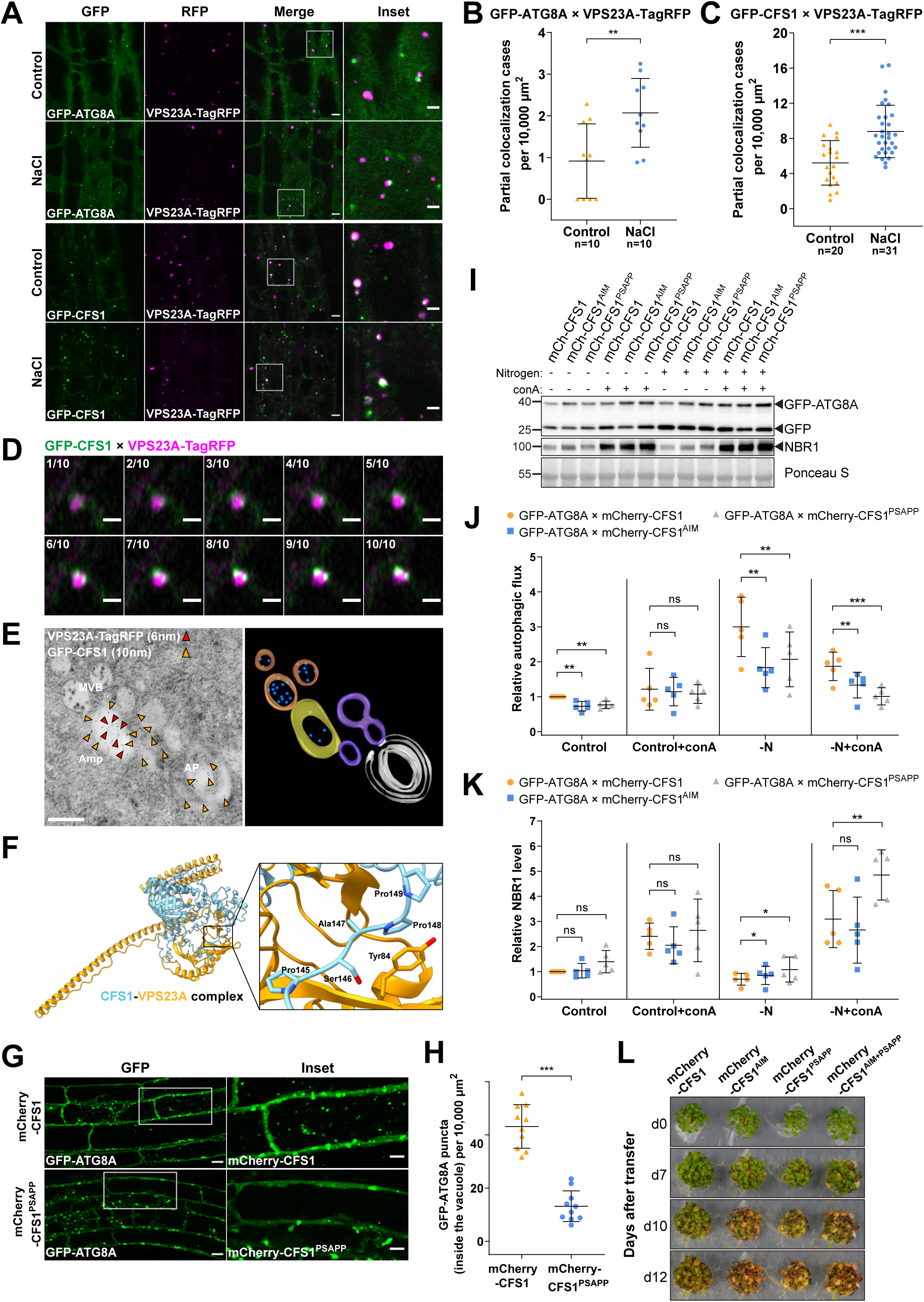
CFS1 interacts with VPS23A and mediates the formation of amphisomes. **(A)** Confocal microscopy images of Arabidopsis root epidermal cells co-expressing pVPS23A::VPS23A- tagRFP with pUBQ::GFP-ATG8A or pUBQ::GFP-CFS1. Five-day-old Arabidopsis seedlings were incubated in either control or 150 mM NaCl-containing 1/2 MS media for 2 h before imaging. Area highlighted in the white-boxed region in the merge panel was further enlarged and presented in the inset panel. Scale bars, 5 μm. Inset scale bars, 2 μm. **(B)** Quantification of confocal experiments in (A) showing the number of colocalizing cases per normalized area (10,000 μm^2^) between GFP-ATG8A and VPS23A-TagRFP. Bars indicate the mean ± standard deviation of 10 replicates. Two-tailed and unpaired Student t-tests were performed to analyze the difference of the number of colocalizing cases between control and NaCl treatment. **, p value < 0.01. **(C)** Quantification of confocal experiments in (A) showing the number of colocalizing cases per normalized area (10,000 μm^2^) between GFP-CFS1 and VPS23A-TagRFP. Bars indicate the mean ± standard deviation of 20 or 31 replicates under either control or NaCl treatment respectively. Unpaired and two-tailed Student t-tests were performed to analyze the difference of the number of colocalizing cases between control and NaCl treatment. ***, p value < 0.001. **(D)** Airyscan time-lapse microscopy images of Arabidopsis root epidermal cells showing the GFP-CFS1 puncta partially colocalizing with the VPS23A-TagRFP puncta. Five-day-old Arabidopsis seedlings co- expressing pVPS23A::VPS23A-TagRFP and pUBQ::GFP-CFS1 were incubated in 150 mM NaCl- containing 1/2 MS media for 1 h for autophagy induction before imaging. Ten continuous layers from one Z-stack image are shown separately in order. Interval of the Z-stack image, 0.15 μm. Scale bars, 1 μm. **(E)** Left panel, transmission electron microscopy micrographs showing the colocalization of VPS23A- TagRFP and GFP-CFS1 in Arabidopsis root cells co-expressing pVPS23A::VPS23A-TagRFP and pUBQ::GFP-CFS1. Five-day-old seedlings were incubated in 150 mM NaCl-containing 1/2 MS media for 2 h for autophagy induction before cryofixation. Sections were labelled with rabbit anti-RFP and chicken anti-GFP primary antibodies and secondary antibodies conjugated to 6 nm or 10 nm gold particles. Yellow and red arrowheads mark the representative RFP and GFP-specific gold particles respectively. MVB, multivesicular bodies; Amp, amphisome; AP, autophagosome. Scale bars, 400 nm. Right panel, the three- dimensional model of the amphisome and its associated compartments shown in the left panel. The model was generated from serial sections flanking the section shown in the left panel. Autophagosome (white), autophagosome associated vesicles (purple), amphisome (yellow), multivesicular bodies (orange) and internal vesicles (blue) were rendered into 3D surfaces. **(F)** Homology modelling of CFS1/Vps23A complex. Prediction of CFS1-VPS23A heterocomplex formation generated by AlphaFold2 as implemented by ColabFold (Jumper et al., 2021; Mirdita et al., 2021). Structure of CFS1 and VPS23A are represented as ribbons and colored in light blue and orange, respectively. The predicted complex interaction interface involving AtCFS1 PSAPP motif is highlighted as a zoom in with the side chains of relevant residues represented as sticks. **(G)** Confocal microscopy images of Arabidopsis root epidermal cells co-expressing pUBQ::GFP-ATG8A with pUBQ::mCherry-CFS1 or pUBQ::mCherry-CFS1^PSAPP^ under NaCl + concanamycin A (conA) treatment. Five-day-old Arabidopsis seedlings were incubated in 1/2 MS media containing 90 mM NaCl and 1μΜ conA for 2 h before imaging. Scale bars, 10 μm. Inset scale bars, 5 μm. **(H)** Quantification of GFP-ATG8A puncta inside the vacuole per normalized area (10,000 μm^2^) of the cells imaged in (G). Bars indicate the mean ± standard deviation of 10 biological replicates. Two-tailed and unpaired Student t-test was performed to analyze the significance of difference between mCherry-CFS1 and mCherry-CFS1^PSAPP^. ***, p value < 0.001. **(I)** Western blots showing the GFP-ATG8A cleavage level and the endogenous NBR1 level in Arabidopsis *cfs1* mutants co-expressing pUBQ::GFP-ATG8A with pUBQ::mCherry-CFS1, pUBQ::mCherry-CFS1^AIM^, or pUBQ::mCherry-CFS1^PSAPP^ under control ± conA or nitrogen-deficient (-N) ± conA conditions. Arabidopsis seeds were first grown in 1/2 MS media under continuous light for one week and 7-day-old seedlings were subsequently transferred to 1/2 MS media ± 1 µM conA or nitrogen-deficient (-N) 1/2 MS media ±1 µM conA for 12 h. Ten μg of total protein extract was loaded and immunoblotted with anti-GFP and anti-NBR1 antibodies. **(J)** Quantification of (I) showing the relative autophagic flux. Values were calculated as the protein band intensities of GFP divided by the protein band intensity of GFP-ATG8A and were normalized to untreated (Control) Col-0. Results are shown as the mean ± standard deviation of 5 independent replicates. One- tailed and paired Student t-tests were performed to analyze the significance of the relative autophagic flux differences. ns, not significant. **, p < 0.01. ***, p < 0.001. **(K)** Quantification of (I) showing the relative NBR1 level in respect to untreated (Control) Col-0. Values were calculated via normalization of protein bands to Ponceau S and shown as the mean ± standard deviation of 5 independent replicates. One-tailed and paired Student t-tests were performed to analyze the significance of the relative NBR1 level difference. ns, not significant. *, p < 0.05. **, p < 0.01. **(L)** Phenotypic characterization of Arabidopsis *cfs1* mutants complemented with pUBQ::mCherry-CFS1, pUBQ::mCherry-CFS1^AIM^, pUBQ::mCherry-CFS1^PSAPP^ or pUBQ::mCherry-CFS1^AIM+PSAPP^ upon nitrogen starvation. Twenty-five seeds per genotype were grown on 1/2 MS media plates (+1% plant agar) for 1 week and 7-day-old seedlings were subsequently transferred to nitrogen-deficient 1/2 MS media plates (+0.8% plant agar) and grown for 2 weeks. Plants were grown at 21°C under LEDs with 85 µM/m²/sec and a 14 h light/10 h dark photoperiod. d0 depicts the day of transfer. Brightness of pictures was enhanced ≤19% with Adobe Photoshop (2020). Representative images of 4 independent replicates are shown.

We then performed immunogold labeling transmission electron microscopy experiments to visualize the compartments where CFS1 and VPS23A colocalize. We used two differently sized gold particles conjugated to anti-GFP or anti-RFP antibodies, recognizing GFP-CFS1 and VPS23A-TagRFP, respectively. Electron micrographs obtained from cyto-fixed Arabidopsis root cells revealed that CFS1 and VPS23A colocalized at amphisomes, hybrid structures where autophagosomes fuse with intraluminal vesicle containing multivesicular bodies (Sanchez-Wandelmer and Reggiori, 2013) (Fig. 5E) This demonstrates that, similar to metazoans, plants have amphisomes where CFS1 colocalizes with VPS23A.

This prompted us to test the significance of amphisomes in autophagic flux. We first measured autophagic flux in *vps23* mutants. Arabidopsis has two VPS23 isoforms, VPS23A/VPS23.1 and VPS23B/VPS23.2 (Nagel et al., 2017). Since Arabidopsis *vps23avps23b* double mutants are lethal, we measured autophagic flux on single *vps23a* or *vps23b* mutants (Nagel et al., 2017). We expressed GFP-ATG8A in Col-0, *cfs1*, *vps23a*, and *vps23b* and quantified the GFP-ATG8A-marked autophagic bodies with and without salt stress and conA treatment. Neither *vps23a* nor *vps23b* mutants showed a significant difference compared to Col-0 (Fig. S15A-B). Consistently, neither *vps23a* nor *vps23b* mutants showed autophagic flux defects in NBR1 degradation assays (Fig. S15C-D). Finally, *vps23a* and *vps23b* were indistinguishable from wild type in nitrogen starvation plate assays (Fig. S15E). Altogether, these data indicate that VPS23A and VPS23B may act redundantly or do not play a role in autophagy.

To circumvent the potential redundancy between VPS23A and VPS23B, we mutated CFS1 residues likely to mediate the VPS23A-CFS1 interaction. Previous studies have shown that VPS23 interacting proteins contain a “PSAPP” motif, which docks into the ubiquitin E2 variant domain on VPS23 (Pornillos et al., 2003; Sutipatanasomboon et al., 2017). Consistently, Alphafold prediction suggested CFS1 and VPS23A interacted in a PSAPP dependent manner (Jumper and Hassabis, 2022) (Fig. 5F). We thus generated the mCherry-CFS1^PSAPP^ mutant, where the PSAPP motif is mutated to alanines (PSAPP-145-AAAAA) and expressed it in the *cfs1* background. Western blot analysis showed that this mutation did not affect the stability of mCherry-CFS1 (Fig. S16). We then measured autophagic flux in mCherry-CFS1 and mCherry-CFS1^PSAPP^ expressing *Arabidopsis thaliana* lines, using three different assays. First, we quantified vacuolar GFP-ATG8A puncta under autophagy inducing conditions. mCherry-CFS1^PSAPP^ lines had significantly fewer autophagic bodies in the vacuole (Fig. 5G-H). Consistently, GFP-release assays showed that mCherry-CFS1^PSAPP^ lines phenocopied mCherry-CFS1^AIM^ lines, with an autophagic flux defect under nitrogen starvation conditions (Fig. 5I-J). Finally, measurement of NBR1 also indicated a defect in autophagic flux (Fig. 5I, 5K). Ultimately, to test the physiological importance of the VPS23A-CFS1 interaction, we performed nitrogen starvation plate assays. Similar to the expression of mCherry-CFS1^AIM^, expression of mCherry-CFS1^PSAPP^ failed to rescue the nitrogen starvation-sensitivity phenotype of *cfs1* (Fig. 5L). Altogether, these findings demonstrate that the CFS1-VPS23A interaction is critical for autophagic flux and loss of interaction leads to early senescence.

## Discussion

Autophagosome maturation involves the trafficking and fusion of double-membraned autophagosomes with lytic compartments. Studies in yeast and metazoans have shown that this maturation step has several similarities with endocytic vesicle fusion and trafficking. Tethering complexes, RAB GTPases, SNARE proteins, and adaptors facilitate the trafficking and fusion of autophagosomes with the endolysosomal compartments (Zhao and Zhang, 2019; Zhao et al., 2021). While well studied in yeast and metazoans, a systematic analysis of autophagosome maturation is still missing in plants. In addition, plant genomes lack homologs of key maturation proteins, such as the autophagic SNARE, Syntaxin 17, suggesting plants have evolved different components for autophagosome maturation. Here, we show that CFS1 is an autophagy adaptor that bridges autophagosomes with multivesicular bodies and mediates the formation of amphisomes. We propose that CFS1 functions as a licensing factor that tethers autophagosomes to multivesicular bodies via the ESCRT-1 complex. This is reminiscent of COPII tethering factor p115 that targets a subpopulation of COPII vesicles to cis-Golgi (Allan et al., 2000).

Our findings are consistent with a role in autophagosome maturation as opposed to autophagosome biogenesis as was recently suggested (Kim et al., 2022): (i) our differential centrifugation-based autophagosome enrichment procedure selects for proteins that associate with closed, mature autophagosomes. AP-MS did not identify biogenesis components but did identify trafficking related proteins such as Myosin 14 (Fig 1B, Table S3), (ii) we observe fully formed autophagosomes in micrographs obtained from *cfs1* mutants indistinguishable from and in similar numbers to wild type (Fig. 3A, Fig. S10), and (iii) we observe a clear colocalization with the ESCRT-1 protein VPS23A in both confocal and electron micrographs (Fig. 5). Although we see significant defects in bulk and selective autophagic flux in *cfs1*, CFS1^AIM^, and CFS1^PSAPP^ mutants, autophagic flux is not fully blocked in any of them. These findings suggest two, not mutually exclusive, explanations: 1) There could be other autophagy adaptors that mediate the maturation of a sub-population of autophagosomes. Consistently, in metazoans, there are several autophagy adaptors that are either involved in SNARE recruitment, Rab GTPase activation or autophagosome trafficking (Zhao et al., 2021). Performing autophagosome enrichment experiments in a GFP-ATG8A/*cfs1* line could reveal these adaptors. 2) Some autophagosomes may be trafficked directly to the vacuole without an intermediary amphisome step. A systematic characterization of the autophagosome fusion machinery would shed light on both of these possibilities.

There are multiple trafficking routes to the vacuole: endocytic trafficking, post-Golgi trafficking, ER to vacuole transport, and autophagy (Aniento et al., 2022). Although we are starting to understand how these routes operate individually, how distinct vacuolar trafficking pathways are coordinated in response to changes in metabolic demands and external stimuli remains elusive Here, based on our findings, we propose that vacuolar trafficking is organized as a hub and spoke type distribution system, where amphisomes serve as sorting hubs for multivesicular bodies and autophagosomes (Figure 6). The hub and spoke model was developed by Delta Airlines in the 1950s, and has since then been successfully employed by various types of supply chain logistics and aviation industry. The model implies that, rather than transportation taking place from A to B as in the point-to-point system, all materials transit through centralized hubs. It thereby reduces logistical costs as fewer routes are necessary. It also permits economies of scale, since complicated operations can be performed in the hubs rather than separately organized in each node (Oti, 2013). Organising vacuolar trafficking via hubs (i) could allow the cell to intricately balance the anabolic and catabolic needs and seamlessly integrate various intrinsic and extrinsic signals, (ii) could facilitate the crosstalk between post-Golgi trafficking, endocytosis, and autophagy, and (iii) could provide a route for loading and secretion of extracellular vesicles.

**Figure 6.**
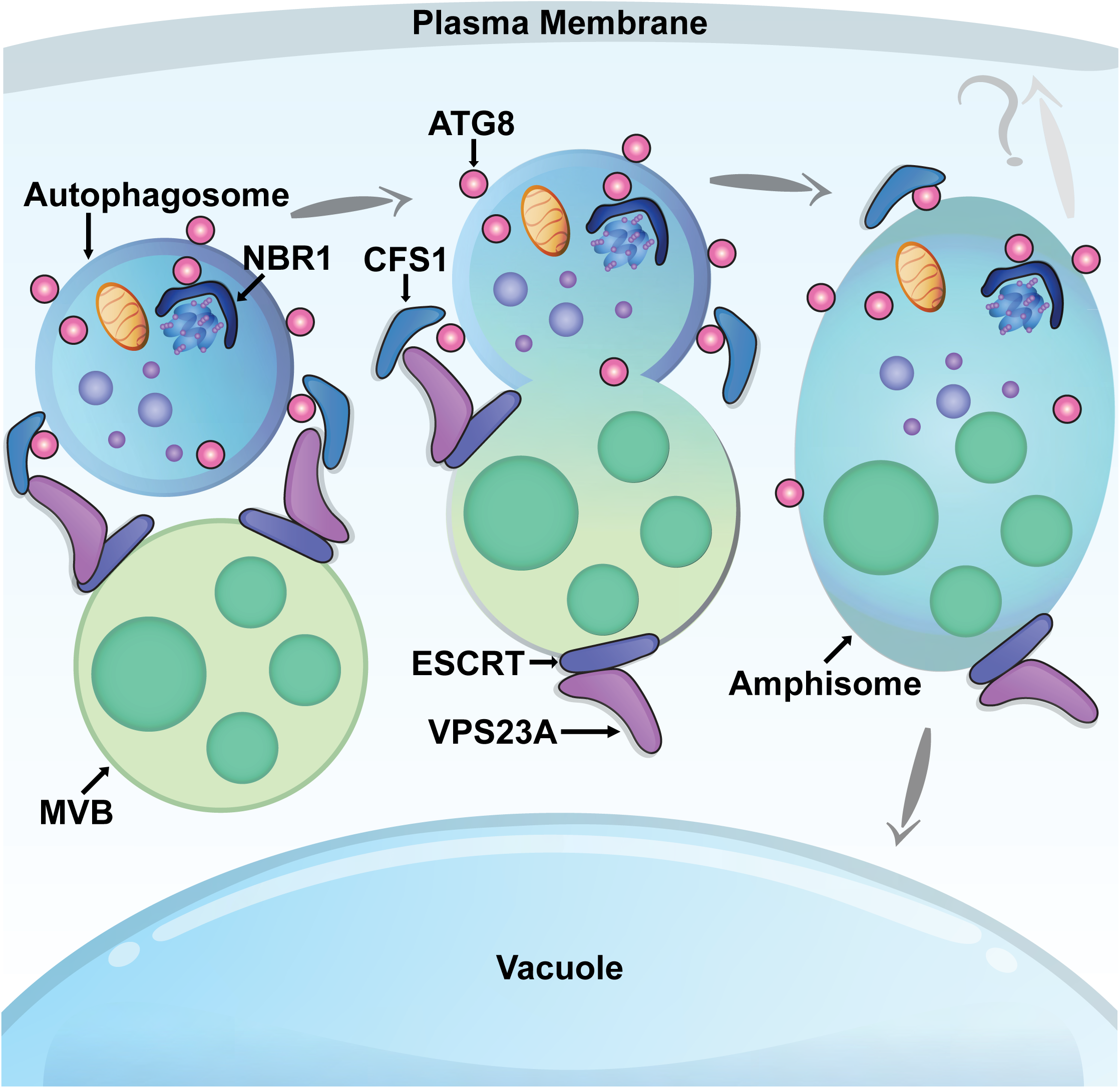
Current working model: Vacuolar trafficking is organized as a hub and spoke type distribution system. CFS1 interacts with both ATG8 and VPS23A and mediates the formation of amphisomes −hybrid prevacuolar compartments that are formed by the fusion of multivesicular bodies and autophagosomes.

One well-known drawback of hub and spoke distribution models is that the hub represents a single point of failure. Any congestion in the hub can severely impact the whole system. Therefore, building a robust hub is critical for a successful supply chain (Oti, 2013). This could be informative for translational approaches that aim to increase plant stress tolerance by engineering plant endomembrane trafficking.

### Speculation

One of our time-lapse videos analyzing CFS1-NBR1 co-trafficking showed an intriguing pattern: A CFS1-NBR1 positive punctum squeezed through a membrane, which looks like movement from one cell to another (Supplementary video 4). We would like to share it with the community and discuss the implications of cell-to-cell movement of autophagosomes: (i) In Arabidopsis roots, central vacuoles mature and acidify by fusion of small vacuoles in a developmentally regulated manner (Cui et al., 2019, 2020; Krüger and Schumacher, 2018). So, in meristematic cells vacuoles may not be hydrolytically active. Autophagosomes in such cells may need to traffic to the neighboring cells to deliver autophagic cargo to an active lytic compartment. (ii) Alternatively, they may be loaded with cargo that are involved in cell-to-cell communication. As the title of the section implies, these scenarios need to be supported or refuted by further studies.

## Acknowledgements

We thank Suayip Ustun, Karin Schumacher, Erika Isono, Gerd Juergens, Takashi Ueda, Daniel Hofius, Liwen Jiang for sharing published materials. We thank Lorenzo Picchianti for drawing the model presented in Figure 6. We acknowledge funding from Austrian Academy of Sciences, Austrian Science Fund (FWF, P 32355, P 34944), Austrian Science Fund (FWF-SFB F79), Vienna Science and Technology Fund (WWTF, LS17-047) to YD; Austrian Academy of Sciences DOC Fellowship to JZ, Marie Curie VIP2 Fellowship to JCDC and MC; Hong Kong Research Grant Council (GRF14121019, 14113921, AoE/M-05/12, C4002-17G) to BHK. We thank Vienna Biocenter Core Facilities (VBCF) Protein Chemistry, Biooptics, Plant Sciences, Molecular Biology, and Protein Technologies. We thank J. Matthew Watson and members of the Dagdas lab for the critical reading and editing of the manuscript.

The authors declare no competing financial interests.

## Data availability

All the raw images, blots, replicates associated with figures are uploaded to the Dryad server and can be accessed under the DOI: doi.org/10.5061/dryad.s7h44j18f and doi.org/10.5061/dryad.w0vt4b8th. Proteomics data are available via ProteomeXchange with identifier PXD031787.

## Author contributions

JZ performed microscopy experiments, flux measurements, prepared figures and wrote the draft. MTB generated Arabidopsis lines, performed flux and protease protection, and phenotypic characterization, and AP-MS experiments and wrote the draft. JM performed electron microscopy experiments and mitophagy flux assays, prepared figures and wrote the draft. FK generated Arabidopsis lines. JCDC performed phylogenetic analysis and Alphafold models, wrote the draft. YC contributed to protease protection assays. SP contributed to phenotypic characterization experiments. AM, MGL, MSG performed the Marchantia related experiments. CG performed the time-lapse imaging experiments. DG performed the PIN2 endocytosis experiments. RK performed the AP-MS experiments. MC performed the IP-MS experiments, analyzed data, prepared figures and wrote the draft. SS generated transgenic Arabidopsis lines. YJ and JF supervised students and contributed to the draft. BHK supervised students, wrote the draft. YD supervised students and postdocs, wrote the draft.

## Materials and methods

### Plant material and cloning procedure

All *Arabidopsis thaliana* lines used in this study originate from the Columbia (Col-0) ecotype and are listed below (Table 1). The primers used for genotyping are listed in Table 4. Coding sequences from gene of interest were amplified from Col-0 cDNA with primers listed in Table 4. Plasmids were assembled via the GreenGate cloning method (Lampropoulos et al., 2013). In short, CFS1 (At3g43230) and CFS2 (At1g29800) were cloned in two parts to remove internal *Bsa*I sites (see Table 4 for primers). For introducing point mutations in mCherry-CFS1^AIM^ (WLNL-267-ALNA), mCherry-CFS1^FYVE^ (RHHCR-195-AHACA) and mCherry-CFS1^PSAPP^ (PSAPP-145-AAAAA) site-directed mutagenesis was applied using primers listed in Table 4. For mCherry-CFS1^SYLF^ (K282A, R288A, K320A), the SYLF domain was mutagenized by ordering a synthetic DNA sequence carrying point mutations (Table 4) and replaced by restriction enzyme digestion with *Nco*I (NEB) and *Xba*I (NEB). All constructs used in this study are listed in Table 3. Transgenic plant lines were generated via the Agrobacterium-mediated floral-dip method (Clough and Bent, 1998) and are listed in Table 1.

**Table 1.**
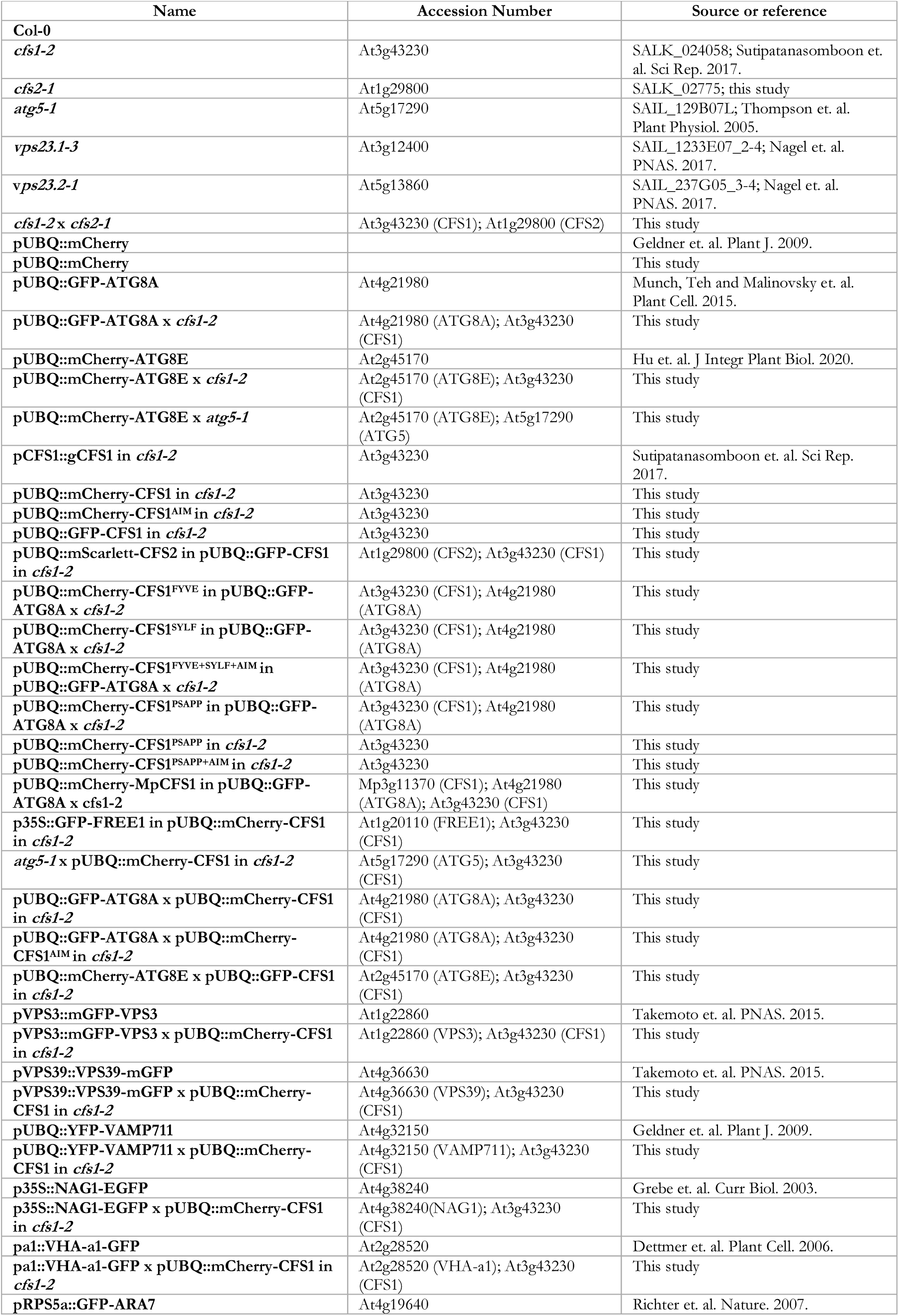
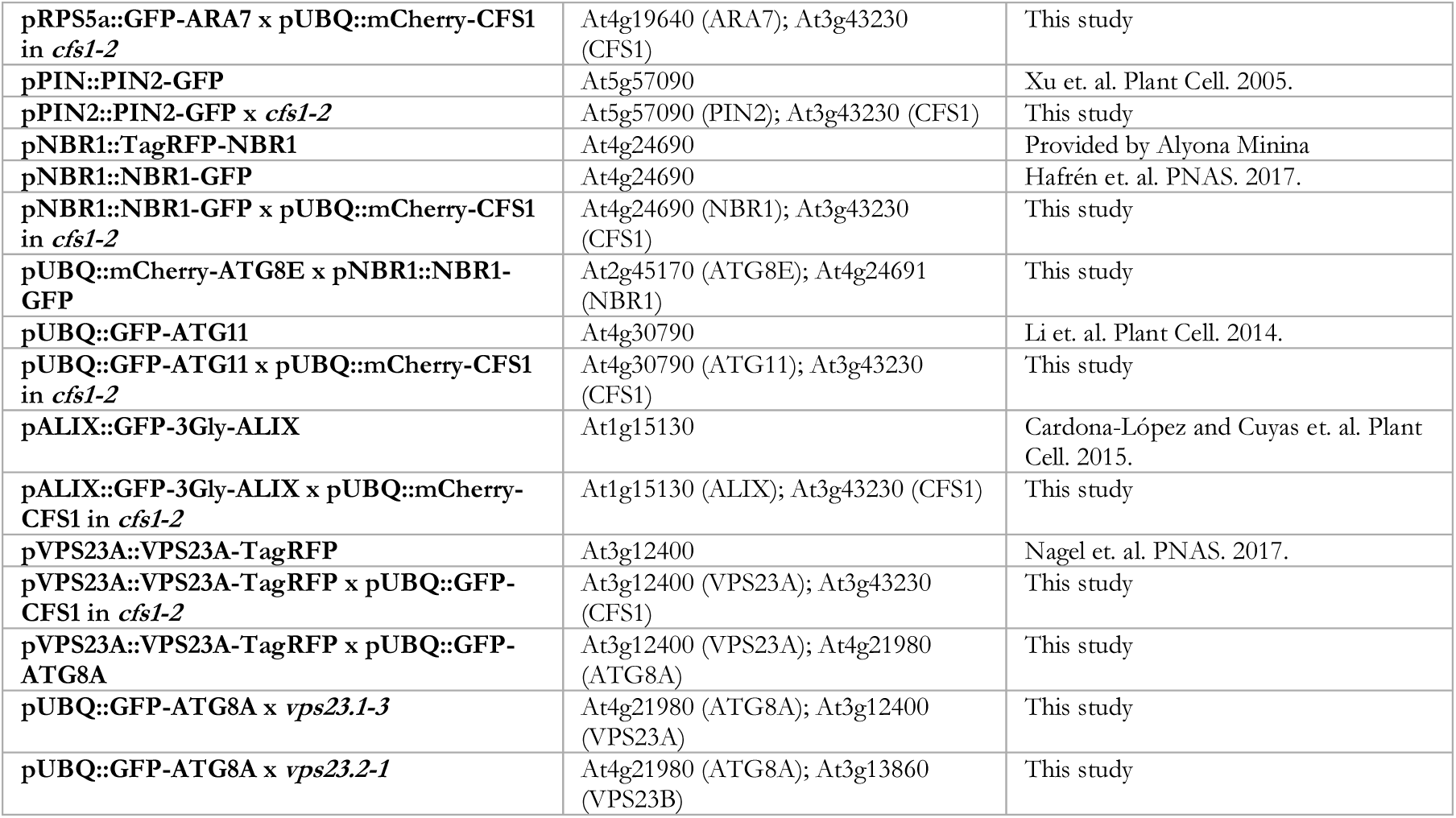
Arabidopsis thaliana lines used in this study.

For *Marchantia polymorpha*, all lines are derived from the Takaragaike-1 (Tak-1) ecotype. Plants were maintained and cultivated asexually cultivated on half-strength (1/2) Gamborg B5 media (Gamborg B5 medium basal salt mixture (Duchefa) supplemented with 0.5 g/L MES and 1% sucrose, pH 5.5) plates (+ 1% plant agar (Duchefa)) under continuous white light with a light intensity of 50 µM/m²/sec at 21°C. For cloning MpATG8A (Mp1g21590) and MpATG8B (Mp5g05930) primers are listed in Table 4. For MpCFS1 (Mp3g11370) a synthetic DNA construct (Table 4) with GreenGate compatible *Bsa*I overhangs was ordered. Plasmids were assembled via the Gateway (Ishizaki et al., 2015) or GreenGate (Lampropoulos et al., 2013) system and introduced through *Agrobacterium tumefaciens* transformation (Kubota et al., 2013). *cfs1* knockout mutants were achieved through CRISPR/Cas9 gene editing via 2 gRNAs. gRNAs were inserted at the *Bsa*I (NEB) sites of the pMpGE_En03 entry vector and subsequently inserted into the pMpGE010 destination vector via LR clonase II reaction (Invitrogen) (Sugano et al., 2018). Successful transformants were verified via sequencing of PCR products from amplified genomic DNA. Stable plant lines and constructs used are listed in Table 2 and Table 3 respectively.

**Table 2.** Marchantia polymorpha lines used in this study.

| Name | Accession Number | Source or reference |
| --- | --- | --- |
| Tak-1 |  |  |
| pEF::GFP-ATG8A | Mp1g21590 | This study |
| *cfs1* in pEF::GFP-ATG8A | Mp3g11370 (CFS1); Mp1g21590 (ATG8A) | This study |
| pEF::mScarlet-CFS1 in pEF::GFP-ATG8A | Mp3g11370 (CFS1); Mp1g21590 (ATG8A) | This study |
| pEF::GFP-ATG8B | Mp5g05930 | This study |
| *cfs1* in pEF::GFP-ATG8B | Mp3g11370 (CFS1); Mp5g05930 (ATG8B) | This study |
| pEF::mScarlet-CFS1 in pEF::GFP-ATG8B | Mp3g11370 (CFS1); Mp5g05930 (ATG8B) | This study |

**Table 3.** List of plasmids used in this study.

| Table 3. List of plasmids used in this study. |  |  |  |  |  |
| --- | --- | --- | --- | --- | --- |
| Name | Accession number | Expression System | Backbone | Additional information | Source or reference |
| pUBQ::mCherry |  | *A. thaliana* | pGGZ003 |  | This study |
| pUBQ::mCherry-CFS1 | At3g43230 | *A. thaliana* | pGGZ003 |  | This study |
| pUBQ::GFP-CFS1 | At3g43231 | *A. thaliana* | pGGZ003 |  | This study |
| pUBQ::mCherry-CFS1<sup>AIM</sup> | At3g43232 | *A. thaliana* | pGGZ003 | WLNL267ALNA | This study |
| pUBQ::mCherry-CFS1<sup>FYVE</sup> | At3g43233 | *A. thaliana* | pGGZ003 | RHHCR195AHACA | This study |
| pUBQ::mCherry-CFS1<sup>SYLF</sup> | At3g43234 | *A. thaliana* | pGGZ003 | K282A, R288A, K320A | This study |
| pUBQ::mCherry-CFS1<sup>FYVE+SYLF+AIM</sup> | At3g43235 | *A. thaliana* | pGGZ003 | RHHCR195AHACA (FYVE); K282A, R288A, K320A (SYLF); WLNL267ALNA (AIM) | This study |
| pUBQ::mCherry-CFS1<sup>PSAPP</sup> | At3g43236 | *A. thaliana* | pGGZ003 | PSAPP145AAAAA (PSAPP) | This study |
| pUBQ::mCherry-CFS1<sup>PSAPP+AIM</sup> | At3g43237 | *A. thaliana* | pGGZ003 | PSAPP145AAAAA (PSAPP); WLNL267ALNA (AIM) | This study |
| pUBQ::mScarlett-CFS2 | At1g29800 | *A. thaliana* | pGGZ003 |  | This study |
| pUBQ::mCherry-MpCFS1 | Mp3g11370 | *A. thaliana* | pGGZ003 |  | This study |
| p35S::GFP-FREE1 | At1g20110 | *A. thaliana* | pMDC43 |  | Provided by Pedro L. Rodriguez |
| GST |  | *E. coli* |  |  | Stephani and Picchianti et al., 2020. |
| GST-ATG8 | At4g21980 | *E. coli* |  |  | Stephani and Picchianti et al., 2020. |
| GST-ATG8A<sup>ADS</sup> | At4g21980 | *E. coli* |  | YL50AA | Stephani and Picchianti et al., 2020. |
| pEF::GFP-ATG8A | Mp1g21590 | *M. polymorpha* | pMpGWB303 |  | This study |
| pEF::GFP-ATG8B | Mp5g05930 | *M. polymorpha* | pMpGWB303 |  | This study |
| pEF::mScarlet-CFS1 | Mp3g11370 | *M. polymorpha* | pGGSUN |  | This study |
| pMpGE010_CFS1g1g2 | Mp3g11370 | *M. polymorpha* | pMpGE010 |  | This study |

**Table 4.**
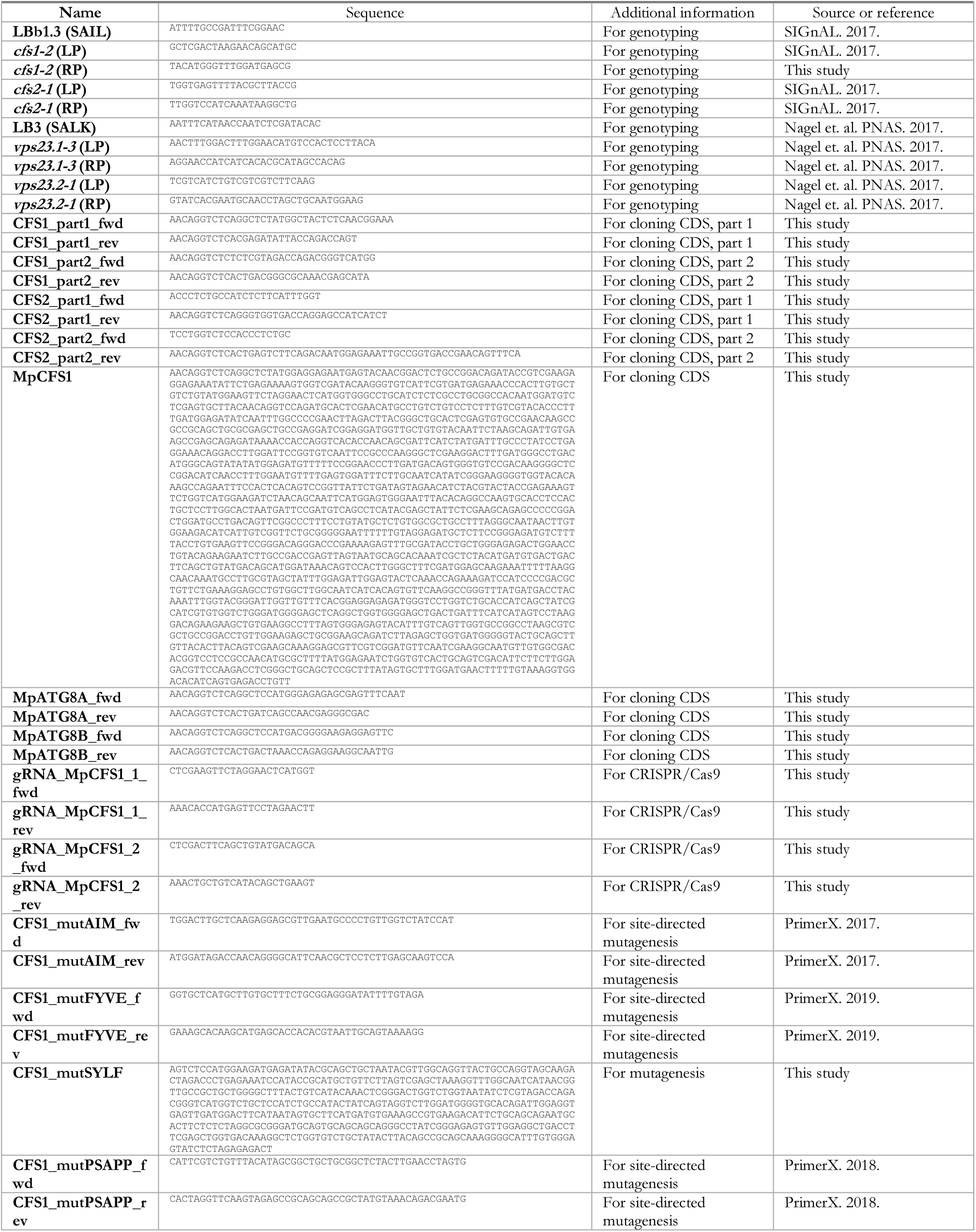
Primer and synthetic sequences used in this study for genotyping, cloning and mutagenesis.

### Plant phenotypic assays

Arabidopsis seedlings were grown as described in Jia et al., 2019 (Jia et al., 2019). Briefly, twenty-five Arabidopsis seeds per bundle were vapor-phase sterilized (90% sodium hypochlorite 13% and 10% HCl 36%) and sown on plates which were layered with a thin nylon mesh on top of 1/2 MS media (Murashige and Skoog salt + Gamborg B5 vitamin mixture (Duchefa) supplemented with 0.5 g/L MES and 1% sucrose, pH 5.7) plates (with 1% plant agar (Duchefa)) followed by 3 days of vernalization at 4 °C in dark. Vernalized seeds were grown at 21 °C under LEDs with 85 µM/m²/sec with a 14 h light/10 h dark photoperiod for 7 d. Seven-day-old seedlings were subsequently transferred to 1/2 MS media or nitrogen-deficient 1/2 MS media (Murashige and Skoog salt without nitrogen + Gamborg B5 vitamin mixture (Duchefa) supplemented with 0.5 g/L MES and 1% sucrose, pH 5.7) plates (+ 0.8% plant agar (Duchefa)) and grown for 2 weeks.

### Sample preparation before protein extraction

Unless stated otherwise, twenty to forty *A. thaliana* seeds were grown in 1/2 MS media in 12-well plates under continuous light and constant shaking at 80 rpm for 7 d. Seven-day-old seedlings were subjected to indicated treatments. For nitrogen starvation, 1/2 MS media was replaced with nitrogen-deficient 1/2 MS media. For salt stress, 150 mM NaCl was added to 1/2 MS media. For drug-treatments, all drugs used were dissolved in DMSO and added to the desired concentration (3 μM Torin (CAS 1222998-36-8; Santa Cruz); 1 μM concanamycin A (conA) (CAS 80890-47-7; Santa Cruz); 50 μM 2,4-dinitrophenol (DNP) (CAS 51-28-5; SIGMA)). Equal amount of pure DMSO was added to control samples. Seedlings were harvested in microcentrifugation tubes with different sized glass beads (2.85-3.45 mm, 1.7-2.1 mm and 0.75-1.00 mm; Lactan GmbH) and flash frozen in liquid nitrogen. Plants were ground with a mixer mill MM400 (3x 30 s, 30 Hz; Retsch).

For protein extraction of *M. polymorpha*, the propagules were grown for 10 d in 1/2 Gamborg B5 media under continuous light with a light intensity of 50 µM/m²/sec at 21°C and subjected to indicated treatments. Samples were flash frozen in liquid nitrogen and ground using a mortar and pestle.

### Western blotting

Arabidopsis protein extraction was achieved by adding 250 μl of protein extraction buffer (100 mM Tris (pH 7.5), 200 mM NaCl, 1 mM EDTA, 2% 2-Mercaptoethanol, 0,2% Triton X-100 and 1 tablet/50 ml Complete, EDTA-free Protease Inhibitor Cocktail (SIGMA), pH 7.8) to grinded samples. Samples were subsequently well mixed and centrifuged at 15000 rpm at 4°C for 10 min. The whole supernatant was transferred to a new tube. After another round of centrifugation at 15000 rpm at 4°C for 10 min, resulting supernatant was diluted with 2x Laemmli buffer (4% SDS, 20% glycerol, 10% 2-mercaptoethanol, 0.004% bromophenol blue and 0.125 M Tris HCl, pH 6.8) and boiled at 95°C for 10 min as the final protein extract sample.

For *M. polymorpha* protein extraction, grinded plant powder was weighed and added with 2x volume of buffer (10% glycerol, 25 mM Tris (pH 7.5), 150 mM NaCl, 1 mM EDTA, 2% PVPP, 1 mM DTT, 0.2% Nonidet P-40/Igepal and 1 tablet/50 ml cOmplete, EDTA-free Protease Inhibitor Cocktail (SIGMA)) for lysing. Protein lysates were cleared via centrifuging samples at 12000 rpm twice. Resulting supernatant was diluted in 4x Laemmli buffer and boiled at 70 °C for 10 min as the final protein extract sample.

For Arabidopsis, protein concentration was measured using the Amido black method (Popov et al., 1975). Ten μl of protein sample was diluted in 190 μl water and added with 1 ml Amido black staining solution (90% methanol, 10% acetic acid, 0.005% (w/v) Amido black 10B (SIGMA)). Samples were mixed thoroughly and centrifuged at 15000 rpm for 10 min. After removal of supernatant, pellets were washed with 1 ml washing solution (90% ethanol and 10% acetic acid) and centrifuged at 15000 rpm for 10 min. Resulting pellets were dissolved in 1 ml 0.2 N NaOH and the corresponding optical density (OD) at 630 nm was measured via a plate reader (Synergy™ HTX Multi-Mode Microplate Reader; BioTek). Protein concentration (C) was calculated via the formula C = (OD-b)/10a, where a and b were calibrated by the Bovine Serum Albumin (BSA) standard curve of the staining solution. For Marchantia, protein concentration was quantified using Bradford assay (SIGMA). Chemiluminescence was acquired via iBright Imaging System (Invitrogen)

For western blotting, indicated amount of protein was loaded on 4-20% Mini-PROTEAN® TGX™ precast gel (Bio-Rad) and blotted on nitrocellulose using the semi-dry Trans-Blot® Turbo™ Transfer System (Bio-Rad). Protein extract was immunoblotted with either anti-NBR1 (1:2000; Agrisera), anti-IDH (1:5000; Agrisera), anti-GFP (1:2000; SIGMA) or anti-RFP (1:2000; Chromotek) antibodies. Images were captured via ChemiDoc™ Touch Imaging System (Bio-Rad) by developing with SuperSignal™ West Pico PLUS Chemiluminescent Substrate (Thermo Fischer). Protein band intensities were quantified via Image Lab 6 (Bio-Rad) as previously described in Stephani and Picchianti et al., 2020 (Stephani et al., 2020).

### Protease protection assay

Roughly 5-10 mg of Arabidopsis seeds were grown in 6-well plates under continuous light and continuous shaking for 7 d. Seven-day-old Arabidopsis seedlings were subjected to Torin treatment (3 μM for 90 min) for autophagy induction and immediately grinded in GTEN-based buffer (GTEN (10% glycerol, 30 mM Tris (pH 7.5), 150 mM NaCl, 1 mM EDTA (pH 8)), 0.4 M sorbitol, 5 mM MgCl2, 1 mM Dithiothreitol (DTT), 100x liquid protease inhibitor cocktail (SIGMA) and 1% Polyvinylpolypyrrolidon (PVPP)) in a 3:1 v/w ratio. Afterwards lysates underwent several differential centrifugation steps where each time the supernatant was transferred. Samples were spun for (1) 10 min at 1000 g, to remove cell debris and nuclei; (2) 10 min at 10000 g, to remove bigger organelles like mitochondria and chloroplasts; (3) 10 min at 15000 g, to further remove organelles (S3 fraction) and finally (4) 60 min at 100000 g (S4 and P4 fraction) (LaMontagne et al., 2016). Protein concentration in S3 was normalized via Bradford (SIGMA) to ensure that equal amount of protein was loaded before ultracentrifugation step. The P4 fraction was dissolved gently in GTEN-based buffer (without PVPP) and was further subjected to 30 ng/μl proteinase K (SIGMA) and 1% Triton X-100 treatment for 30 min on ice. The reaction was stopped with 5 mM phenylmethylsulfonyl fluoride (PMSF). Proteins were then precipitated overnight with 0.1% sodium deoxycholate (NaDOC) and 11% trichloroacetic acid (TCA). Resulting pellets were washed twice with 100% acetone and dissolved in 2x Laemmli buffer. Proteins were quantified again via the Amido black method as described above and 5 μg were loaded on the gel.

### *In vitro* pull-downs

Recombinant proteins were expressed using Rosetta2(DE3)pLysS *E. coli* strain. Bacteria were grown to an OD_600_ of 0.6 - 0.7 followed by induction with 300 mM IPTG and overnight incubation at room temperature (RT). 50 ml of culture was pelleted and resuspended in 5 ml of EDTA-free buffer (10% Glycerol, 25 mM Tris/HCl pH 7.5, 150 mM NaCl, 0.1 mM TCEP, 0.1% Nonidet P-40/Igepal, 10 μM ZnCl_2_ and 1 tablet/50 ml cOmplete, EDTA-free Protease Inhibitor Cocktail (SIGMA)) containing 10x FastBreak™ Cell Lysis reagent (Promega) and Benzonase. Cells were broken open via incubation on a spinning wheel at room temperature for 10-20 min. Lysates were clarified via centrifugation at 15000 rpm at 4 °C for 10 min. For proteins expressed *in planta*, forty seeds per genotype were grown. Frozen plant tissue was homogenized as described above and 700 μl EDTA-free buffer containing 2% PVPP was added. Lysate were centrifuged twice at 15000 rpm for at 4 °C 10 min. 100 μl of *E. coli* lysate and 400 μl of plant lysate was mixed and incubated with 10 μl of equilibrated Glutathione High-Capacity Magnetic Agarose Beads (SIGMA) for 1 h at 4 °C on a spinning wheel. For peptide competition, peptides were added to a final concentration of 200 μM as described in Stephani and Picchianti et al., 2020. Beads were washed 5 times with EDTA-free buffer, without TCEP, eventually eluted in 50 μl 2x Laemmli buffer and boiled for 10 min at 95 °C. Gels were loaded with 15 μl of sample per well. Anti-GST HRP conjugate (SIGMA) were used in a 1:2000 dilution.

### *In vivo* co-immunoprecipitation

For co-immunoprecipitation, forty seeds per genotype were grown in 1/2 MS media for 7 d. Proteins were extracted by adding 800 μl of EDTA-free buffer with 2% PVPP. Lysates were cleared by centrifugation at 15000 rpm at 4 °C for 10 min twice. 500 μl supernatant was incubated with 20 μl RFP-Trap^®^ Magnetic Agarose beads (Chromotek) for 1 h. Beads were washed 3 and 5 times with EDTA-free buffer, without TCEP, before and after incubation with lysate respectively. Beads were eluted in 50 μl 2x Laemmli buffer, boiled for 10 min at 95°C and subjected to western blot analysis with indicated antibodies.

### Affinity purification-mass spectrometry (AP-MS)

For affinity purification, S4, P4, and P4 + proteinase K samples described in Fig. 1A were prepared as same as the method described above for the protease protection sample preparation, except that samples were incubated for 1 h with 40 μl RFP-Trap^®^ Magnetic Agarose beads (Chromotek) after proteinase K treatment. Mass spectrometry sample preparation and measurement were performed as previously described in Stephani and Picchianti et al., 2020 (Stephani et al., 2020).

### Mass spectrometry data processing

The total number of MS/MS fragmentation spectra was used to quantify each protein (Table S1). The data matrix of spectral count values (Table S2) was submitted to a negative-binomial test using the R package IPinquiry4 (https://github.com/hzuber67/IPinquiry4) that calculates fold change and p-values using the quasi-likelihood negative binomial generalized log-linear model implemented in the edgeR package. The pairwise comparisons were the following: 1-mCherry P4 *vs.* mCherry-ATG8E P4, 2-mCherry-ATG8E S4 *vs.* mCherry-ATG8E P4 and 3- mCherry-ATG8E P4 + protease K *vs.* mCherry-ATG8E P4. In each case, comparisons were obtained from two independent biological replicates. Only proteins with log(FC)>0 and p value<0.05 were considered to build the Venn diagram in Fig. 1B. Venn diagram was built using the Venny 2.1.0 online tool (https://bioinfogp.cnb.csic.es/tools/venny/index.html) and then redrawn manually. Analysis results are shown in Table S3. The mass spectrometry proteomics data have been deposited to the ProteomeXchange Consortium via the PRIDE (Perez-Riverol et al., 2019) partner repository.

### Yeast two hybrid screening

The screening was performed by Hybrigenics against the Arabidopsis cDNA library. Results are shown in Table S4.

### Preparation of *Arabidopsis thaliana* samples for confocal microscopy

For all experiments except PIN2 endocytosis imaging, Arabidopsis seeds were vapor-phase sterilized by chlorine (generated by a 10:1 mixture of 13% sodium hypochlorite and 36% HCl) for 15 min and were subsequently stored at 4°C for 2 d for vernalization. Vernalized seeds were spread on 1/2 MS media plates (+ 1% plant agar (Duchefa)) and grown at 21°C at 60% humidity under LEDs with 50 mM/m^2^s a and a 16 h light/8 h dark photoperiod for 5 d. Plates were placed vertically to let the roots elongate along the media surface. Five-day-old seedlings were placed in 1/2 MS media and treated with salt or chemicals as indicated in each experiment before confocal imaging. For nitrogen starvation, 1/2 MS media was replaced by nitrogen-deficient 1/2 MS media. For PIN2 endocytosis imaging, Arabidopsis seeds were vapor-phase sterilized by chlorine (generated by a 25:1 mixture of 2.6% sodium hypochlorite and 36% HCl) for 3 to 4 hours. The sterilized seeds were spread on 1/2 MS media plates (+ 1% plant agar (Duchefa)). The plated seeds were subsequently stored at 4°C for 2 d for vernalization. Vernalized seeds were then grown at 21°C at 60% humidity under LEDs with 50 mM/m^2^s a and a 16 h light/8 h dark photoperiod for 5 d. Plates were placed vertically to let the roots elongate along the media surface. Five-day-old Arabidopsis seedlings were grown on 1/2 MS media plates under light or 6 h dark conditions before imaging.

For confocal microscopy, Arabidopsis seedlings were placed on a microscope slide with water or water with 0.002 mg/ml propidium iodide and covered with a coverslip. The epidermal cells of root meristem zone were used for CFS1 and FM 4-64 colocalization imaging, PIN2 endocytosis imaging, or vacuolar morphology imaging. The epidermal cells of root elongation zone were used for confocal imaging in Fig. 3D and time-lapse imaging in Supplementary videos 1-3. For all other experiments, the epidermal cells of root transition zone were used for image acquisition.

### Preparation of *Marchantia polymorpha* samples for confocal microscopy

The *Marchantia polymorpha* asexual gemmae were incubated in 1/2 Gamborg B5 media for 2 d before imaging. Two-day-old *M. polymorpha* thalli were placed on a microscope slide with water and covered with a coverslip. The apical meristem region was used for image acquisition.

### Confocal microscopy

All images except for Airyscan imaging, PIN2 endocytosis imaging and time-lapse imaging were acquired by an upright point laser scanning confocal microscope ZEISS LSM800 Axio Imager.Z2 (Carl Zeiss) equipped with high-sensitive GaAsP detectors (Gallium Arsenide), a LD C-Apochromat 40X objective lens (numerical aperture 1.1, water immersion) and ZEN software (blue edition, Carl Zeiss). GFP and BCECF-AM fluorescence were excited at 488 nm and detected between 488 nm and 545 nm. TagRFP, mCherry and mScarlet fluorescence was excited at 561 nm and detected between 570 nm and 617 nm. FM 4-64 fluorescence was excited at 561 nm and detected between 656 nm and 700 nm (Fig. S4A) or 576 nm and 700 nm. For Z-stack imaging, interval between layers was set as 1 μm. For each experiment, all replicate images were acquired using identical confocal microscopic parameters. Confocal images were processed with Fiji (version 1.52, Fiji) and Imaris (version 9.0.1, BITPLANE).

For Airyscan imaging, Z-stack images were acquired by an inverted point laser scanning confocal microscope ZEISS LSM980 Axio Observer.Z1/7 (Carl Zeiss) equipped with a hexagonal GaAsP detector Airyscan II (Gallium Arsenide), a plan-Apochromat 63x objective lens (numerical aperture 1.40, oil immersion) and ZEN software (blue edition, Carl Zeiss) under room temperature. GFP fluorescence was excited at 488 nm and detected between 495 nm and 550 nm. TagRFP fluorescence was excited at 561 nm and detected between 573 nm and 627 nm. Interval between layers was set as 0.15 μm. Original images were deconvoluted by ZEN software using the default mode (blue edition, Carl Zeiss). Deconvoluted images were processed with Fiji (version 1.52, Fiji).

For PIN2 endocytosis imaging, images were acquired by an inverted Zeiss microscope AxioObserver Z1 (Carl Zeiss) equipped with a spinning disk module CSU-W1-T3 (Yokogawa), a Prime 95B camera (http://www.photometrics.com/), a Plan-Apochromat 63x objective lens (numerical aperture 1.4, oil immersion) and the Metamorph acquisition software (Molecular Devices). GFP was excited with a 488 nm laser (150mW) and was detected by a 525/50 nm BrightLine® single-band bandpass filter (Semrock). Confocal images were processed with Fiji (version 1.52, Fiji).

For time-lapse imaging, images were acquired by an Andor Dragonfly confocal platform (Andor) equipped with a spinning disc confocal microscope (Nikon Ti2E inverted microscope). A 40X objective lens (numerical aperture 1.15, water immersion) was installed for acquiring images. The pinhole disk pattern was set as 40 μm. Green fluorescence signals were excited by 488 nm laser and detected via an Andor Zyla sCMOS camera. Red fluorescence signals were excited by 561 nm laser and detected via an Andor iXon 888 EMCCD camera. The two cameras acquire the image at the same time. For time-lapse mode imaging, the total imaging time is 60 seconds, with an interval of 1 second. Confocal images were processed with Fiji (version 1.52, Fiji).

### Image processing and quantification

Mander’s colocalization analyses were performed by Fiji (version 1.52, Fiji). Confocal images were background-subtracted with 25 pixels of rolling ball radius. Mander’s coefficients M1 and M2 were calculated through JACoP plugin (Bolte and Cordelieres, 2006). Thresholds in JACoP plugin settings were adjusted according to the puncta signals in original confocal images.

Puncta quantification was performed by Fiji (version 1.52, Fiji). Z-stack images (at least 5 layers) were background-subtracted with 25 pixels of rolling ball radius. Each Z-stack image was subsequently thresholded using the MaxEntropy method and was converted to an 8-bit gray-scale image. Threshold values were adjusted according to the puncta signals in original confocal images. The number of puncta in thresholded images were counted by the Analyze Particles function in Fiji. For all puncta quantification, puncta with the size between 0.10-4.00 μm^2^ were counted.

Imaris (version 9.0.1, BITPLANE) was used for 3D construction of the vacuole structure. Default settings were used for 3D construction while the surface signals were smoothed with a surface detail of 0.25 μm.

### Transmission electron microscopy (TEM)

For TEM samples preparation, high-pressure freezing, freeze substitution, resin embedding, and ultramicrotomy were performed as described before (Ma et al., 2021; Kang, 2010; Wang et al., 2017). Briefly, seven-day-old Arabidopsis seedlings were incubated in 150 mM NaCl or 50 μM DNP for 1 hour and were then rapidly frozen with an EM ICE high-pressure freezer (Leica Microsystems).

For Epon resin embedded samples, the samples were freeze-substituted in 2% OsO_4_ with acetone at -80 °C for 24 h and were then slowly warmed up to room temperature over 48 h. Excess OsO_4_ was removed by rinsing with acetone at room temperature. Root samples were separated from planchettes and embedded in Embed- 812 resin (Electron Microscopy Sciences, Cat. No. 14120) and the resin was polymerized at 65 °C. Ultrathin (100 nm) sections from the samples were collected on copper slot grids coated with formvar.

For immuno-gold labeling samples, frozen specimens were freeze substituted in anhydrous acetone containing 0.25% glutaraldehyde and 0.1% uranyl acetate at −80 °C for 24 h and were slowly warmed up to −45 °C. After rinsing with precooled acetone, the cells were embedded in Lowicryl HM20 resin at −45 °C and the resin was polymerized by ultraviolet illumination. Ultrathin (100 nm) sections from the samples were collected on nickel slot grids coated with formvar. The sections were probed with rabbit anti-ATG8 (Agrisera), rabbit anti-RFP (Abcam) or chicken anti-GFP polyclonal primary antibodies (Abcam) and gold particles (6 nm, 10 nm) conjugated secondary antibodies (Ted Pella). Sections were post-stained and examined with a Hitachi 7400 TEM (Hitachi-High Technologies) operated at 80 kV.

### Statistical analyses

All quantification analyses and statistical tests were performed with GraphPad Prism 8 software. F-test was used to check whether the variances were significantly different (p < 0.05). For comparing the significance of differences between two experimental groups, student’s t-tests were performed as indicated in each experiment. The significance level of differences between two experimental groups was marked as *, p<0.05; **, p<0.01; ***, p<0.001; ns, not significant. For comparing the significance of differences between multiple experimental groups, one-way ANOVA was performed as indicated in each experiment.

For PIN2 endocytosis qualitative quantification, the Arabidopsis seedling with at least 5 root epidermal cells that contain visible PIN2-GFP in the vacuole is considered as the one with high PIN2 endocytic activities.

### Phylogenic tree analysis

Coding sequences of CFS1 and CFS2 homologs were obtained by using the BLAST tool against representative species of different plant lineages in Phytozome (Goodstein et al., 2012). The tree was inferred from a 2283- nt-long alignment by using the Maximum Likelihood method and Tamura-Nei model as implemented by MEGA X (Tamura and Nei, 1993; Kumar et al., 2018). 100 bootstrap method and a discrete Gamma distribution was used to model evolutionary rate differences among sites (5 categories (+G, parameter = 0.8072)) (Felsenstein, 1985). Best-scoring ML tree (−98686.19) is shown with purple circles indicating bootstrap values above 80 on their respective clades. The tree is represented using Interactive Tree Of Life (iTOL) v4 (Letunic and Bork, 2019).

### Homology modelling

Cartoon representations of AtCFS1 structure and prediction of AtCFS1-Vps23A heterocomplex formation were generated by homology modelling using AlphaFold2 (Jumper et al., 2021) as implemented by ColabFold (Mirdita et al., 2021).

---

**Figure S1.**
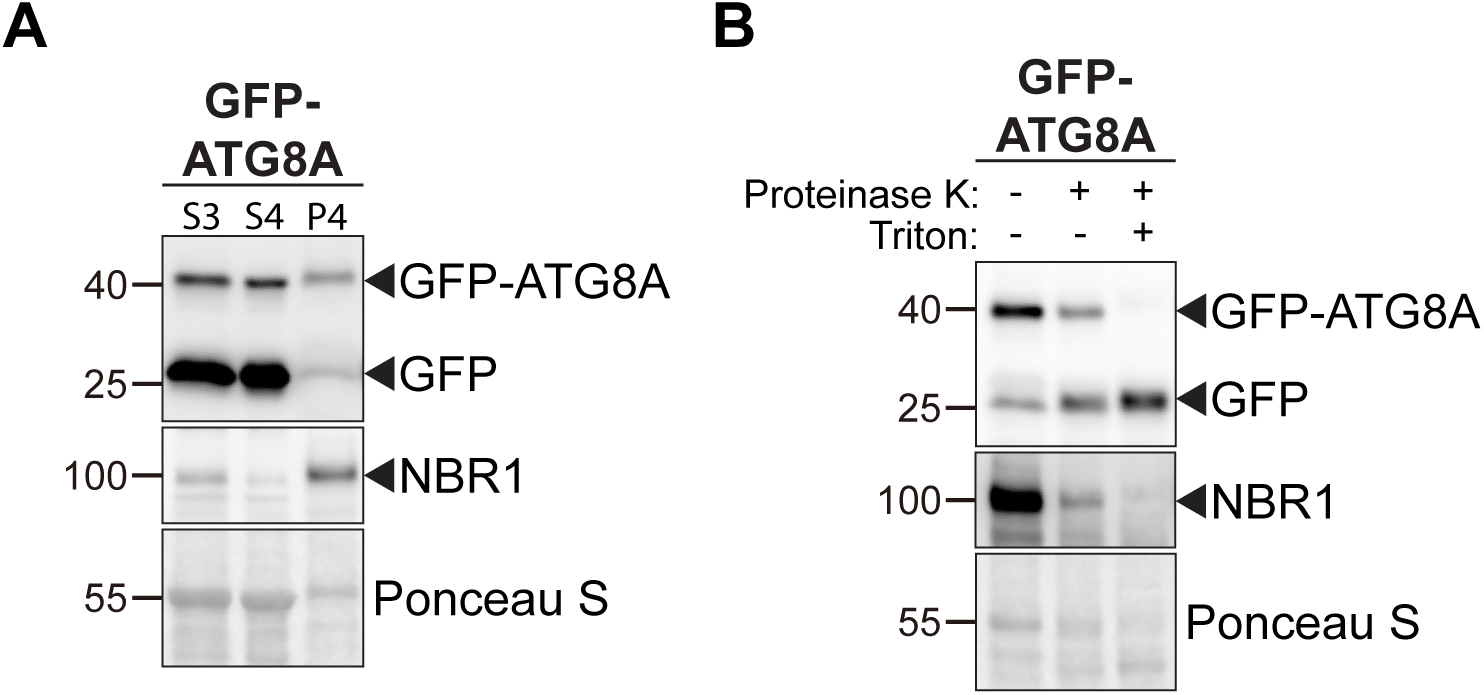
Ultracentrifugation enriches for intact autophagosomes. **(A)** Western blot analysis of 7-day-old Col-0 seedlings expressing pUBQ::GFP-ATG8A. Arabidopsis seedlings were treated with 3 μM Torin for 90 min prior to differential centrifugation described in Figure 1A. A total of 5 μg of protein was loaded in each lane. Protein extracts were immunoblotted with anti-GFP and anti-NBR1 antibodies. **(B)** Protease protection assay of enriched autophagosomes in (A). Autophagosomes were treated with 30 ng/μl proteinase K in absence or presence of 1% Triton X-100. A total of 5 μg of protein was loaded in each lane. Protein extracts were immunoblotted with anti-GFP and anti-NBR1 antibodies.

**Figure S2.**
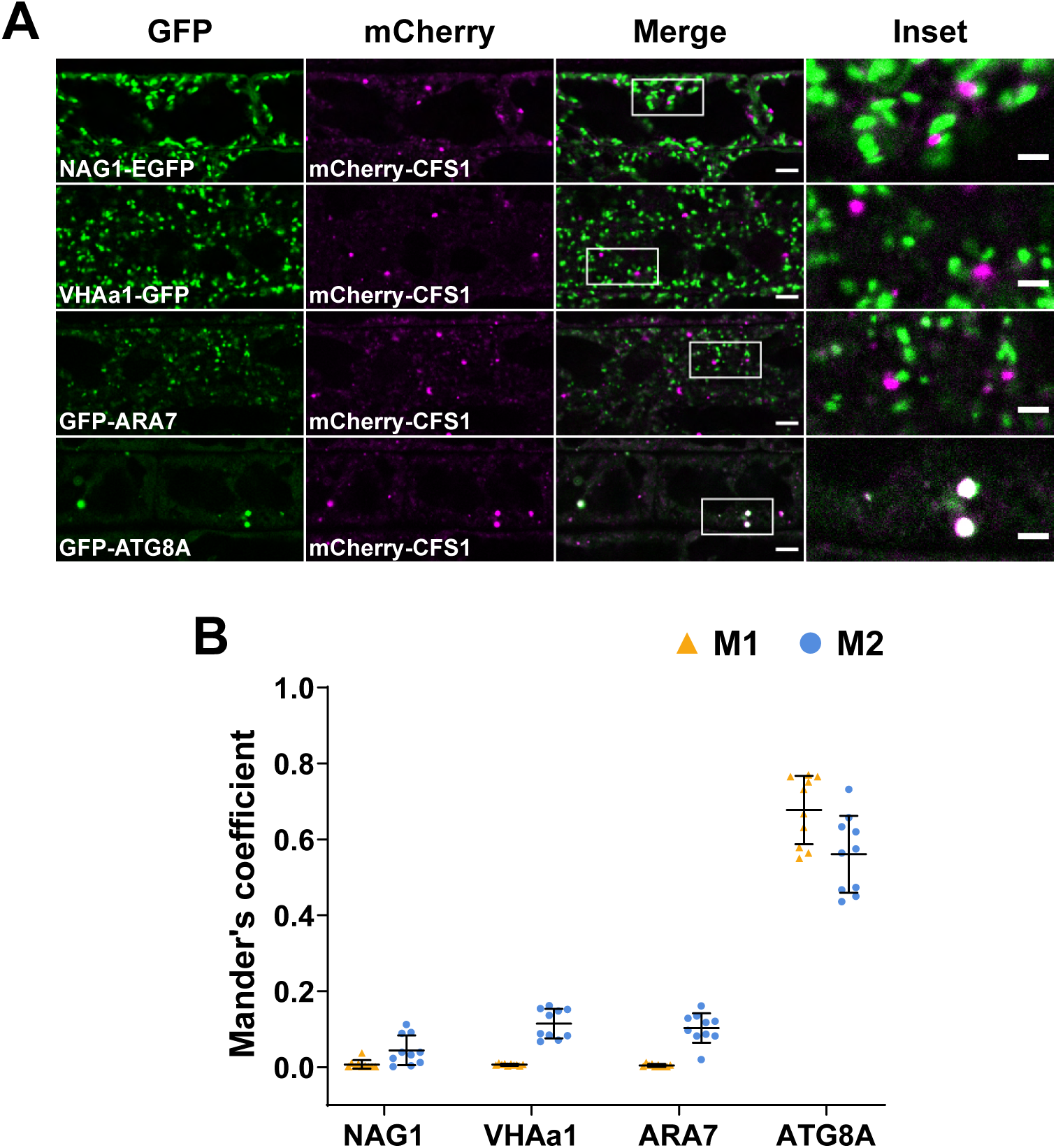
CFS1 localizes to the autophagosomes under control conditions. **(A)** Confocal microscopy images of Arabidopsis root epidermal cells co-expressing pUBQ::mCherry-CFS1 with either Golgi body marker p35S::NAG1-EGFP, trans-Golgi network marker pa1::VHAa1-GFP, late endosome marker pRPS5a::GFP-ARA7 or autophagosome marker pUBQ::GFP-ATG8A under control conditions. Five-day-old Arabidopsis seedlings were incubated in control 1/2 MS media before imaging. Area highlighted in the white-boxed region in the merge panel was further enlarged and presented in the inset panel. Scale bars, 5 μm. Inset scale bars, 2 μm. **(B)** Quantification of confocal experiments in (A) showing the Mander’s colocalization coefficients between mCherry-CFS1 and the GFP-fused marker NAG1, VHAa1, ARA7 or ATG8A. M1, fraction of GFP-fused marker signal that overlaps with mCherry-CFS1 signal. M2, fraction of mCherry-CFS1 signal that overlaps with GFP-fused marker signal. Bars indicate the mean ± standard deviation of 10 biological replicates.

**Figure S3.**
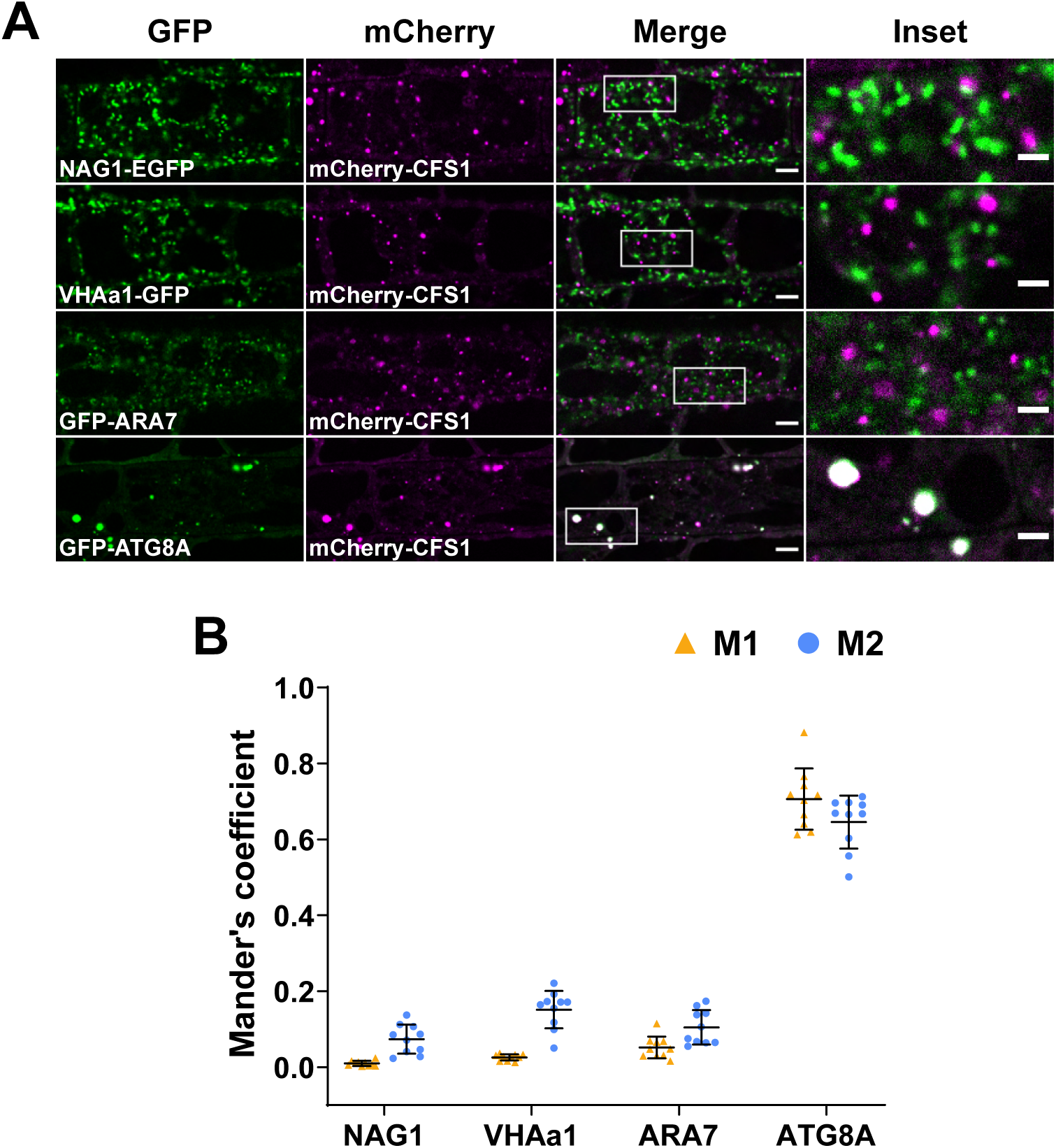
CFS1 localizes to the autophagosomes under salt stress. **(A)** Confocal microscopy images of Arabidopsis root epidermal cells co-expressing pUBQ::mCherry-CFS1 with either Golgi body marker p35S::NAG1-EGFP, trans-Golgi network marker pa1::VHAa1-GFP, MVB marker pRPS5a::GFP-ARA7 or autophagosome marker pUBQ::GFP-ATG8A under salt stress. Five-day- old Arabidopsis seedlings were incubated in 150 mM NaCl-containing 1/2 MS media for 1 h for autophagy induction before imaging. Area highlighted in the white-boxed region in the merge panel was further enlarged and presented in the inset panel. Scale bars, 5 μm. Inset scale bars, 2 μm. **(B)** Quantification of confocal experiments in (A) showing the Mander’s colocalization coefficients between mCherry-CFS1 and the GFP-fused marker NAG1, VHAa1, ARA7 or ATG8A. M1, fraction of GFP-fused marker signal that overlaps with mCherry-CFS1 signal. M2, fraction of mCherry-CFS1 signal that overlaps with GFP-fused marker signal. Bars indicate the mean ± standard deviation of 10 biological replicates.

**Figure S4.**
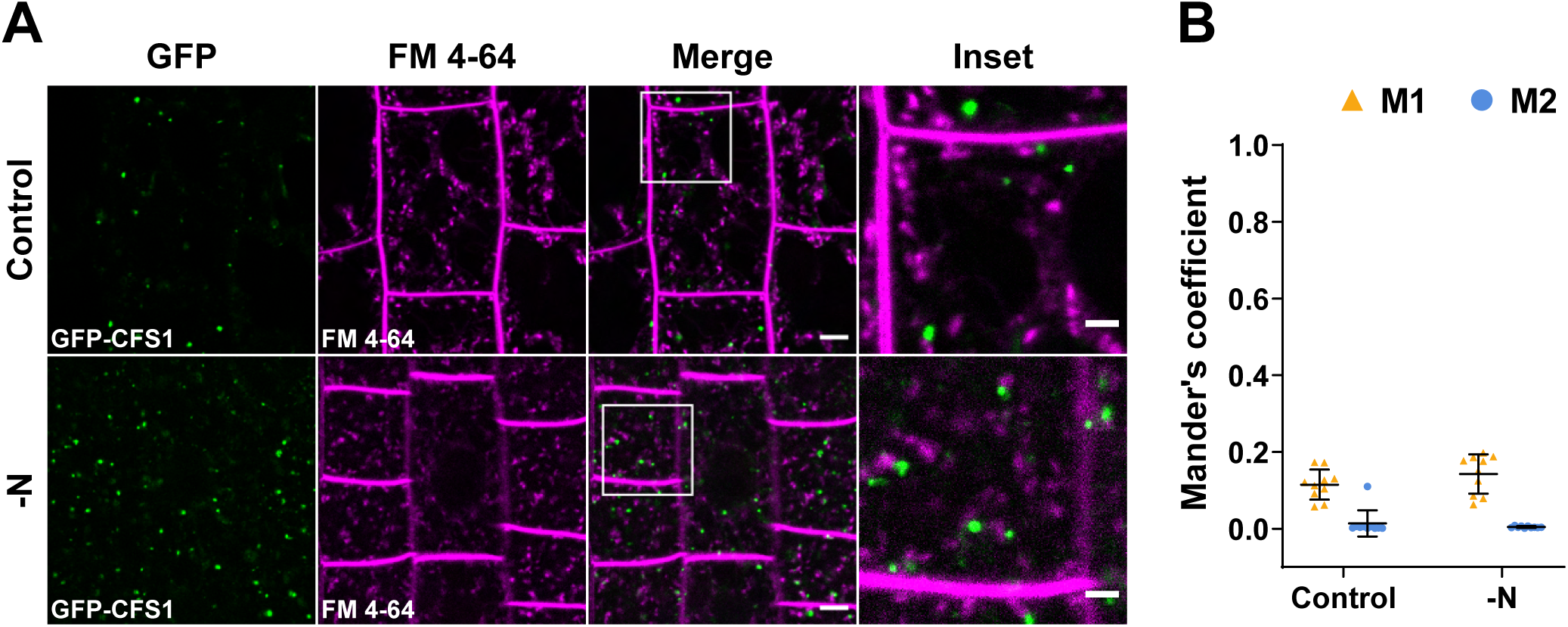
CFS1 does not colocalize with the endocytic marker dye FM 4-64. **(A)** Confocal microscopy images of Arabidopsis root epidermal cells expressing pUBQ::GFP-CFS1 and stained with FM 4-64. Five-day-old Arabidopsis seedlings were first incubated in either control or nitrogen- deficient (-N) 1/2 MS media for 4 hours and were then incubated in either control or nitrogen-deficient 1/2 MS media containing 4 μΜ FM 4-64 for 30 min before imaging. Area highlighted in the white-boxed region in the merge panel was further enlarged and presented in the inset panel. Scale bars, 5 μm. Inset scale bars, 2 μm **(B)** Quantification of confocal experiments in (A) showing the Mander’s colocalization coefficients between GFP-CFS1 and FM 4-64. M1, fraction of GFP-CFS1 signal that overlaps with FM 4-64 signal. M2, fraction of FM 4-64 signal that overlaps with GFP-CFS1 signal. Bars indicate the mean ± standard deviation of 10 biological replicates.

**Figure S5.**
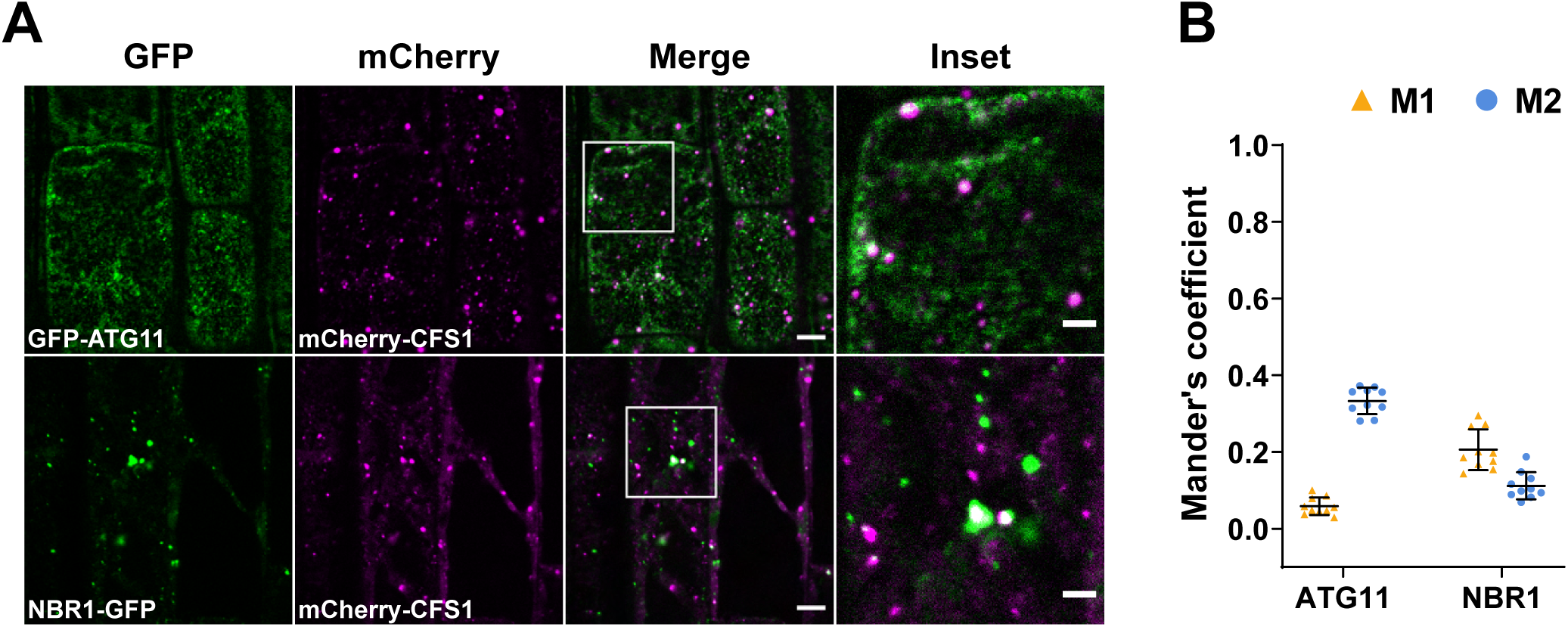
CFS1 colocalizes with the autophagosome marker proteins ATG11 and NBR1 during salt stress. **(A)** Confocal microscopy images of Arabidopsis root epidermal cells co-expressing pUBQ::mCherry-CFS1 and pUBQ::GFP-ATG11 or pNBR1::NBR1-GFP. Five-day-old Arabidopsis seedlings were incubated in 150 mM NaCl-containing 1/2 MS media for 1 h for autophagy induction before imaging. Area highlighted in the white-boxed region in the merge panel was further enlarged and presented in the inset panel. Scale bars, 5 μm. Inset scale bars, 2 μm. **(B)** Quantification of confocal experiments in (A) showing the Mander’s colocalization coefficients between mCherry-CFS1 and the GFP-fused marker ATG11 or NBR1. M1, fraction of GFP-fused marker signal that overlaps with mCherry-CFS1 signal. M2, fraction of mCherry-CFS1 signal that overlaps with GFP-fused marker signal. Bars indicate the mean ± standard deviation of 10 biological replicates.

**Figure S6.**
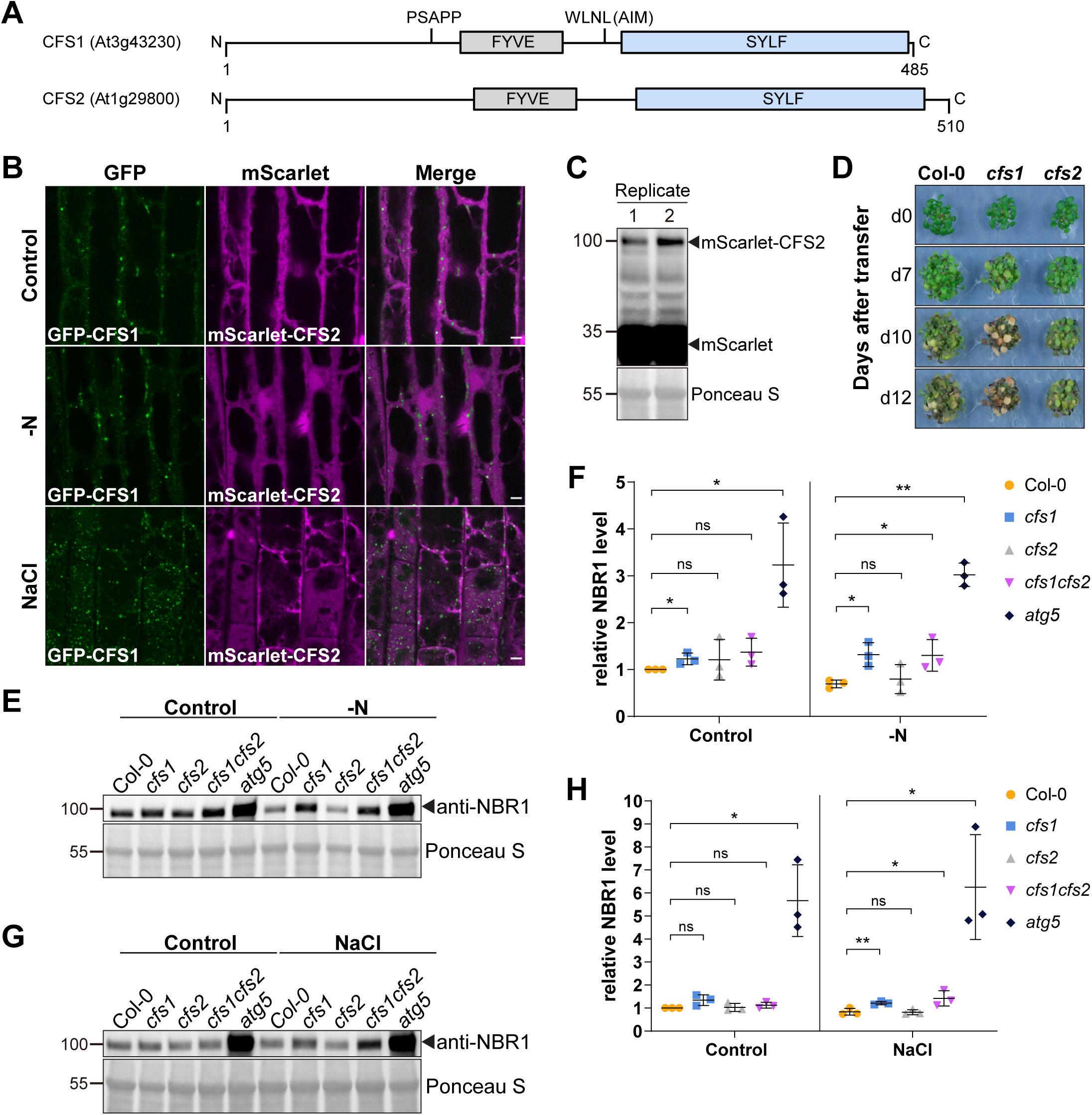
CFS2 does not play a role in autophagic flux. **(A)** Schematic diagrams showing the domain structures of Arabidopsis CFS1 and CFS2. **(B)** Confocal microscopy images of Arabidopsis root epidermal cells co-expressing pUBQ::GFP-CFS1 and pUBQ::mScarlet-CFS2. Five-day-old Arabidopsis seedlings were incubated in either control, nitrogen- deficient (-N) or 150 mM NaCl-containing 1/2 MS media before imaging. Note that no CFS2 puncta signals could be detected. Scale bars, 5 μm. **(C)** Western blot analysis of Arabidopsis seedlings expressing pUBQ::mScarlet-CFS2 used in (B). Total lysates were immunoblotted with anti-RFP antibodies. Images of two biological replicates are shown. **(D)** Phenotypic characterization of Arabidopsis *cfs1* and *cfs2* mutants upon nitrogen starvation. Twenty-five Arabidopsis seeds per genotype were first grown on 1/2 MS media plates (+1% plant agar) for 1 week and 7-day-old seedlings were subsequently transferred to nitrogen-deficient (-N) 1/2 MS media plates (+0.8% plant agar) and grown for 2 weeks. Plants were grown at 21°C under LEDs with 85 µM/m²/sec and a 14 h light/10 h dark photoperiod. d0 depicts the day of transfer. Brightness of pictures was enhanced ≤19% with Adobe Photoshop (2020). Representative images of 4 biological replicates are shown. **(E)** Western blots showing the endogenous NBR1 level in Col-0, *cf*s1, *cfs2*, *cfs1cfs2* or *atg5* under control or nitrogen starved (-N) conditions. Arabidopsis seeds were first grown in 1/2 MS media under continuous light for one week and 7-day-old seedlings were subsequently transferred to control or nitrogen-deficient 1/2 MS media for 12 h. Ten μl of total seedling extract was loaded and immunoblotted with anti-NBR1 antibodies. **(F)** Quantification of (E) showing the relative NBR1 level of Col-0, *cfs1*, *cfs2*, *cfs1cfs2* or *atg5* under control or nitrogen starved conditions. Values were calculated via normalization of protein bands to Ponceau S and to untreated (Control) Col-0 and shown as the mean ± standard deviation of 3 independent replicates. One- tailed and paired Student t-tests were performed to analyze the significance of the relative NBR1 level differences between Col-0 and each mutant. ns, not significant. *, p value < 0.05. **, p value < 0.01. **(G)** Western blot showing the endogenous NBR1 level in Col-0, *cfs*1, *cfs2*, *cfs1cfs2*, or *atg5* under control or salt stressed (NaCl) conditions. Arabidopsis seeds were first grown in 1/2 MS media under continuous light for one week and 7-day-old seedlings were subsequently transferred to control or 150 mM NaCl-containing 1/2 MS media for 16 h. Ten μl of total seedling extract was loaded and immunoblotted with anti-NBR1 antibodies. **(H)** Quantification of (G) showing the relative NBR1 level of Col-0, *cfs1*, *cfs2*, *cfs1cfs2*, or *atg5* under control or salt stress (NaCl) conditions. Values were calculated via normalization of protein bands to Ponceau S and to untreated (Control) Col-0 and shown as the mean ± standard deviation of 3 independent replicates. One-tailed and paired Student t-tests were performed to analyze the significance of the relative NBR1 level difference between Col-0 and each mutant. ns, not significant. *, p value < 0.05. **, p value < 0.01.

**Figure S7.**
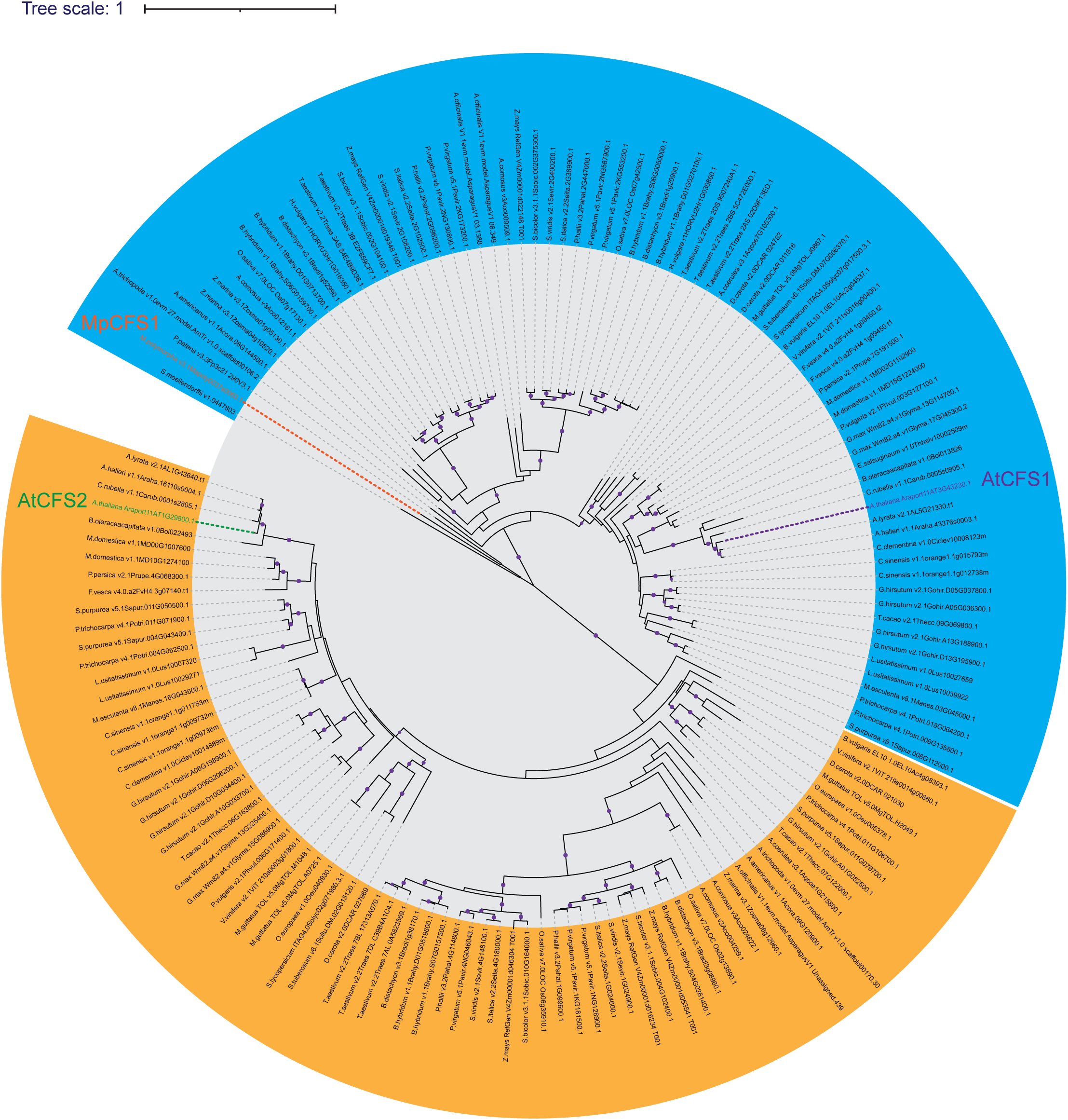
Maximum likelihood (ML) phylogenetic tree of homologous CFS1 and CFS2 from diverse plant species. Coding sequences of CFS1 and CFS2 homologs were obtained by using the BLAST tool against representative species of different plant lineages in Phytozome (Goodstein et al., 2012). The tree was inferred from a 2283-nt-long alignment by using the Maximum Likelihood method and Tamura-Nei model as implemented by MEGA X (Tamura and Nei, 1993; Kumar et al., 2018). 100 bootstrap method and a discrete Gamma distribution was used to model evolutionary rate differences among sites (5 categories (+G, parameter = 0.8072)) (Felsenstein, 1985). The tree is represented using Interactive Tree Of Life (iTOL) v4 (Letunic and Bork, 2019). Best-scoring ML tree (−98686.19) is shown with purple circles indicating bootstrap values above 80 on their respective clades. The scale bar indicates the evolutionary distance based on the nucleotide substitution rate. All CFS1 homologs are grouped in the blue region while all CFS2 homologs are grouped in the orange region. Genes that encode *Arabidopsis thaliana* CFS1 (AtCFS1), *A. thaliana* CFS2 (AtCFS2) and *Marchantia polymorpha* CFS1 (MpCFS1) are highlighted with purple, green and orange colors, respectively.

**Figure S8.**
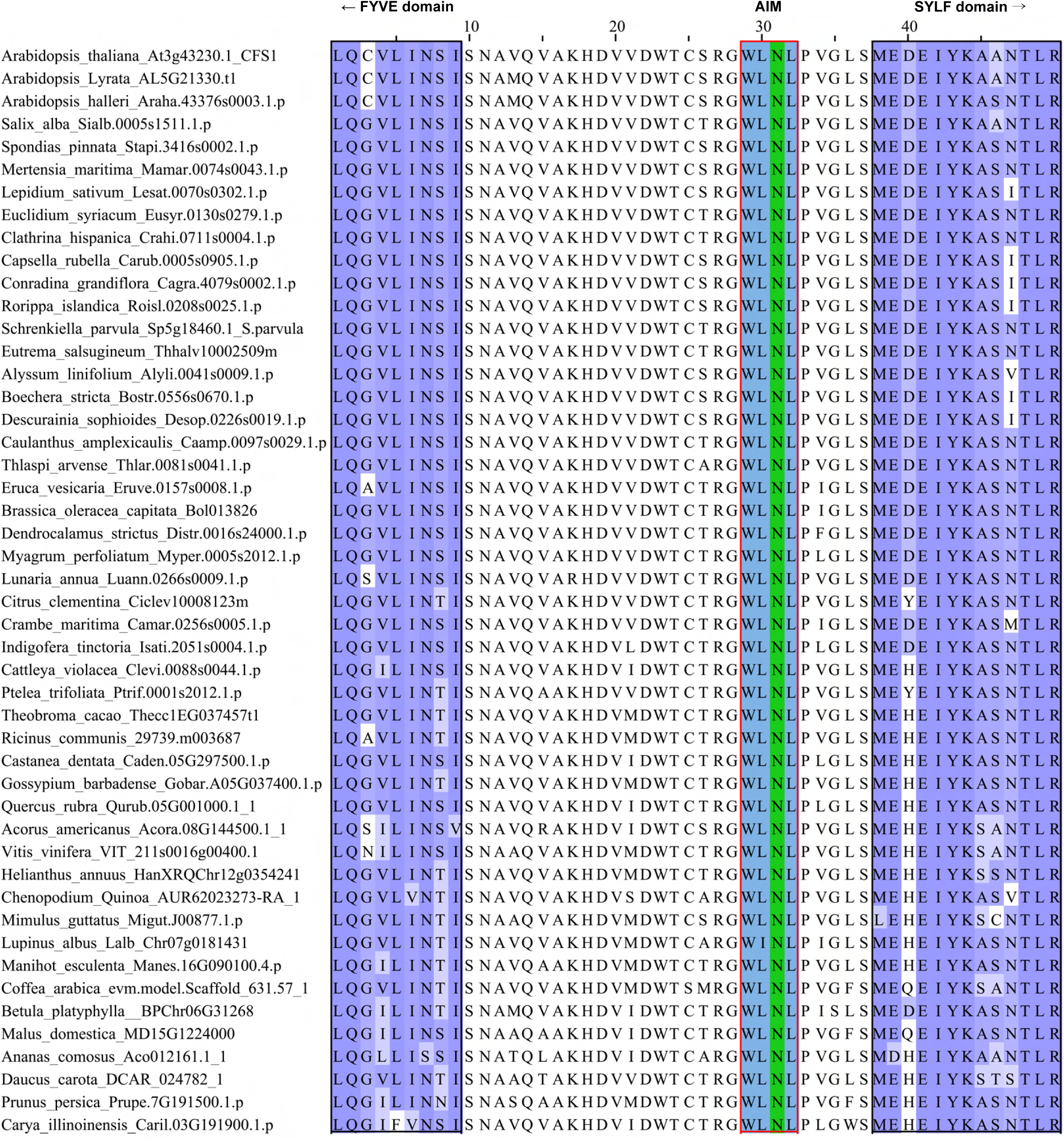
The AIM between the FYVE and SYLF domains of CFS1 is conserved in plants. Multiple sequencing alignments showing the conserved AIM (WLNL) between the FYVE and SYLF domains of CFS1. The regions that belong to FYVE or SYLF domains are labeled with black boxes.

**Figure S9.**
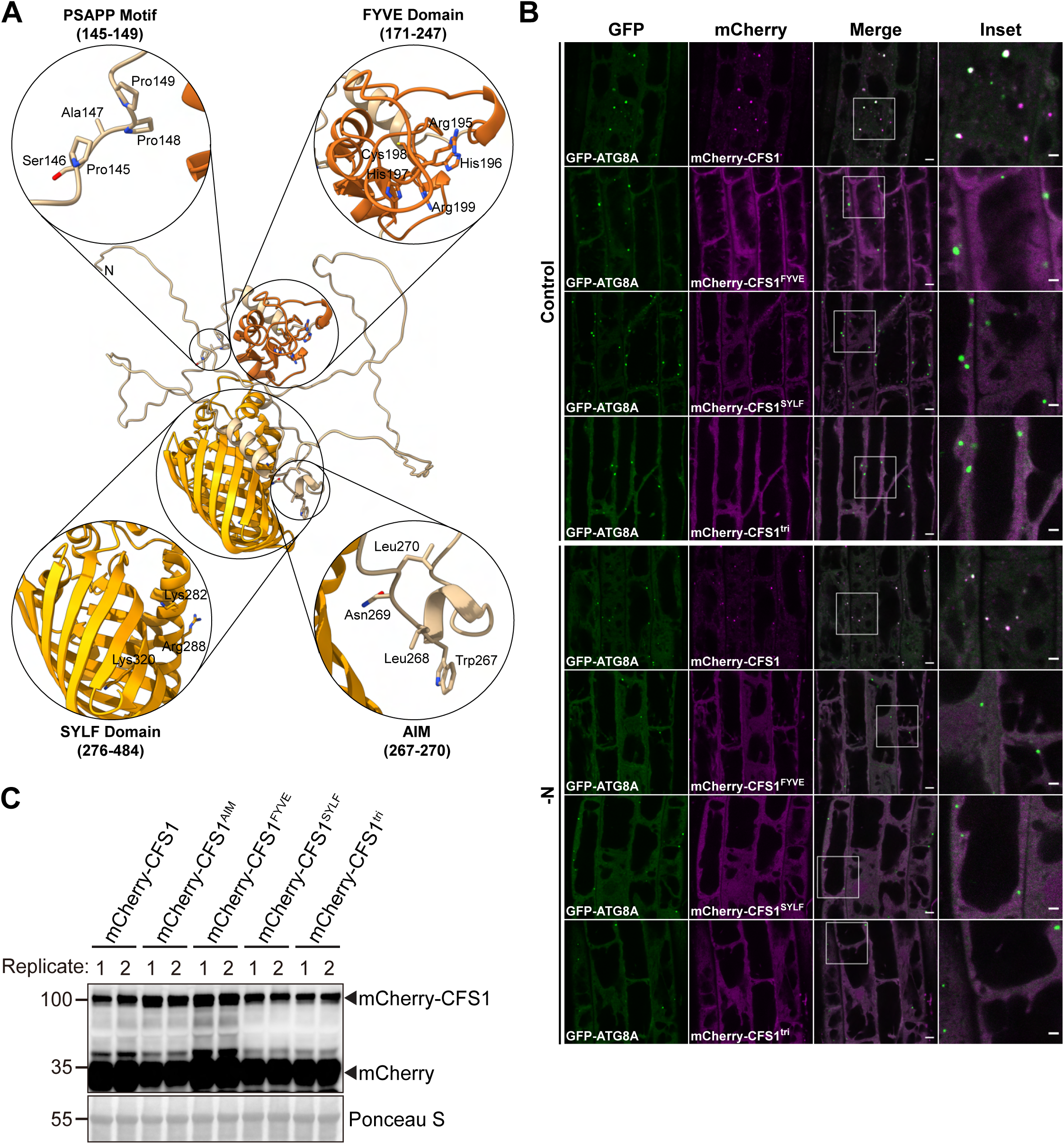
FYVE and SYLF domains are crucial for the autophagosome localization of CFS1. **(A)** Homology modelling and domain representation of CFS1. CFS1 structure is shown as ribbons and relevant motifs and domains are highlighted as zoom-in, with the side chains of relevant residues represented as stick. For clarity, the FYVE and SYLF domains of CFS1 are colored in brick red and orange, respectively. **(B)** Confocal microscopy images of *cfs1* mutants co-expressing pUBQ::GFP-ATG8A with pUBQ::mCherry-CFS1, pUBQ::mCherry-CFS1^FYVE^, pUBQ::mCherry-CFS1^SYLF^ or pUBQ::mCherry- CFS1^FYVE+SYLF+AIM^ (mCherry-CFS1^tri^). Five-day-old Arabidopsis seedlings were incubated in either control or nitrogen-deficient (-N) 1/2 MS media for 4 h before imaging. Area highlighted in the white-boxed region in the merge panel was further enlarged and presented in the inset panel. Scale bars, 5 μm. Inset scale bars, 2 μm. **(C)** Western blot showing the mCherry-CFS1 protein stability of the plant lines used in (B). Arabidopsis seeds were first grown in 1/2 MS media under continuous light for one week and 7-day-old seedlings were subsequently transferred to fresh 1/2 MS media for 16 h before protein extraction. Ten μg of total Arabidopsis seedling lysates was loaded and immunoblotted with anti-RFP antibodies. Images of 2 biological replicates are shown.

**Figure S10.**
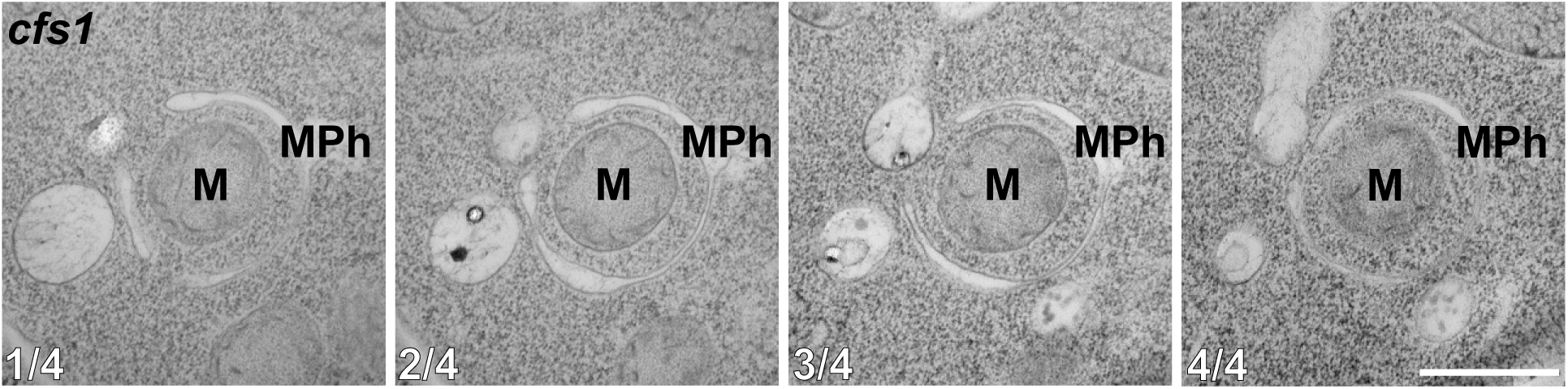
Mitophagosome formation is not disrupted in Arabidopsis *cfs1* mutants. Serial sections of transmission electron microscopy micrographs showing a mitophagosome engulfing a mitochondrion. Seven-day-old Arabidopsis *cfs1* seedlings were incubated in 1/2 MS media containing 50 μM DNP for 1 h before cryofixation. Scale bar, 500 nm. MPh, mitophagosome. M, mitochondrion.

**Figure S11.**
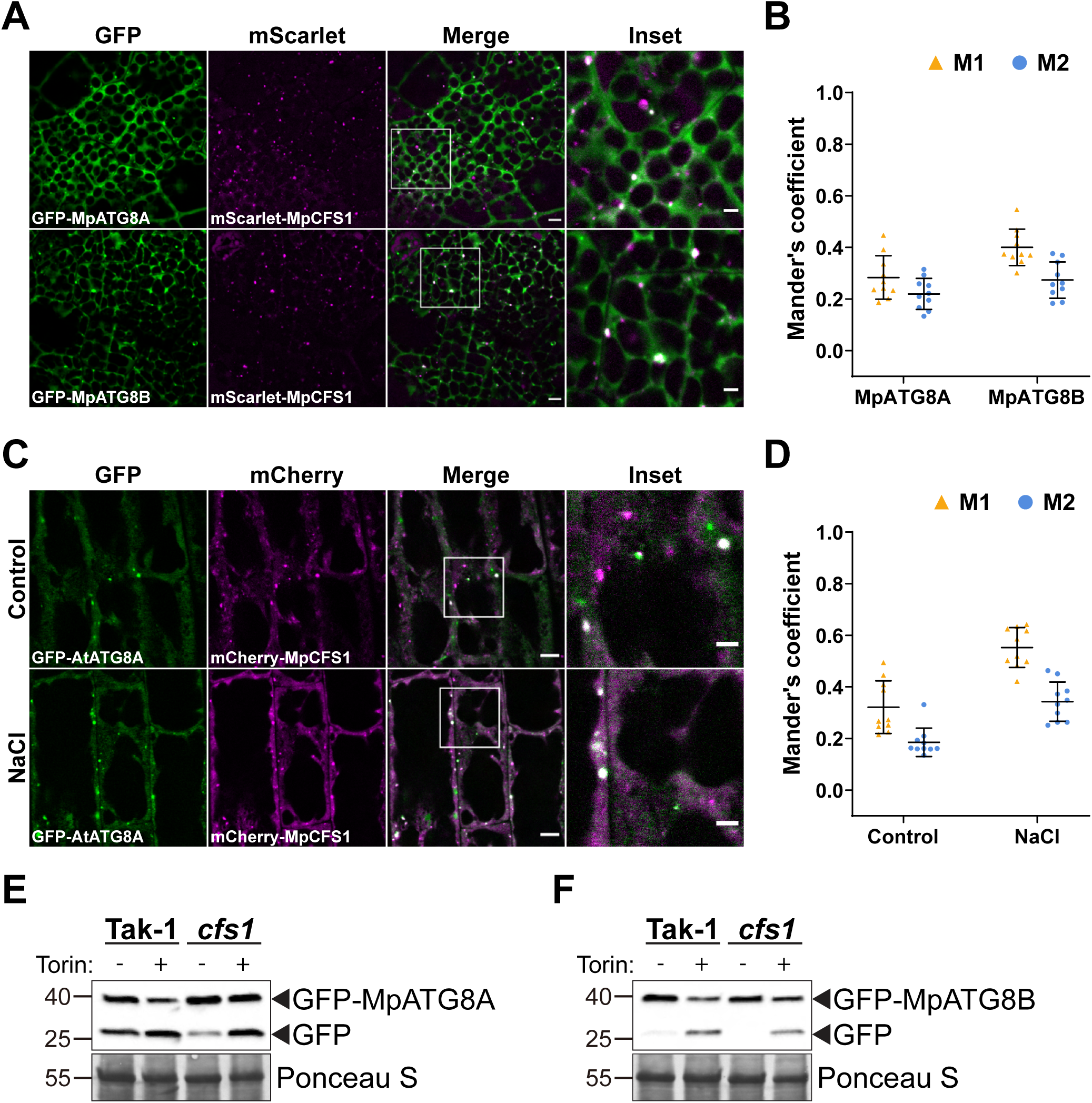
CFS1 function is conserved in *Marchantia polymorpha*. **(A)** Confocal microscopy images of *M. polymorpha* thallus cells co-expressing pEF1::mScarlet-MpCFS1 with pEF1::GFP-MpATG8A or pEF1::GFP-MpATG8B. Two-day-old thalli were incubated in 1/2 Gamborg B5 media before imaging. Area highlighted in the white-boxed region in the merge panel was further enlarged and presented in the inset panel. Scale bars, 5 μm. Inset scale bars, 2 μm. **(B)** Quantification of confocal experiments in (A) showing the Mander’s colocalization coefficients between mScarlet-MpCFS1 and GFP-fused MpATG8A or MpATG8B. M1, fraction of the GFP-fused MpATG8A or MpATG8B signals that overlaps with mScarlet-MpCFS1 signal. M2, fraction of mScarlet- MpCFS1 signal that overlaps with GFP-fused MpATG8A or MpATG8B signals. Bars indicate the mean ± standard deviation of 10 biological replicates. **(C)** Confocal microscopy images of Arabidopsis root epidermal cells co-expressing pUBQ::mCherry- MpCFS1 and pUBQ::GFP-AtATG8A. Five-day-old seedlings were incubated in either control or 150 mM NaCl-containing 1/2 MS media for 1 h before imaging. Scale bars, 5 μm. Inset scale bars, 2 μm. **(D)** Quantification of confocal experiments in (C) showing the Mander’s colocalization coefficients between mCherry-MpCFS1 and GFP-AtATG8A under control or salt stressed (NaCl) conditions. M1, fraction of the GFP-AtATG8A signal that overlaps with the mCherry-MpCFS1 signal. M2, fraction of the mCherry-MpCFS1 signal that overlaps with the GFP-AtATG8A signal. Bars indicate the mean ± standard deviation of 10 biological replicates. **(E)** GFP cleavage assay of pEF1::GFP-MpATG8A in *M. polymorpha* wild-type (Tak-1) or *cfs1* mutants. Ten- day-old propagules were treated with 12 μM Torin for 5 h before protein extraction. Fifteen μg of total protein extract was loaded and immunoblotted with anti-GFP antibodies. Representative images of 2 biological replicates are shown. **(F)** GFP cleavage assay of pEF1::GFP-MpATG8B in *M. polymorpha* wild-type (Tak-1) or *cfs1* mutants. Ten- day-old propagules were treated with 12 μM Torin for 5 h before protein extraction. Fifteen μg of total protein extract was loaded and was immunoblotted with anti-GFP antibodies. Representative images of 2 biological replicates are shown.

**Figure S12.**
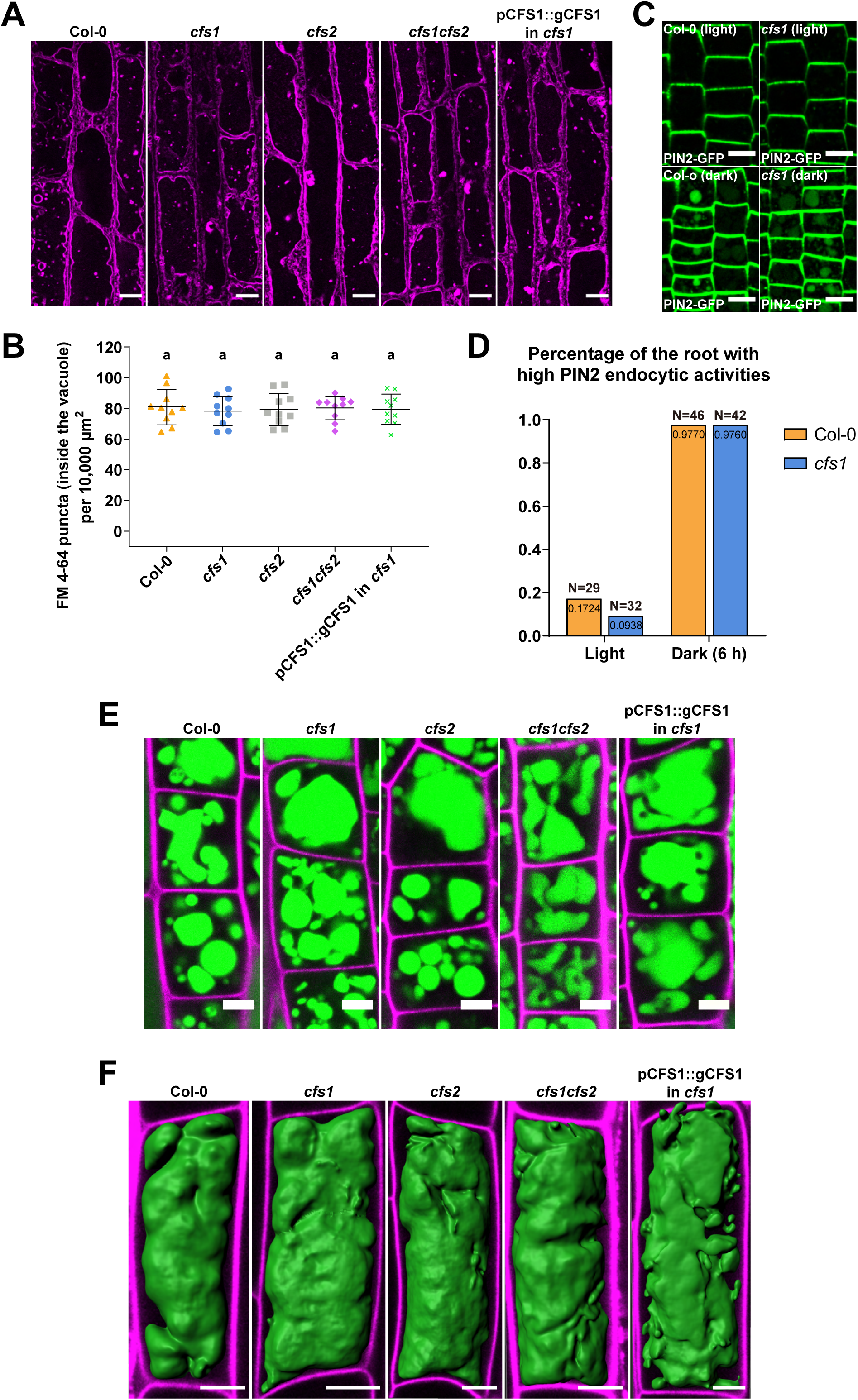
Endocytic trafficking is not affected in *cfs1* mutants. **(A)** Confocal microscopy images of Arabidopsis root epidermal cells of Col-0, *cfs1*, *cfs2*, *cfs1cfs2*, and *cfs1* complemented with pCFS1::gCFS1(pCFS1::gCFS1 in *cfs1*). Five-day-old Arabidopsis seedlings were first incubated in 4 μΜ FM 4-64-containing 1/2 MS media for 30 min and then transferred to 1 μΜ concanamycin A-containing 1/2 MS media for 2 hours before imaging. Scale bars, 10 μm. **(B)** Quantification of the FM 4-64 stained puncta inside the vacuole per normalized area (10,000 μm^2^) of the cells imaged in (A). Bars indicate the mean ± standard deviation of 10 biological replicates. One-way ANOVA tests were performed to analyze the differences of the number of FM 4-64 stained puncta between each group. Tukey’s multiple comparison tests were used for multiple comparisons. Family-wise significance and confidence level, 0.05 (95% confidence interval). **(C)** Representative microscopy images showing PIN2 endocytosis in the epidermal cells in the root tip meristem region of Col-0 and *cfs1* under light or 6 h dark conditions. Five-day-old Arabidopsis seedlings expressing pPIN2::PIN2-GFP were grown on 1/2 MS media plates (+1% plant agar) under light or 6 h dark conditions before imaging. Scale bars, 10 μm. **(D)** Quantification of PIN2 endocytic activities in Col-0 and *cfs1* shown in (C). The Arabidopsis seedlings with at least 5 root epidermal cells that contained visible PIN2-GFP in the vacuole were considered as high PIN2 endocytic activities. The percentage of Col-0 and *cfs1* with high PIN2 endocytic activities under light or 6 h dark conditions are shown in the graph. Numbers inside the bars represent the exact value (4 decimals) of each bar. N represents the total number of the Col-0 or *cfs1* seedlings used for imaging and quantification in 3 independent experiments. **(E)** Confocal microscopy images showing the BCECF-AM-stained root epidermal cells in the meristem region of Col-0, *cfs1*, *cfs2*, *cfs1cfs2*, or pCFS1::gCFS1 in *cfs1*. Five-day-old Arabidopsis seedlings were incubated in 1/2 MS media containing 5 μΜ BCECF-AM for 30 min before imaging. Samples were mounted on slides with 0.002 mg/ml propidium iodide. Green signals indicate the BCECF-AM stained vacuole. Magenta signals indicate the propidium iodide-stained cell wall. Scale bars, 5 μm. **(F)** Three-dimensional images showing the vacuolar structure of the BCECF-AM-stained root epidermal cells in the transition region of Col-0, *cfs1*, *cfs2*, *cfs1cfs2*, or pCFS1::gCFS1 in *cfs1*. Five-day-old Arabidopsis seedlings were incubated in 1/2 MS media containing 5 μΜ BCECF-AM for 30 min before imaging. Samples were mounted on slides with 0.002 mg/ml propidium iodide. Green signals indicate the BCECF- AM stained vacuole. Magenta signals indicate the propidium iodide-stained cell wall. Scale bars, 10 μm.

**Figure S13.**
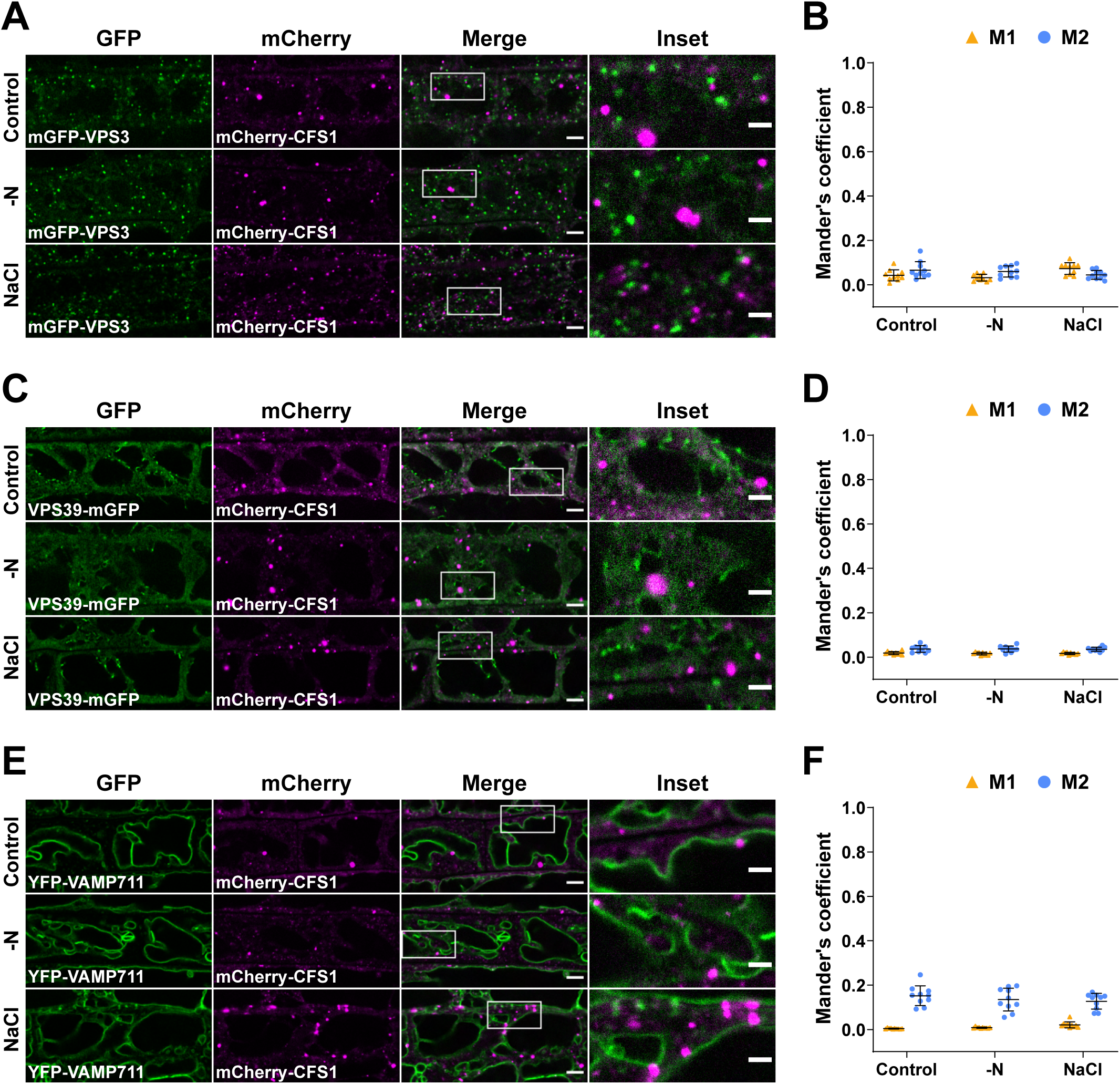
CFS1 does not colocalize with the CORVET complex component VPS3, the HOPS complex component VPS39, or the tonoplast marker VAMP711. **(A)** Confocal microscopy images of Arabidopsis root epidermal cells co-expressing pVPS3::mGFP-VPS3 with pUBQ::mCherry-CFS1. Five-day-old Arabidopsis seedlings were incubated in either control, nitrogen- deficient (-N) or 150 mM NaCl-containing 1/2 MS media before imaging. Area highlighted in the white- boxed region in the merge panel was further enlarged and presented in the inset panel. Scale bars, 5 μm. Inset scale bars, 2 μm. **(B)** Quantification of confocal experiments in (A) showing the Mander’s colocalization coefficients between mCherry-CFS1 and mGFP-VPS3. M1, fraction of mGFP-VPS3 signal that overlapped with the mCherry-CFS1 signal. M2, fraction of mCherry-CFS1 signal that overlapped with the mGFP-Vps3 signal. Bars indicate the mean ± standard deviation of 10 biological replicates. **(C)** Confocal microscopy images of Arabidopsis root epidermal cells co-expressing pVPS39::VPS39-mGFP with pUBQ::mCherry-CFS1. Five-day-old Arabidopsis seedlings were incubated in either control, nitrogen- deficient (-N) or 150 mM NaCl-containing 1/2 MS media before imaging. Area highlighted in the white- boxed region in the merge panel was further enlarged and presented in the inset panel. Scale bars, 5 μm. Inset scale bars, 2 μm. **(D)** Quantification of confocal experiments in (C) showing the Mander’s colocalization coefficients between mCherry-CFS1 and VPS39-mGFP. M1, fraction of VPS39-mGFP signal that overlapped with the mCherry-CFS1 signal. M2, fraction of mCherry-CFS1 signal that overlapped with the VPS39-mGFP signal. Bars indicate the mean ± standard deviation of 10 biological replicates. **(E)** Confocal microscopy images of Arabidopsis root epidermal cells co-expressing pUBQ::YFP- VAMP711 with pUBQ::mCherry-CFS1. Five-day-old Arabidopsis seedlings were incubated in either control, nitrogen-deficient (-N) or 150 mM NaCl-containing 1/2 MS media before imaging. Area highlighted in the white-boxed region in the merge panel was further enlarged and presented in the inset panel. Scale bars, 5 μm. Inset scale bars, 2 μm. **(F)** Quantification of confocal experiments in (E) showing the Mander’s colocalization coefficients between mCherry-CFS1 and YFP-VAMP711. M1, fraction of YFP-VAMP711 signal that overlapped with the mCherry-CFS1 signal. M2, fraction of mCherry-CFS1 signal that overlapped with the YFP-VAMP711 signal. Bars indicate the mean ± standard deviation of 10 biological replicates.

**Figure S14.**
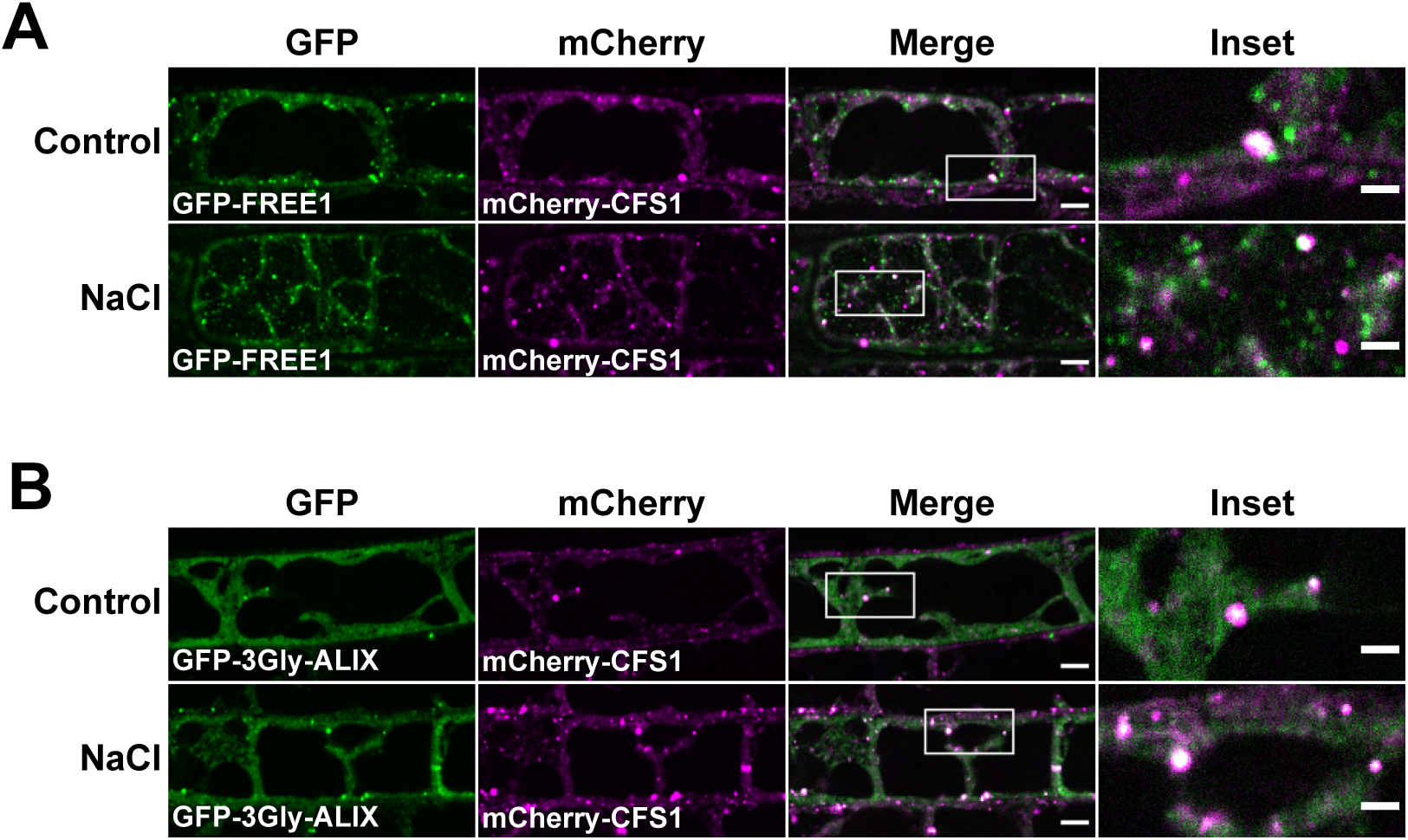
CFS1 partially colocalizes with FREE1 and ALIX. **(A)** Confocal microscopy images of Arabidopsis root epidermal cells co-expressing p35S::GFP-FREE1 with pUBQ::mCherry-CFS1. Five-day-old Arabidopsis seedlings were incubated in either control or 150 mM NaCl-containing 1/2 MS media for 1 hour before imaging. Area highlighted in the white-boxed region in the merge panel was further enlarged and presented in the inset panel. Scale bars, 5 μm. Inset scale bars, 2 μm. **(B)** Confocal microscopy images of Arabidopsis root epidermal cells co-expressing pALIX::GFP-3Gly- ALIX with pUBQ::mCherry-CFS1. Five-day-old Arabidopsis seedlings were incubated in either control or 150 mM NaCl-containing 1/2 MS media for 1 hour before imaging. Area highlighted in the white-boxed region in the merge panel was further enlarged and presented in the inset panel. Scale bars, 5 μm. Inset scale bars, 2 μm.

**Figure S15.**
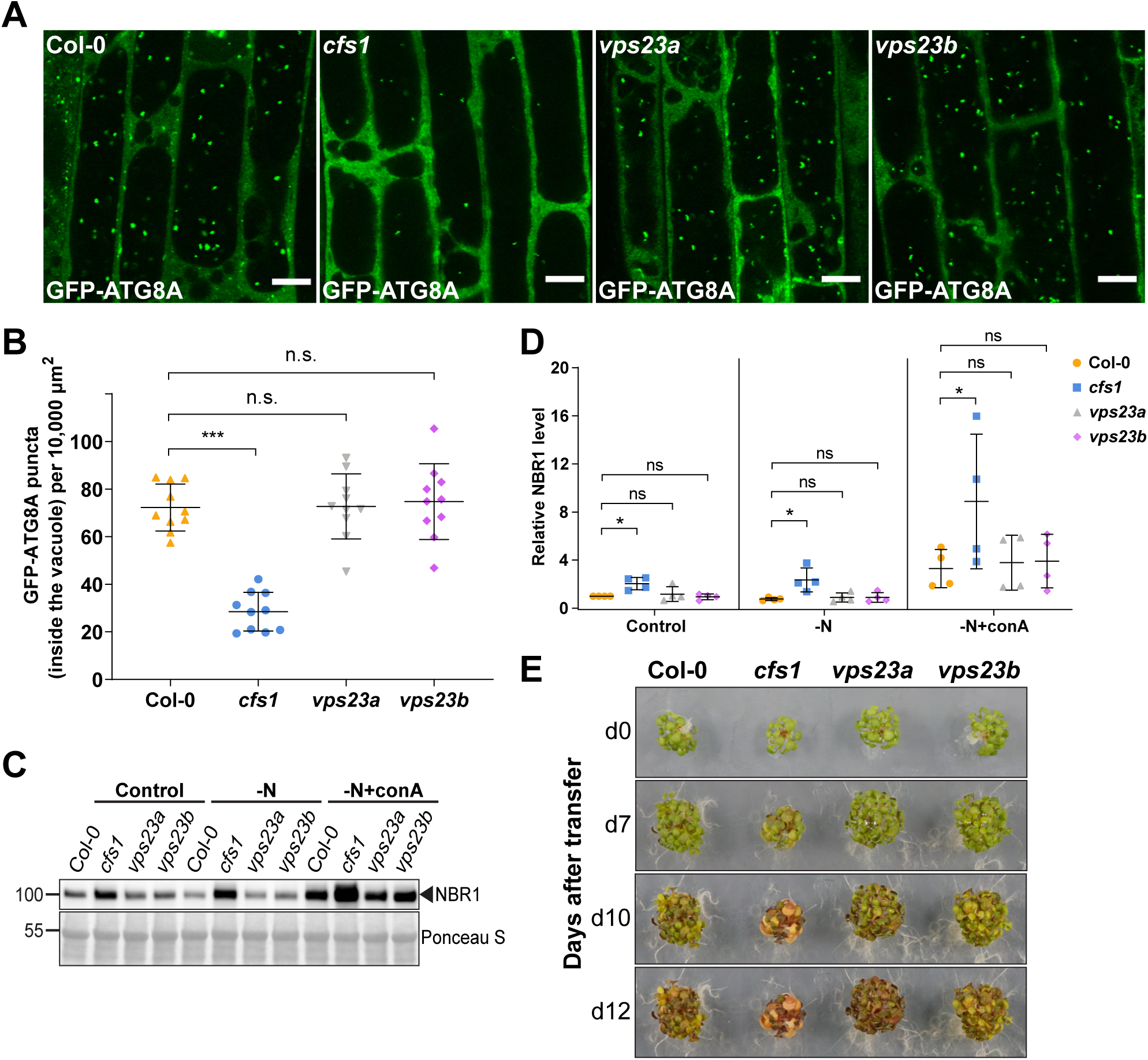
*vps23a* and *vps23b* single mutants do not have defects in autophagic flux. **(A)** Confocal microscopy images of root epidermal cells of Col-0, *cfs1*, *vps23a*, or *vps23b* seedlings expressing pUBQ::GFP-ATG8A under NaCl + concanamycin A (conA) treatment. Five-day-old Arabidopsis seedlings were incubated in 1/2 MS media containing 90 mM NaCl and 1 μΜ conA for 2 hours before imaging. Scale bars, 10 μm. **(B)** Quantification of the number of GFP-ATG8A puncta inside the vacuole per normalized area (10,000 μm^2^) of the cells imaged in (A). Bars indicate the mean ± standard deviation of 10 biological replicates. Two-tailed and unpaired Student t-tests were performed to analyze the significance of GFP-ATG8A puncta density differences between Col-0 and *cfs1*, Col-0 and *vps23a*, or Col-0 and *vps23b*. ns, not significant. ***, p value < 0.001. **(C)** Western blot showing the endogenous NBR1 level in Col-0, *cfs1*, *vps23a* or *vps23b* under control or nitrogen-deficient ± conA conditions. Arabidopsis seeds were first grown in 1/2 MS media under continuous light for one week and 7-day-old seedlings were subsequently transferred to 1/2 MS media (Control), nitrogen-deficient 1/2 MS media (-N) or nitrogen-deficient 1/2 MS media containing 1 µM conA (-N+conA) for 12 h. Ten μg of total protein extract was loaded and immunoblotted with anti-NBR1 antibodies. **(D)** Quantification of the relative NBR1 level in (C) compared to untreated (control) Col-0. Values were normalized to untreated (Control) Col-0 and calculated via normalization of protein bands to Ponceau S and shown as the mean ± standard deviation of 4 independent replicates. One-tailed and paired Student t-tests were performed to analyze the significance of the relative NBR1 level difference. ns, not significant. *, p < 0.05. **(E)** Phenotypic characterization of Arabidopsis *cfs1*, *vps23a* and *vps23b* mutants during nitrogen starvation. Twenty-five seeds per genotype were grown on 1/2 MS media plates (+1% plant agar) for 1 week and 7- day-old seedlings were subsequently transferred to nitrogen-deficient 1/2 MS media plates (+0.8% plant agar) and grown for 2 weeks. Plants were grown at 21°C under LEDs with 85 µM/m²/sec and a 14 h light/10 h dark photoperiod. d0 depicts the day of transfer. Brightness of pictures was enhanced ≤30% with Adobe Photoshop (2020). Representative images of 4 independent replicates are shown.

**Figure S16.**
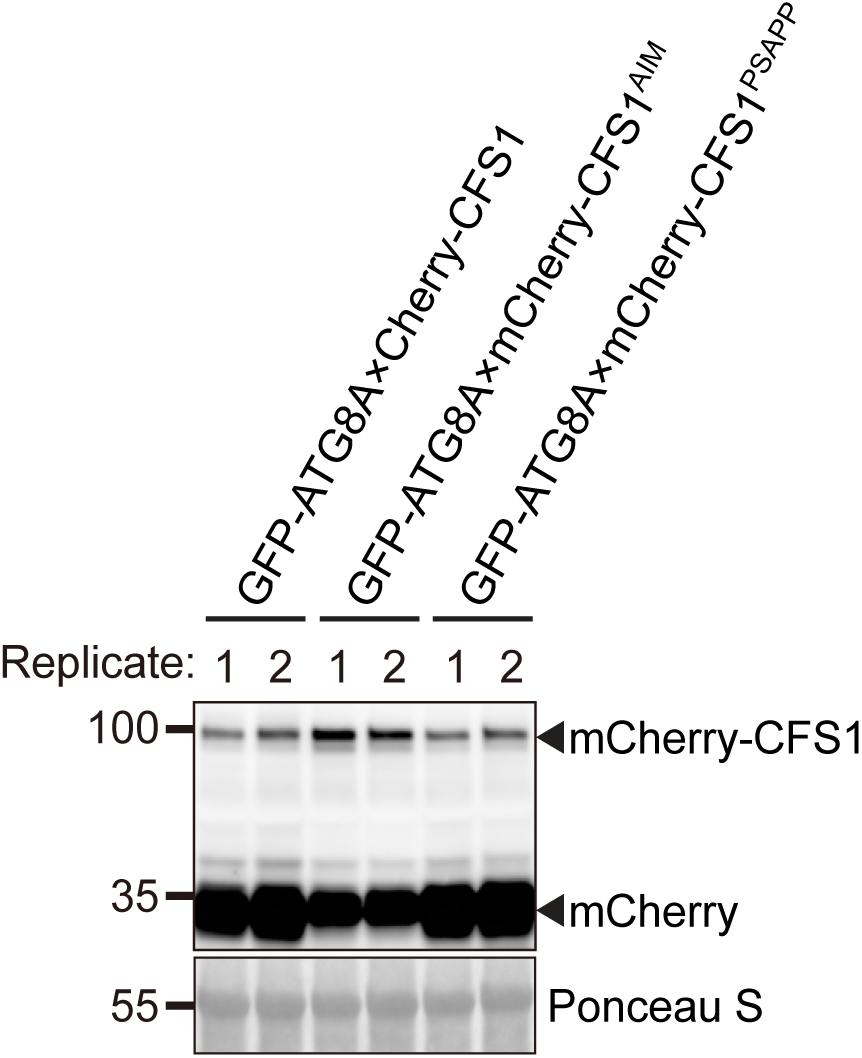
mCherry-CFS1 mutants are stably expressed in transgenic Arabidopsis lines. Western blot showing the steady expression of mCherry-CFS1, mCherry-CFS1^AIM^ or mCherry-CFS1^PSAPP^ in the Arabidopsis lines used in Figure 5G and 5I. Ten μg of total plant lysates of Arabidopsis *cfs1* mutants co-expressing pUBQ::GFP-ATG8A and pUBQ::mCherry-CFS1, pUBQ::mCherry-CFS1^AIM^ or pUBQ::mCherry-CFS1^PSAPP^ were immunoblotted with anti-RFP antibodies. Images of 2 biological replicates are shown.

**Supplementary video 1.** Time-lapse video showing that mCherry-CFS1 moves together with GFP-ATG8A in Arabidopsis root epidermal cells. Five-day-old Arabidopsis seedlings co- expressing pUBQ::GFP-ATG8A and pUBQ::mCherry-CFS1 were incubated in 150 mM NaCl- containing 1/2 MS media for 1 h for autophagy induction before imaging. Total imaging time, 60 seconds. Interval, 1 second. Scale bar, 10 μm.

**Supplementary video 2.** Time-lapse video showing that mCherry-CFS1 moves together with NBR1-GFP in Arabidopsis root epidermal cells. Five-day-old Arabidopsis seedlings co-expressing pNBR1::NBR1-GFP and pUBQ::mCherry-CFS1 were incubated in 150 mM NaCl-containing 1/2 MS media for 1 h for autophagy induction before imaging. Total imaging time, 60 seconds. Interval, 1 second. Scale bar, 10 μm.

**Supplementary video 3.** Time-lapse video showing the partial colocalization and the associated movement between GFP-CFS1 and VPS23A-TagRFP in Arabidopsis root epidermal cells. Five- day-old Arabidopsis seedlings co-expressing pUBQ::GFP-CFS1 and pVPS23A::VPS23A-TagRFP were incubated in 150 mM NaCl-containing 1/2 MS media for 1 h for autophagy induction before imaging. Total imaging time, 60 seconds. Interval, 1 second. Scale bar, 10 μm.

**Supplementary video 4.** Time-lapse video showing cell-to-cell movement of NBR1- GFP/mCherry-CFS1 punctum in Arabidopsis root cells. Five-day-old Arabidopsis seedlings co- expressing pNBR1::NBR1-GFP and pUBQ::mCherry-CFS1 were incubated in 150 mM NaCl-containing 1/2 MS media for 1 h for autophagy induction before imaging. Total imaging time, 60 seconds. Interval, 1 second. Scale bar, 5 μm.

## References

Allan, B.B., B.D. Moyer, and W.E. Balch. 2000. Rab1 recruitment of p115 into a cis-SNARE complex: programming budding COPII vesicles for fusion. Science. 289:444–448.

Aniento, F., V. Sánchez de Medina Hernández, Y. Dagdas, M. Rojas-Pierce, and E. Russinova. 2022. Molecular mechanisms of endomembrane trafficking in plants. Plant Cell. 34:146–173.

Bassham, D.C. 2015. Methods for analysis of autophagy in plants. Methods. 75:181–188.

Birgisdottir, Å.B., T. Lamark, and T. Johansen. 2013. The LIR motif – crucial for selective autophagy. Journal of Cell Science. 126:3237–3247. doi:10.1242/jcs.126128.

Borner, G.H.H., D.J. Sherrier, T. Weimar, L.V. Michaelson, N.D. Hawkins, A. Macaskill, J.A. Napier, M.H. Beale, K.S. Lilley, and P. Dupree. 2005. Analysis of detergent-resistant membranes in Arabidopsis. Evidence for plasma membrane lipid rafts. Plant Physiol. 137:104–116.

Chang, C., L.E. Jensen, and J.H. Hurley. 2021. Autophagosome biogenesis comes out of the black box. Nat. Cell Biol. doi:10.1038/s41556-021-00669-y.

Clough, S.J., and A.F. Bent. 1998. Floral dip: a simplified method forAgrobacterium-mediated transformation ofArabidopsis thaliana. Plant J. 16:735–743.

Cui, Y., W. Cao, Y. He, Q. Zhao, M. Wakazaki, X. Zhuang, J. Gao, Y. Zeng, C. Gao, Y. Ding, H.Y. Wong, W.S. Wong, H.K. Lam, P. Wang, T. Ueda, M. Rojas-Pierce, K. Toyooka, B.-H. Kang, and L. Jiang. 2019. A whole-cell electron tomography model of vacuole biogenesis in Arabidopsis root cells. Nat Plants. 5:95–105.

Cui, Y., Q. Zhao, S. Hu, and L. Jiang. 2020. Vacuole Biogenesis in Plants: How Many Vacuoles, How Many Models? Trends Plant Sci. 25:538–548.

Dikic, I. 2017. Proteasomal and Autophagic Degradation Systems. Annu. Rev. Biochem. 86:193–224.

Felsenstein, J. 1985. CONFIDENCE LIMITS ON PHYLOGENIES: AN APPROACH USING THE BOOTSTRAP. Evolution. 39:783–791.

Gao, C., X. Zhuang, Y. Cui, X. Fu, Y. He, Q. Zhao, Y. Zeng, J. Shen, M. Luo, and L. Jiang. 2015. Dual roles of an Arabidopsis ESCRT component FREE1 in regulating vacuolar protein transport and autophagic degradation. Proc. Natl. Acad. Sci. U. S. A. 112:1886–1891.

Geldner, N., V. Dénervaud-Tendon, D.L. Hyman, U. Mayer, Y.-D. Stierhof, and J. Chory. 2009. Rapid, combinatorial analysis of membrane compartments in intact plants with a multicolor marker set. The Plant Journal. 59:169–178. doi:10.1111/j.1365-313x.2009.03851.x.

Goodstein, D.M., S. Shu, R. Howson, R. Neupane, R.D. Hayes, J. Fazo, T. Mitros, W. Dirks, U. Hellsten, N. Putnam, and D.S. Rokhsar. 2012. Phytozome: a comparative platform for green plant genomics. Nucleic Acids Res. 40:D1178–86.

Ishizaki, K., R. Nishihama, M. Ueda, K. Inoue, S. Ishida, Y. Nishimura, T. Shikanai, and T. Kohchi. 2015. Development of Gateway Binary Vector Series with Four Different Selection Markers for the Liverwort Marchantia polymorpha. PLoS One. 10:e0138876.

Jia, M., X. Liu, H. Xue, Y. Wu, L. Shi, R. Wang, Y. Chen, N. Xu, J. Zhao, J. Shao, Y. Qi, L. An, J. Sheen, and F. Yu. 2019. Noncanonical ATG8-ABS3 interaction controls senescence in plants. Nat Plants. 5:212– 224.

Jumper, J., and D. Hassabis. 2022. Protein structure predictions to atomic accuracy with AlphaFold. Nat. Methods. 19:11–12.

Kalinowska, K., M.-K. Nagel, K. Goodman, L. Cuyas, F. Anzenberger, A. Alkofer, J. Paz-Ares, P. Braun, V. Rubio, M.S. Otegui, and E. Isono. 2015. Arabidopsis ALIX is required for the endosomal localization of the deubiquitinating enzyme AMSH3. Proc. Natl. Acad. Sci. U. S. A. 112:E5543–51.

Kang, B.-H. 2010. Electron microscopy and high-pressure freezing of Arabidopsis. Methods Cell Biol. 96:259– 283.

Kim, J.H., H.N. Lee, X. Huang, H. Jung, M.S. Otegui, F. Li, and T. Chung. 2022. FYVE2, a phosphatidylinositol 3-phosphate effector, interacts with the COPII machinery to control autophagosome formation in Arabidopsis. Plant Cell. 34:351–373.

Kleine-Vehn, J., J. Leitner, M. Zwiewka, M. Sauer, L. Abas, C. Luschnig, and J. Friml. 2008. Differential degradation of PIN2 auxin efflux carrier by retromer-dependent vacuolar targeting. Proc. Natl. Acad. Sci. U. S. A. 105:17812–17817.

Krebs, M., D. Beyhl, E. Görlich, K.A.S. Al-Rasheid, I. Marten, Y.-D. Stierhof, R. Hedrich, and K. Schumacher. 2010. Arabidopsis V-ATPase activity at the tonoplast is required for efficient nutrient storage but not for sodium accumulation. Proc. Natl. Acad. Sci. U. S. A. 107:3251–3256.

Krüger, F., and K. Schumacher. 2018. Pumping up the volume- vacuole biogenesis in Arabidopsis thaliana. In Seminars in cell & developmental biology. Elsevier. 106–112.

Kubota, A., K. Ishizaki, M. Hosaka, and T. Kohchi. 2013. Efficient Agrobacterium-mediated transformation of the liverwort Marchantia polymorpha using regenerating thalli. Biosci. Biotechnol. Biochem. 77:167–172.

Kumar, S., G. Stecher, M. Li, C. Knyaz, and K. Tamura. 2018. MEGA X: Molecular Evolutionary Genetics Analysis across computing platforms. Mol. Biol. Evol. 35:1547–1549.

Kurusu, T., T. Koyano, S. Hanamata, T. Kubo, Y. Noguchi, C. Yagi, N. Nagata, T. Yamamoto, T. Ohnishi, Y. Okazaki, N. Kitahata, D. Ando, M. Ishikawa, S. Wada, A. Miyao, H. Hirochika, H. Shimada, A. Makino, K. Saito, H. Ishida, T. Kinoshita, N. Kurata, and K. Kuchitsu. 2014. OsATG7 is required for autophagy-dependent lipid metabolism in rice postmeiotic anther development. Autophagy. 10:878–888.

LaMontagne, E.D., C.A. Collins, S.C. Peck, and A. Heese. 2016. Isolation of Microsomal Membrane Proteins from Arabidopsis thaliana. Curr Protoc Plant Biol. 1:217–234.

Lampropoulos, A., Z. Sutikovic, C. Wenzl, I. Maegele, J.U. Lohmann, and J. Forner. 2013. GreenGate---a novel, versatile, and efficient cloning system for plant transgenesis. PLoS One. 8:e83043.

Letunic, I., and P. Bork. 2019. Interactive Tree Of Life (iTOL) v4: recent updates and new developments. Nucleic Acids Res. 47:W256–W259.

Liu, Y., and D.C. Bassham. 2010. TOR is a negative regulator of autophagy in Arabidopsis thaliana. PLoS One. 5:e11883.

Liu, Y., Y. Xiong, and D.C. Bassham. 2009. Autophagy is required for tolerance of drought and salt stress in plants. Autophagy. 5:954–963.

Ma, J., Z. Liang, J. Zhao, P. Wang, W. Ma, K.K. Mai, J.A. Fernandez Andrade, Y. Zeng, N. Grujic, L. Jiang, Y. Dagdas, and B.-H. Kang. 2021. Friendly mediates membrane depolarization-induced mitophagy in Arabidopsis. Current Biology. 31:1931–1944.e4. doi:10.1016/j.cub.2021.02.034.

Marshall, R.S., and R.D. Vierstra. 2018. Autophagy: The Master of Bulk and Selective Recycling. Annu. Rev. Plant Biol. 69:173–208.

McLoughlin, F., R.C. Augustine, R.S. Marshall, F. Li, L.D. Kirkpatrick, M.S. Otegui, and R.D. Vierstra. 2018. Maize multi-omics reveal roles for autophagic recycling in proteome remodelling and lipid turnover. Nat Plants. 4:1056–1070.

McLoughlin, F., R.S. Marshall, X. Ding, E.C. Chatt, L.D. Kirkpatrick, R.C. Augustine, F. Li, M.S. Otegui, and R.D. Vierstra. 2020. Autophagy Plays Prominent Roles in Amino Acid, Nucleotide, and Carbohydrate Metabolism during Fixed-Carbon Starvation in Maize. Plant Cell. 32:2699–2724.

Melia, T.J., A.H. Lystad, and A. Simonsen. 2020. Autophagosome biogenesis: From membrane growth to closure. J. Cell Biol. 219. doi:10.1083/jcb.202002085.

Nagel, M.-K., K. Kalinowska, K. Vogel, G.D. Reynolds, Z. Wu, F. Anzenberger, M. Ichikawa, C. Tsutsumi, M.H. Sato, B. Kuster, S.Y. Bednarek, and E. Isono. 2017. Arabidopsis SH3P2 is an ubiquitin-binding protein that functions together with ESCRT-I and the deubiquitylating enzyme AMSH3. Proc. Natl. Acad. Sci. U. S. A. 114:E7197–E7204.

Nakatogawa, H. 2020. Mechanisms governing autophagosome biogenesis. Nat. Rev. Mol. Cell Biol. 21:439–458.

Norizuki, T., N. Minamino, H. Tsukaya, and T. Ueda. 2021. Bryophyte spermiogenesis occurs through multimode autophagic and nonautophagic degradation. bioRxiv. 2021.08.17.456730.

Oti, O.O. 2013. Hub and Spoke Network Design for the Inbound Supply Chain. Massachusetts Institute of Technology, Sloan School of Management; and, (S.M.)--Massachusetts Institute of Technology, Engineering Systems Division; in conjunction with the Leaders for Global Operations Program at MIT. 74 pp.

Perez-Riverol, Y., A. Csordas, J. Bai, M. Bernal-Llinares, S. Hewapathirana, D.J. Kundu, A. Inuganti, J. Griss, G. Mayer, M. Eisenacher, E. Pérez, J. Uszkoreit, J. Pfeuffer, T. Sachsenberg, S. Yilmaz, S. Tiwary, J. Cox, E. Audain, M. Walzer, A.F. Jarnuczak, T. Ternent, A. Brazma, and J.A. Vizcaíno. 2019. The PRIDE database and related tools and resources in 2019: improving support for quantification data. Nucleic acids research. 47:D442–D450.

Phillips, A.R., A. Suttangkakul, and R.D. Vierstra. 2008. The ATG12-conjugating enzyme ATG10 Is essential for autophagic vesicle formation in Arabidopsis thaliana. Genetics. 178:1339–1353.

Popov, N., M. Schmitt, S. Schulzeck, and H. Matthies. 1975. Reliable micromethod for determination of the protein content in tissue homogenates. Acta Biol. Med. Ger. 34:1441–1446.

Pornillos, O., D.S. Higginson, K.M. Stray, R.D. Fisher, J.E. Garrus, M. Payne, G.-P. He, H.E. Wang, S.G. Morham, and W.I. Sundquist. 2003. HIV Gag mimics the Tsg101-recruiting activity of the human Hrs protein. Journal of Cell Biology. 162:425–434. doi:10.1083/jcb.200302138.

Rabinowitz, J.D., and E. White. 2010. Autophagy and metabolism. Science. 330:1344–1348.

Rigal, A., S.M. Doyle, and S. Robert. 2015. Live cell imaging of FM4-64, a tool for tracing the endocytic pathways in Arabidopsis root cells. Methods Mol. Biol. 1242:93–103.

Rodriguez, E., J. Chevalier, J. Olsen, J. Ansbøl, V. Kapousidou, Z. Zuo, S. Svenning, C. Loefke, S. Koemeda, P.S. Drozdowskyj, J. Jez, G. Durnberger, F. Kuenzl, M. Schutzbier, K. Mechtler, E.N. Ebstrup, S. Lolle, Y. Dagdas, and M. Petersen. 2020. Autophagy mediates temporary reprogramming and dedifferentiation in plant somatic cells. EMBO J. 39. doi:10.15252/embj.2019103315.

Sanchez-Wandelmer, J., and F. Reggiori. 2013. Amphisomes: out of the autophagosome shadow? EMBO J. 32:3116–3118.

Shen, J., Q. Zhao, X. Wang, C. Gao, Y. Zhu, Y. Zeng, and L. Jiang. 2018. A plant Bro1 domain protein BRAF regulates multivesicular body biogenesis and membrane protein homeostasis. Nat. Commun. 9:3784.

Signorelli, S., Ł.P. Tarkowski, W. Van den Ende, and D.C. Bassham. 2019. Linking Autophagy to Abiotic and Biotic Stress Responses. Trends Plant Sci. 24:413–430.

Stephani, M., L. Picchianti, A. Gajic, R. Beveridge, E. Skarwan, V. Sanchez de Medina Hernandez, A. Mohseni, M. Clavel, Y. Zeng, C. Naumann, M. Matuszkiewicz, E. Turco, C. Loefke, B. Li, G. Dürnberger, M. Schutzbier, H.T. Chen, A. Abdrakhmanov, A. Savova, K.-S. Chia, A. Djamei, I. Schaffner, S. Abel, L. Jiang, K. Mechtler, F. Ikeda, S. Martens, T. Clausen, and Y. Dagdas. 2020. A cross-kingdom conserved ER-phagy receptor maintains endoplasmic reticulum homeostasis during stress. Elife. 9. doi:10.7554/eLife.58396.

Stolz, A., A. Ernst, and I. Dikic. 2014. Cargo recognition and trafficking in selective autophagy. Nat. Cell Biol. 16:495–501.

Sugano, S.S., R. Nishihama, M. Shirakawa, J. Takagi, Y. Matsuda, S. Ishida, T. Shimada, I. Hara-Nishimura, K. Osakabe, and T. Kohchi. 2018. Efficient CRISPR/Cas9-based genome editing and its application to conditional genetic analysis in Marchantia polymorpha. PLoS One. 13:e0205117.

Sutipatanasomboon, A., S. Herberth, E.G. Alwood, H. Häweker, B. Müller, M. Shahriari, A.Y. Zienert, B. Marin, S. Robatzek, G.J.K. Praefcke, K.R. Ayscough, M. Hülskamp, and S. Schellmann. 2017. Disruption of the plant-specific CFS 1 gene impairs autophagosome turnover and triggers EDS1- dependent cell death. Sci. Rep. 7:1–14.

Svenning, S., T. Lamark, K. Krause, and T. Johansen. 2011. Plant NBR1 is a selective autophagy substrate and a functional hybrid of the mammalian autophagic adapters NBR1 and p62/SQSTM1. Autophagy. 7:993– 1010.

Takemoto, K., K. Ebine, J.C. Askani, F. Krüger, Z.A. Gonzalez, E. Ito, T. Goh, K. Schumacher, A. Nakano, and T. Ueda. 2018. Distinct sets of tethering complexes, SNARE complexes, and Rab GTPases mediate membrane fusion at the vacuole in Arabidopsis. Proc. Natl. Acad. Sci. U. S. A. 115:E2457– E2466.

Tamura, K., and M. Nei. 1993. Estimation of the number of nucleotide substitutions in the control region of mitochondrial DNA in humans and chimpanzees. Mol. Biol. Evol. 10:512–526.

Thompson, A.R., J.H. Doelling, A. Suttangkakul, and R.D. Vierstra. 2005. Autophagic nutrient recycling in Arabidopsis directed by the ATG8 and ATG12 conjugation pathways. Plant Physiol. 138:2097–2110.

Wada, S., Y. Hayashida, M. Izumi, T. Kurusu, S. Hanamata, K. Kanno, S. Kojima, T. Yamaya, K. Kuchitsu, A. Makino, and H. Ishida. 2015. Autophagy supports biomass production and nitrogen use efficiency at the vegetative stage in rice. Plant Physiol. 168:60–73.

Wang, P., X. Chen, C. Goldbeck, E. Chung, and B.-H. Kang. 2017. A distinct class of vesicles derived from the trans-Golgi mediates secretion of xylogalacturonan in the root border cell. Plant J. 92:596–610.

Weidberg, H., E. Shvets, and Z. Elazar. 2011. Biogenesis and cargo selectivity of autophagosomes. Annu. Rev. Biochem. 80:125–156.

Wywial, E., and S.M. Singh. 2010. Identification and structural characterization of FYVE domain-containing proteins of Arabidopsis thaliana. BMC Plant Biol. 10:157.

Yoshii, S.R., and N. Mizushima. 2017. Monitoring and Measuring Autophagy. Int. J. Mol. Sci. 18. doi:10.3390/ijms18091865.

Zaffagnini, G., and S. Martens. 2016. Mechanisms of Selective Autophagy. J. Mol. Biol. 428:1714–1724.

Zhao, Y.G., P. Codogno, and H. Zhang. 2021. Machinery, regulation and pathophysiological implications of autophagosome maturation. Nat. Rev. Mol. Cell Biol. 22:733–750.

Zhao, Y.G., and H. Zhang. 2019. Autophagosome maturation: An epic journey from the ER to lysosomes. J. Cell Biol. 218:757–770.

Zientara-Rytter, K., and S. Subramani. 2020. Mechanistic Insights into the Role of Atg11 in Selective Autophagy. J. Mol. Biol. 432:104–122.

